# Pallatom2: Navigating the *De Novo* Protein Universe with Atom-to-Atom Modeling

**DOI:** 10.64898/2026.11.10.765209

**Authors:** Jiawei Guan, Rui Ma, Susu Lin, Yuting Wei, Jiaxin Jin, Wenjun Ye, Dechuan Liu, Haobo Wang, Weikun Wu

## Abstract

Protein binder design requires precise coordination of molecular structure, binding geometry, and experimentally measurable performance. We introduce Pallatom2, an efficient, precise Atom-to-Atom generative model that jointly models protein sequence, backbone conformation, side-chain geometry, and target-interface contacts. A unified Atom14 representation and residue-guided sparse all-atom attention enable bidirectional information exchange between residue-level and atom-level representations, coordinating global structure with local atomic detail in a compact model of just 65.9 million parameters. Its conditioning system provides fine-grained control over binder–target engagement and global and local protein conformations, supporting diverse target-specific design requirements. In silico evaluation demonstrates broad generative capability across long-chain monomers, multi-domain proteins, and homomultimers, with monomer sampling extending to 1,600 residues. In binder design, Pallatom2 achieves state-of-the-art in silico screening performance across all nine targets with cross-method benchmark data under the reported protocols. These in silico results translate into substantial experimental success: biolayer interferometry identifies 112 active designs among 210 tested candidates (53.3%), with binders obtained for all ten experimentally evaluated targets. Six targets yield picomolar-scale apparent affinities, including *K_D_* values of 3.96 pM for SC2RBD and 33.8 pM for PD-L1, while TNF*α* reaches 1.166 nM. Beyond *de novo* discovery, starting-interface-guided BCMA design preserves association kinetics despite substantial sequence divergence; partial diffusion improves TNF*α* affinity by up to 7.57-fold; and CDR-associated interface remodeling converts an inactive *de novo* nanobody into binders with apparent *K_D_* values of 4.31 and 5.93 nM. Together, these results establish Pallatom2 as a versatile model for protein design, combining efficient atomic modeling, leading in silico screening performance, and experimentally demonstrated capabilities in high-affinity binder discovery, interface transfer, affinity maturation, and recovery of inactive designs.

## 1. Introduction

Protein binders are programmable molecular recognition modules and potential carriers of biological intent from a target hypothesis to an engineered intervention. Their uses include mechanism modulation, diagnostics and delivery, affinity purification and imaging, and synthetic-biology control. Compared with antibody discovery, display-library screening, or directed evolution, *de novo* binder design can expose the intended epitope, molecular format, and candidate diversity before experimental construction, and may reach target classes that standard libraries do not systematically cover [1, 2]. However, a usable binder cannot be defined by structural appearance alone. It must form a geometrically and chemically complementary interface at the intended epitope, remain foldable, expressible, purifiable, and monodisperse, and pass experimental tests of affinity, specificity, mechanism, function, and developability. Binder design is therefore a constrained inverse-design problem whose endpoint is experimental verification rather than structure generation alone.

Recent methods address complementary parts of this problem. AlphaFold2 makes accurate protein-fold prediction broadly accessible, providing a structural foundation for assessing whether a designed sequence adopts its intended conformation [3]. AlphaFold3 extends structure prediction to complexes containing proteins and other biomolecules, supporting the assessment of molecular interactions and binding geometries [4]. RFdiffusion adapts a structure-prediction network into a diffusion-based backbone generator, enabling new folds, motif scaffolds, and target-conditioned binders beyond existing structural templates [5]. BindCraft uses AlphaFold-guided hallucination to optimize binder–target complexes, followed by sequence optimization and filtering, bringing interface design and candidate evaluation into an automated, accessible pipeline [1].

Other models focus on experimental efficiency, joint generation, and broader molecular scope. AlphaProteo generates high-affinity binders without experimental affinity maturation [6]. Chai-2 combines all-atom generation and folding-based selection for antibodies and miniproteins [7]. Latent-X jointly generates binder sequence and all-atom structure [8]. BoltzGen unifies structure prediction and design across diverse biomolecular targets [9]. SeedProteo uses an AF3-like diffusion model with self-conditioning for monomer and binder generation [10]. Proteina-Complexa couples joint sequence–structure generation with reward-guided search in a partially latent representation [2]. RFD4-Proteina (RFD4) extends generation to diverse chemistries in a shared latent space, using recurrent refinement and guided search to optimize molecular identity and conformation [11].

Together, these advances expand what can be generated and improve the prospects for experimental success. However, plausible folds and favorable in silico scores alone do not establish atomic consistency or strong experimental binding. Building on sequence-design and atomistic modeling methods [12, 13], further progress requires coordinated modeling of backbone geometry, side-chain packing, and interfacial contacts, efficient information exchange among atoms, and more reliable translation from in silico candidates to high-affinity binders.

This design problem presents the following challenges:

- **Atomic-level modeling across backbone, side chains, and binding interfaces.** Backbone geometry, side-chain packing, and interfacial contacts are tightly coupled. Modeling them jointly while preserving atomic consistency remains difficult: a plausible overall fold can still contain incompatible local geometries or unfavorable interface chemistry. The challenge is to coordinate global conformation and local atomic interactions throughout generation.
- **Efficient and accurate Atom-to-Atom information exchange.** Explicit attention among atoms imposes substantial computational and memory costs as protein size increases. The modeling challenge is equally important: information exchange must capture relevant interactions across residues and structural regions while retaining the precision needed for local atomic geometry. Achieving both tractable computation and effective atomic modeling remains difficult.
- **Reliable translation from in silico screening to high-affinity binding.** Candidates with favorable in silico scores do not consistently yield strong binding in wet-lab screening. Experimental success rates and measured affinities remain variable, leaving substantial room for improvement. In particular, in silico screening cannot reliably ensure picomolar affinity on challenging targets, making robust high-affinity binding a continuing experimental hurdle.

To address these challenges, we introduce Pallatom2, a programmable Atom-to-Atom model for protein design. It aims to translate atomic-level modeling into experimentally testable protein designs, with performance assessed through in silico benchmarks and wet-lab validation.

The main contributions of this work are:

- **A new and efficient Atom-to-Atom model for all-atom structure modeling.** Building on Pallatom1, we develop an Atom-to-Atom model with efficient all-atom attention. Residue-level relationships define sparse attention connections between atoms, enabling detailed atomic modeling with reduced computational and memory demands.
- **Unified programmable control of binding modes and generated conformations.** The model’s conditioning system jointly controls how a binder engages its target and how its global and local structure is organized. Precise control over binding contacts, interaction geometry, and generated conformations enables designs to accommodate the distinct requirements of different targets.
- **State-of-the-art in silico design performance.** Pallatom2 achieves state-of-the-art screening performance across nine benchmark targets, with high success rates for proteins such as BHRF1, PD-L1, and TrkA. It also achieves high screening success on the challenging target TNF*α* under the reported sampling and screening protocols.
- **Picomolar experimental affinity across multiple targets.** Wet-lab measurements demonstrate picomolar affinities for SC2RBD (3.96 pM), PD-L1 (33.8 pM), VEGF-A (110 pM), IL-7RA (182 pM), BCMA (187 pM), and H1 (828 pM). The challenging target TNF*α* reaches an affinity of 1.166 nM. These measurements provide experimental evidence of strong target binding beyond in silico screening performance.

The following sections present the all-atom generative model, programmable design controls, and integrated design workflow, followed by in silico evaluation and wet-lab validation.

## 2. Methods

### 2.1. All-atom generative model

Candidate generation connects design objectives and programmable constraints to the molecules selected for experimental testing. Pallatom2 performs this step through all-atom generation, jointly modeling binder conformations and their atomic contacts with the target. This allows the intended binding geometry and local interface chemistry to be considered together during candidate construction. The model thus provides structurally explicit molecular hypotheses for downstream in silico prioritization and wet-lab validation, linking design specifications to experimentally testable candidates.

#### 2.1.1. All-atom generation: rationale and process

Specifically, **Pallatom2 generates all-atom binders through iterative denoising conditioned on the target structure and specified design constraints.** These constraints include those governing binder–target engagement and binder conformation, among others, and are described in detail in subsequent sections. Starting from an initial noise state, the model progressively organizes the binder backbone and side chains within this target context, shaping their spatial arrangement and interface contacts under the specified conditions. Figure 1 illustrates the fixed-target case: a disordered atomic distribution gradually develops into an organized binder structure with defined interface contacts.

**Fig. 1.**
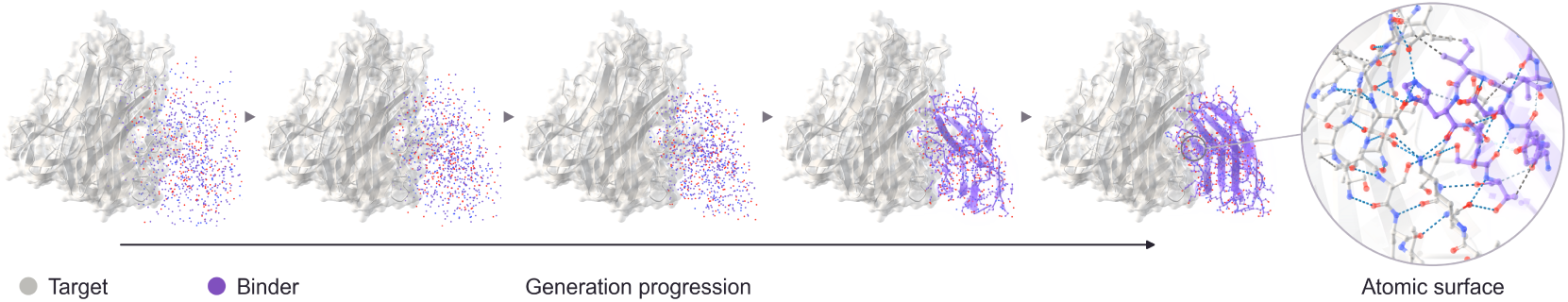
| Schematic of all-atom binder generation. With the target structure (gray) held fixed, iterative denoising progressively organizes a disordered atomic distribution into a binder conformation (purple), with backbone and side-chain geometry represented jointly. The enlarged view highlights atomic detail at the binding interface.

Within this generation process, the distinction between backbone-only and all-atom modeling is particularly important at the binding interface. Backbone-only generation primarily defines the overall fold and spatial arrangement of residues, leaving side-chain conformations and their specific atomic contacts to be resolved in subsequent steps. A plausible backbone arrangement therefore does not by itself establish favorable interface chemistry. A staged approach that fixes the backbone before sequence design also limits coordinated optimization of backbone conformation, residue identity, and side-chain arrangement.

All-atom generation, by contrast, explicitly represents both backbone and side-chain geometry, allowing their conformations and target contacts to be shaped together. Pallatom2 can thus account for interface packing, hydrogen-bond geometry, charge complementarity, and steric compatibility during candidate construction, coupling global structural organization with local chemical requirements.

#### 2.1.2. Unified Atom14 representation

Atom14 represents 39 amino acid types, encompassing L– and D-amino acids, using a fixed set of 14 atom slots per residue. Each residue is encoded as a coordinate array of shape [14, 3], with three spatial coordinates for each slot. For residues with fewer than 14 atoms, the remaining slots are padded with virtual atoms placed at the coordinates of real atoms. This representation maintains 14 atom slots per residue throughout generation, providing a consistent coordinate format across residue types. Pallatom2’s Atom14 representation distinguishes amino acids with similar conformations and resolves conformational differences between amino acids of different chirality within the same fixed-size format.

#### 2.1.3. Model architecture and generation mechanism

Building on the unified Atom14 representation, Pallatom2 directly models protein structure with an Atom-to-Atom Transformer. Atom-level single and pair representations describe individual atoms and their relationships, providing a common basis for modeling backbone geometry, side-chain conformations, and interfacial contacts throughout denoising.

##### Atom-to-Atom Transformer with a coordinate-sensitive AtomPairFormer

Pallatom2 directly models protein structure through atom-level single and pair representations: single features describe individual atoms and their structural environments as node features, while pair features encode relationships between atoms as edge features, together forming an atom-level relational graph. We construct an AtomPairFormer sensitive to three-dimensional coordinates to fuse interatomic distances, directional information, and bonding relationships into a unified atom-pair representation with 384 channels. This integration captures spatial geometry, chemical connectivity, and chirality information, supporting atom-to-atom information exchange during denoising. By representing protein structure and design conditions directly at atomic resolution, the model provides a precise basis for modeling and controlling side-chain orientations, local atomic geometry, and specified contacts throughout generation.

##### Sparse all-atom attention guided by residue-level relationships

Making atom-to-atom information exchange efficient is particularly important for large protein structures, where dense attention across all atoms must process a rapidly growing number of pairwise relationships, imposing substantial computational and memory costs. Pallatom2 addresses this burden with sparse all-atom attention guided by residue-level correlations. These correlations identify residue pairs with strong modeled relationships and define a residue relationship graph. The graph then specifies local groups of residues within which atom-level attention is computed, focusing detailed information exchange on atoms belonging to the selected residues. In this way, residue-level relationships deter-mine the connectivity of sparse all-atom attention, reducing the number of atom pairs processed while directing atomic refinement toward the interactions prioritized at the residue level.

##### Condition initialization and design constraints

Conditioning information from the target structure and specified design requirements is incorporated into the initial representations according to its granularity. For example, protein secondary-structure information is incorporated into the residue-level single representation. At the atom level, a control that fixes an amino acid side-chain orientation provides atomic structural information for initializing the atom-level pair representation. These condition-informed representations provide both a starting point and constraints for modeling the protein structural framework and refining atomic geometry. Subsequent information exchange between the two levels allows the conditions to inform both aspects of structural refinement.

##### Iterative prediction and reuse of intermediate information

The denoising network uses the current structural state and conditions to predict a refined all-atom configuration. Successive sampling steps use these predictions to update the coordinates as the noise level decreases. Within this iterative computation, recycling revisits intermediate representations, while self-conditioning supplies earlier predictions as context for subsequent updates. These mechanisms allow later predictions to build on the structural information already established during generation. The Atom14 format is retained throughout, so backbone and side-chain coordinates remain part of a common structural state across successive updates.

Together, the Atom-to-Atom Transformer, rich atom-level single and pair representations, sparse all-atom attention, and iterative refinement support direct modeling of protein structure at atomic resolution. This architecture provides the computational foundation for programmable design conditions to guide precise structural updates and atomic interactions. The next section introduces these conditions and explains how they specify binder–target engagement and control the generated protein structure.

### 2.2. Conditional control for protein design

Protein binder design is not defined solely by simply finding a plausible binding pose. A candidate must engage a biologically relevant epitope while avoiding neighboring surfaces or sites whose engagement could compromise selectivity, and it may need to accommodate local target flexibility. Some projects also call for reusing a known atomic interaction geometry or requiring selected binder residues to participate in the interface. These requirements determine which regions, conformations, and contact patterns should be explored. A target structure or hotspot list alone cannot express all of this design intent.

Making such intent explicit during generation matters because the model searches a large sequence and structure space. Without task-specific conditions, it may produce a structurally plausible complex that does not meet the biological objective. Post-generation filters can reject candidates that violate a requirement, but they do not direct sampling toward the desired contacts or conformations; later redesign can also alter the geometry that motivated a candidate. Encoding intent as generation conditions therefore links the biological hypothesis to candidate construction and makes adherence to the specification easier to assess. This improves controllability and traceability, but does not establish experimental binding or function.

Pallatom2 organizes these requirements into two complementary groups of conditions: **controls over how a binder engages its target, and controls over the protein’s own structure, sequence, and atomic geometry.** The former describe the intended recognition pattern; the latter delimit the molecular conformations available to realize it. Together, they translate a design objective into specifications that can be expressed during generation.

#### 2.2.1. Binder–target binding controls

Directing a designed binder to a target protein requires specifying how the two proteins should form the intended binding interface. **Binding control therefore concerns contacts on both the target and binder sides of the interface.** It links the target surface selected for recognition with the binder surface that engages it, guiding their relative placement and contact pattern during generation.

Pallatom2 expresses these requirements through five programmable conditions: Hotspot, Coldspot, Binderspot, Flexspot, and Unindex. Hotspot identifies target regions to engage, while Coldspot specifies regions to avoid. Binderspot identifies residues on the binder side that should participate in target engagement. Flexspot allows designated target segments to adjust locally, enabling interface design to accommodate conformational adaptation. Unindex supplies known side-chain geometry as local atomic contact information, allowing an existing interaction geometry to guide generation. **These conditions translate the intended binding mode into concrete requirements for generation.** Supplementary Table S1 summarizes the design intent and typical uses of each control; Fig. 2a illustrates their scope at the binder–target interface.

**Fig. 2.**
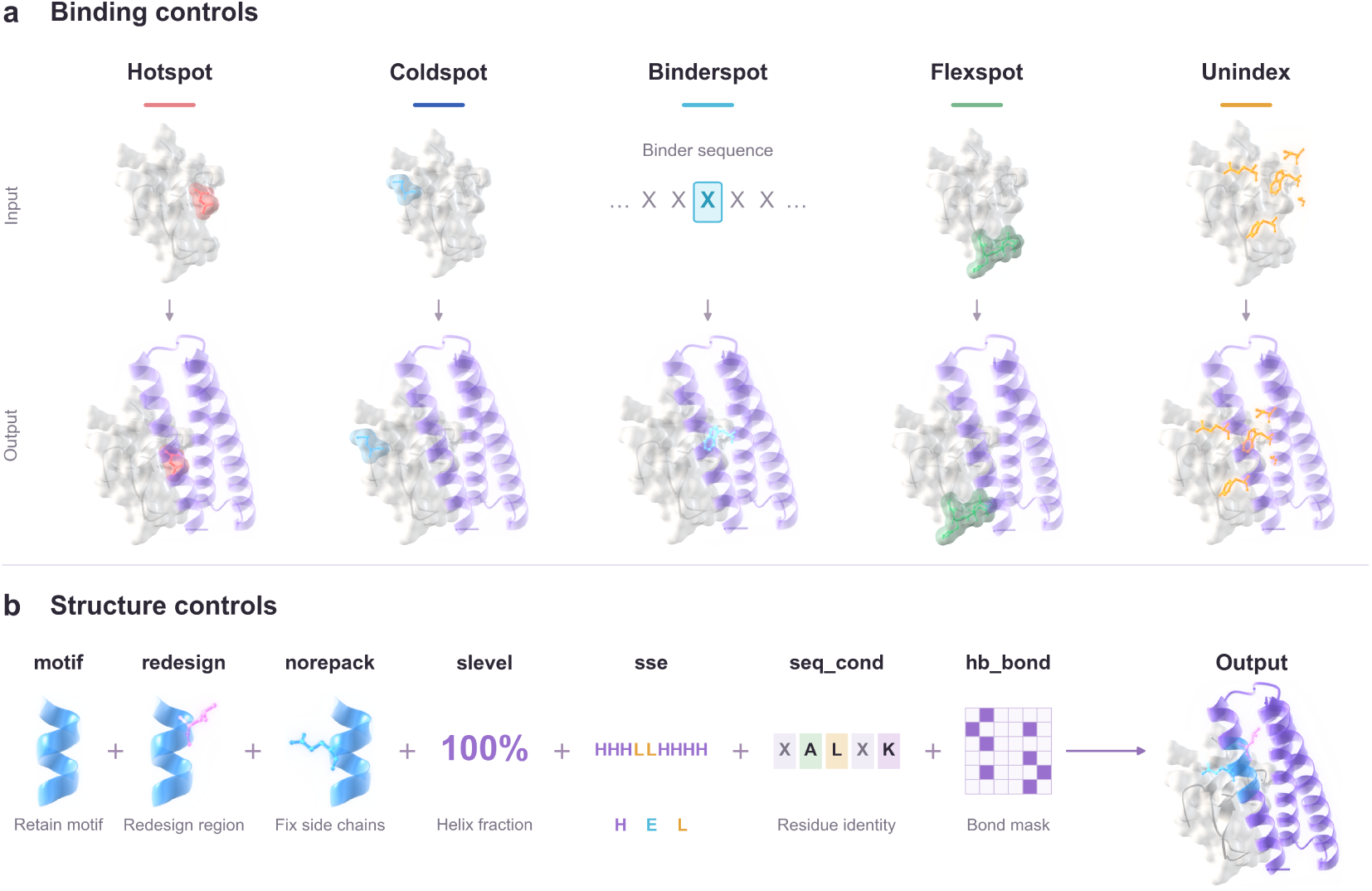
| Conditional control of binder–target binding and protein structure. **a,** Five binding controls specify target engagement and avoidance, binder-side participation, local target flexibility, and reuse of known side-chain geometry. **b,** Combined structural conditions guide protein conformation and local atomic geometry.

#### 2.2.2. Protein structure controls

Whereas binding controls specify how a binder engages its target, protein structure controls define the structural features that the generated protein should adopt or retain. The same binding objective can be realized by different protein scaffolds and local conformations, so interface requirements alone do not determine the desired protein structure. Structural controls address this additional design freedom by specifying global backbone organization, local structural features, and atomic geometry, supporting both generation under structural requirements and directed modification of an existing protein.

##### Pallatom2 organizes all-atom structural controls across three levels: backbone organization, residue identities and editable regions, and side-chain and atomic-contact geometry

At the backbone level, controls can guide the overall fold or selected local elements; for example, slevel specifies global or fragment-level secondary-structure composition, while motif specifies a structural motif to preserve. Sequence and editability controls define which regions can be redesigned and which residue identities are specified. At finer resolution, controls such as norepack preserve selected side-chain conformations, while atomic-contact conditions specify hydrogen-bond geometry. Supplementary Table S2 summarizes the individual controls and their uses.

Figure 2b illustrates how these controls can be combined within a single design task. A retained motif provides a structural anchor, redesign identifies regions available for modification, and norepack preserves selected side-chain conformations. Additional conditions specify secondary-structure composition through slevel, position-specific secondary-structure states through sse, residue identities through seq_cond, and hydrogen-bond relation-ships through an hb_bond mask. The displayed 100% helix fraction is an example input specification. Together, these conditions guide a common output structure, allowing designated regions to change while maintaining the requested global and local features.

Together, binding controls and all-atom structural controls define the design objective: the former specify how a binder engages its target, while the latter specify the global structure and local atomic geometry of the generated protein. Combining these two groups of conditions enables coordinated guidance of binder–target engagement and the protein’s own structure within the same generation process.

### 2.3. Design workflow

The all-atom model and conditional controls described above are integrated into a design workflow. Assessing Pallatom2’s practical design capability requires a systematic path from in silico generation to experimental testing. The model produces candidate structures with intended molecular properties; whether those properties are realized must be established experimentally. We therefore organize objective specification, programmable all-atom generation, candidate prioritization, and wet-lab validation into a unified workflow (Fig. 3). Computation translates target-engagement and structural requirements into candidates for testing, while experiments evaluate those candidates against the design objectives. This workflow enables systematic assessment of model outputs and uses experimental evidence to guide subsequent design, forming a cycle of **in silico design, experimental validation, and evidence-guided feedback**.

**Fig. 3.**
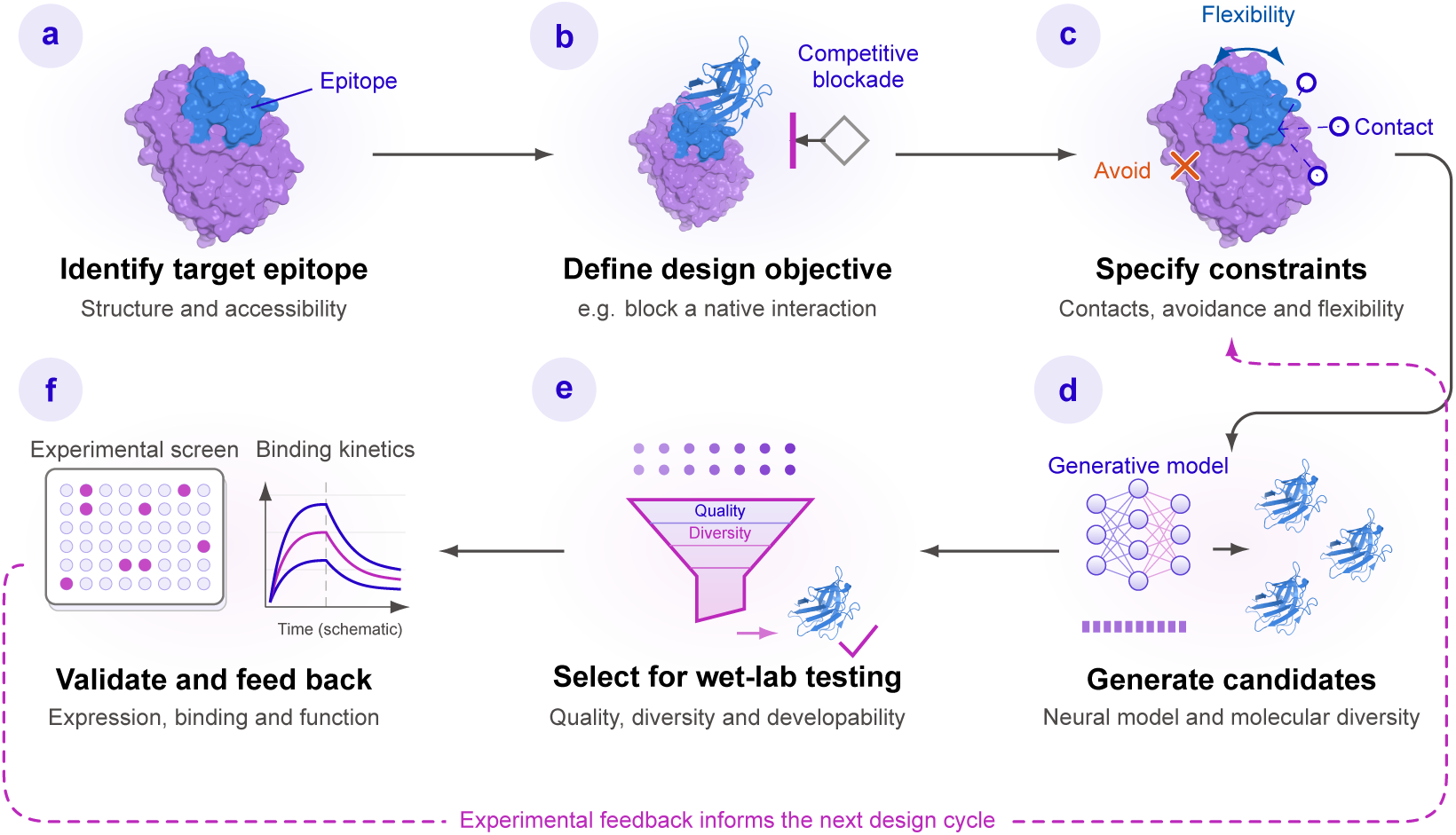
| Protein design workflow. (a) Identify the target epitope from the target structure and its accessibility. (b) Define the design objective, for example blocking a native interaction. (c) Encode executable constraints, including target regions to engage or avoid and regions permitted to undergo local conformational changes. (d) Generate all-atom binder–target candidate complexes under these constraints. (e) Select a panel for wet-lab validation on the basis of structural quality, diversity, and developability. (f) Assess expression, binding, affinity, specificity, and function through wet-lab validation against design objectives, and use the results to guide the next design round.

### Wet-lab validation provides a critical test of model outputs

Expression, binding, affinity, specificity, functional, and structural measurements provide empirical evidence of whether in silico designs translate into the intended molecular behavior. This evidence is essential for assessing the model’s practical design capability: predicted structures and favorable in silico scores alone cannot establish experimental performance. Outcomes from tested candidates also help identify where in silico expectations agree with measurements and where further investigation or design refinement is needed.

(a) **Identify the target epitope.** Begin with the target structure and identify an accessible epitope. This establishes the target region that the binder is intended to recognize.
(b) **Define the design objective.** Specify the intended biological outcome, for example blocking a native interaction. The objective provides the basis for formulating the molecular hypothesis and assessing whether a candidate meets the design intent.
(c) **Encode executable constraints.** Translate the objective into programmable conditions, including required contacts, regions to avoid, and local flexibility. These conditions specify how candidates should engage the target during generation.
(d) **Generate all-atom candidates.** Explore sequence and structure space under the specified constraints to produce all-atom binder–target complexes. The generation model translates target specifications and design intent into molecular candidates for subsequent evaluation.
(e) **Prioritize candidates for testing.** Select a panel on the basis of structural plausibility, interface quality, diversity, developability, and experimental tractability. This stage determines which in silico hypotheses advance to wet-lab validation.
(f) **Validate experimentally.** Evaluate expression, binding, affinity, specificity, function, and structure through staged wet-lab testing. Relate the measurements to the in silico design objectives to determine whether selected model outputs meet the intended criteria under the conditions tested. Each assay contributes evidence for the property it measures.

#### Feedback into the next design round

Experimental outcomes, including designs that do not meet the intended criteria, inform revisions to target selection, design constraints, and candidate prioritization. Computation supplies molecular candidates and testable expectations; wet-lab measurements assess those expectations and provide evidence for subsequent design decisions. This feedback connects generation, evaluation, and refinement within the same workflow.

This workflow connects the generation and control capabilities described above to in silico evaluation and experimental validation. We next assess the generated candidates through in silico benchmarks, followed by wet-lab testing.

## 3. In silico Benchmarks

The preceding section introduced Pallatom2’s all-atom generation mechanism, programmable design controls, and integrated design workflow. Here, in silico benchmarks assess two aspects of the model’s generative capability. Unconditional protein monomer sampling assesses structural plausibility and diversity across protein lengths and domain architectures. Conditional protein binder design assesses structural consistency and interface quality under target and epitope conditions. The following sections present these benchmarks and extend the binder analysis to D-form designs against L-form targets.

### 3.1. Unconditional all-atom protein structure generation

Unconditional sampling removes the target and directly evaluates the learned generative prior. We examine long-chain monomers, multi-domain monomers, and homomultimers to assess sequence–structure consistency across chain lengths and structural organizations, together with sampling diversity for the monomer benchmarks.

Sequence–structure consistency is evaluated by refolding the designed sequences with AlphaFold3 and comparing the predicted structures with the generated structures; a sample passes when its **all-atom RMSD is *≤* 2.0 Å** and its **mean pLDDT is *≥* 80.0**. Designability is the fraction of evaluated samples that pass both criteria. Structural diversity is assessed by clustering passing designs at a TM-score threshold of 0.5. The cluster rate is the number of structural clusters among passing designs divided by the total number of samples evaluated, multiplied by 100%; it therefore measures the yield of structurally distinct, passing designs.

#### Long-chain monomers

A monomer consists of a single protein chain. Long-chain sampling tests the model’s ability to generate coherent full-length folds, with tested lengths extending beyond the training range to explore generalization. As chain length increases, the model must coordinate long-range residue interactions while maintaining consistency between the designed sequence and the overall fold. Evaluation focuses on sequence–structure consistency and structural diversity across length bins, with particular attention to how performance changes be-yond the training range. Figure 4 summarizes the monomer benchmarks, including length-dependent sampling, secondary-structure classes, and multi-domain organization.

**Fig. 4.**
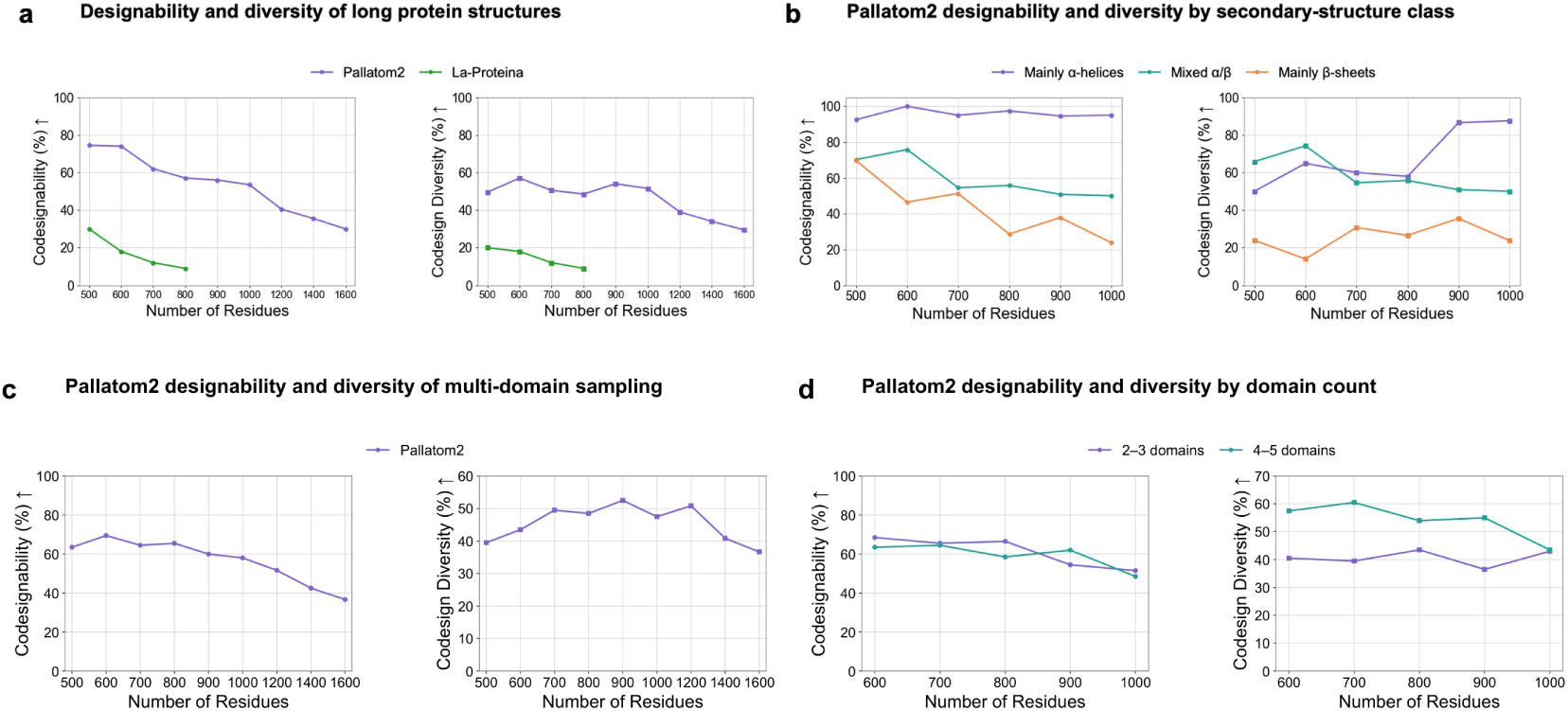
| Designability and structural diversity across protein lengths and sampling conditions. Each panel shows designability (left) and diversity (right). (a) Long-protein sampling over 500–1600 residues, with La-Proteina reference points at 500–800 residues. (b) Pallatom2 sampling by secondary-structure class: Mainly *α*-helices, Mixed *α*/*β*, and Mainly *β*-sheets. (c) Pallatom2 multi-domain sampling over 500–1600 residues. (d) Sampling with 2–3 versus 4–5 domains. Panel a uses 200 total designs per length; panel b shows 500–1000 residues, with rates computed within each secondary-structure class. Diversity is successful cluster count divided by the corresponding total sample count. Panels c and d retain the previous multi-domain results. Tested length bins are equally spaced.

#### Length-dependent sampling results

Over 500–1600 residues, Pallatom2 designability decreases from 74.5% at 500 residues to 53.5% at 1000 residues and 30.0% at 1600 residues. The cluster rate ranges from 29.5% to 57.0%, peaking at 600 residues (Figure 4a). Both rates use 200 generated samples per length as the denominator. La-Proteina reference points are shown over 500–800 residues; these are descriptive comparisons rather than a matched-protocol evaluation.

#### Secondary-structure dependence

In the updated sampling batch, Mainly *α*-helices designs achieve 92.5–100% designability over 500–1000 residues, reaching 100% at 600 residues. Mixed *α*/*β* designs range from 50.0% to 75.8%, while Mainly *β*-sheets designs range from 23.8% to 69.6%. At 1000 residues, the within-class cluster rates are 87.5%, 50.0%, and 23.8%, respectively (Figure 4b). These class-specific rates use each class’s sample count as the denominator, whereas panel a summarizes all samples at each length.

To examine sequence–structure consistency at longer chain lengths, Figure 5 presents six selected monomer designs spanning 500–1000 residues. These examples complement the aggregate sampling evaluation by showing the generated folds and their agreement with structures repredicted from the designed sequences.

**Fig. 5.**
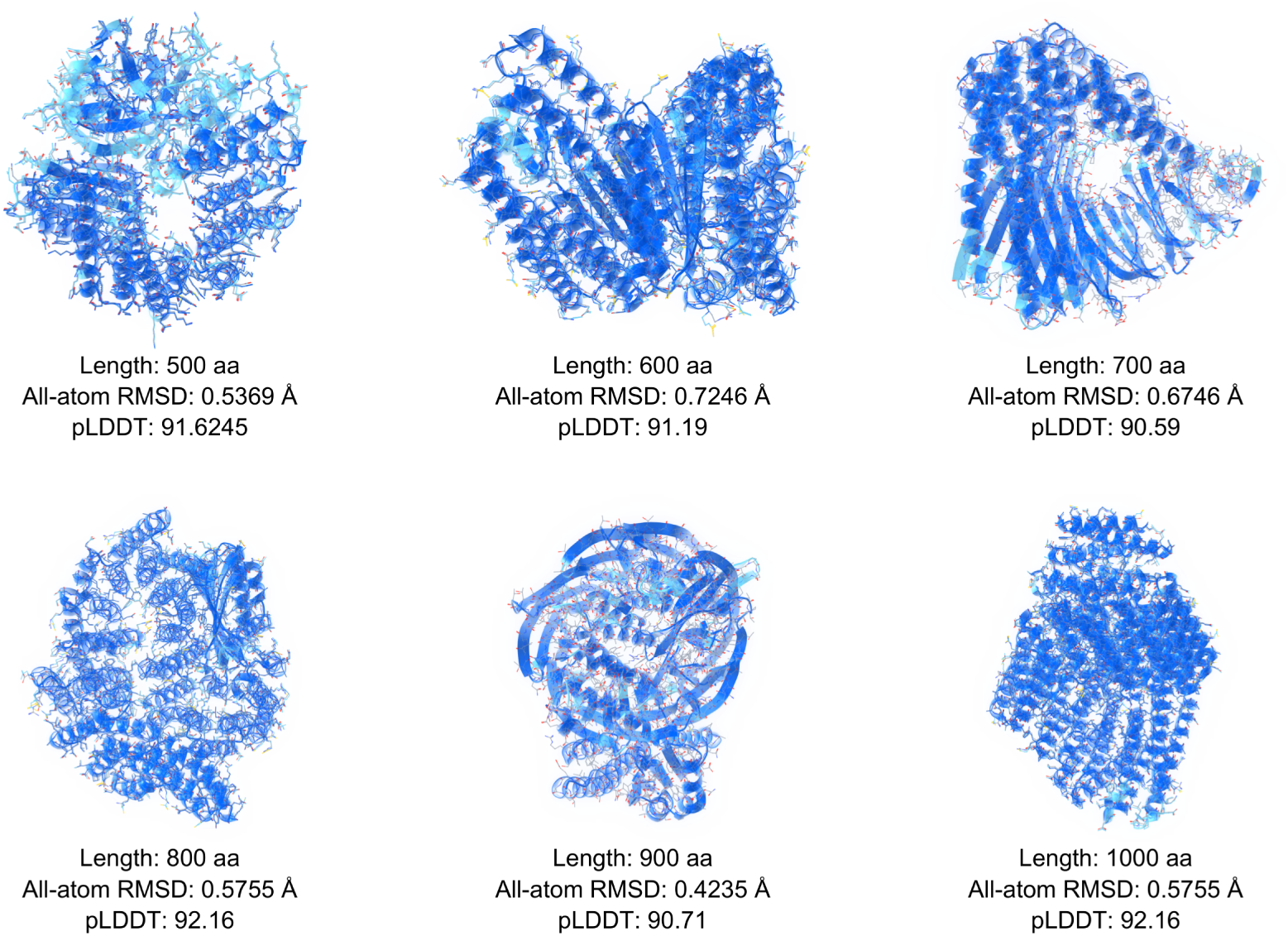
| Structural examples from unconditional protein monomer sampling. From left to right and top to bottom, protein lengths are 500, 600, 700, 800, 900, and 1000 residues. Each structure is annotated with its length, all-atom RMSD, and mean pLDDT. These metrics provide evidence of consistency between sequence and structure.

The six examples have all-atom RMSD values of 0.4235–0.7246 Å and mean pLDDT values of 90.59–92.16. The 1000-residue example retains an all-atom RMSD of 0.5755 Å and a pLDDT of 92.16, supporting sequence–structure consistency in this long-chain design. These selected cases demonstrate attainable sequence–structure consistency across the displayed lengths; length-dependent sampling success is assessed separately by the aggregate results in Figure 4.

#### Multi-domain monomers

Beyond chain length, domain organization adds another level of structural complexity. A multi-domain monomer contains multiple distinct folded units within one protein chain. Analysis of our dataset shows that longer proteins tend to contain multiple domains, while single-domain monomers longer than 500 residues are rare. This is consistent with structural databases, in which long proteins commonly combine smaller domains and most domains are shorter than 300 residues, a typical size range rather than a strict upper bound [14]. Multi-domain sampling therefore tests whether the model can coordinate the folding of individual domains with their relative orientations, inter-domain packing, and connecting regions to form a coherent full-length structure.

#### Multi-domain sampling and domain-count dependence

Across the same 500–1000-residue range, multidomain sampling shows more stable designability with increasing length than the long-chain monomer sampling results in Figure 4a. Multi-domain designability starts at 63.5% at 500 residues, reaches 69.5% at 600 residues, and then decreases overall to 58.0% at 1000 residues, spanning a narrower range than the corresponding long-chain monomer results (53.5–74.5%). Over this interval, the multi-domain cluster rate increases from 39.5% to 47.5%, with a peak of 52.5% at 900 residues (Figure 4c). Panels c and d retain the earlier multi-domain sampling batch. Across 600–1000 residues, the 4–5-domain group has higher cluster rates than the 2–3-domain group (Figure 4d). Comparisons with the updated panels a and b should account for differences in sampling batches and denominators.

Figure 6 complements these sampling statistics with nine selected designs containing two to six domains and spanning 500–1600 residues, illustrating sequence–structure consistency across both chain length and domain organization.

**Fig. 6.**
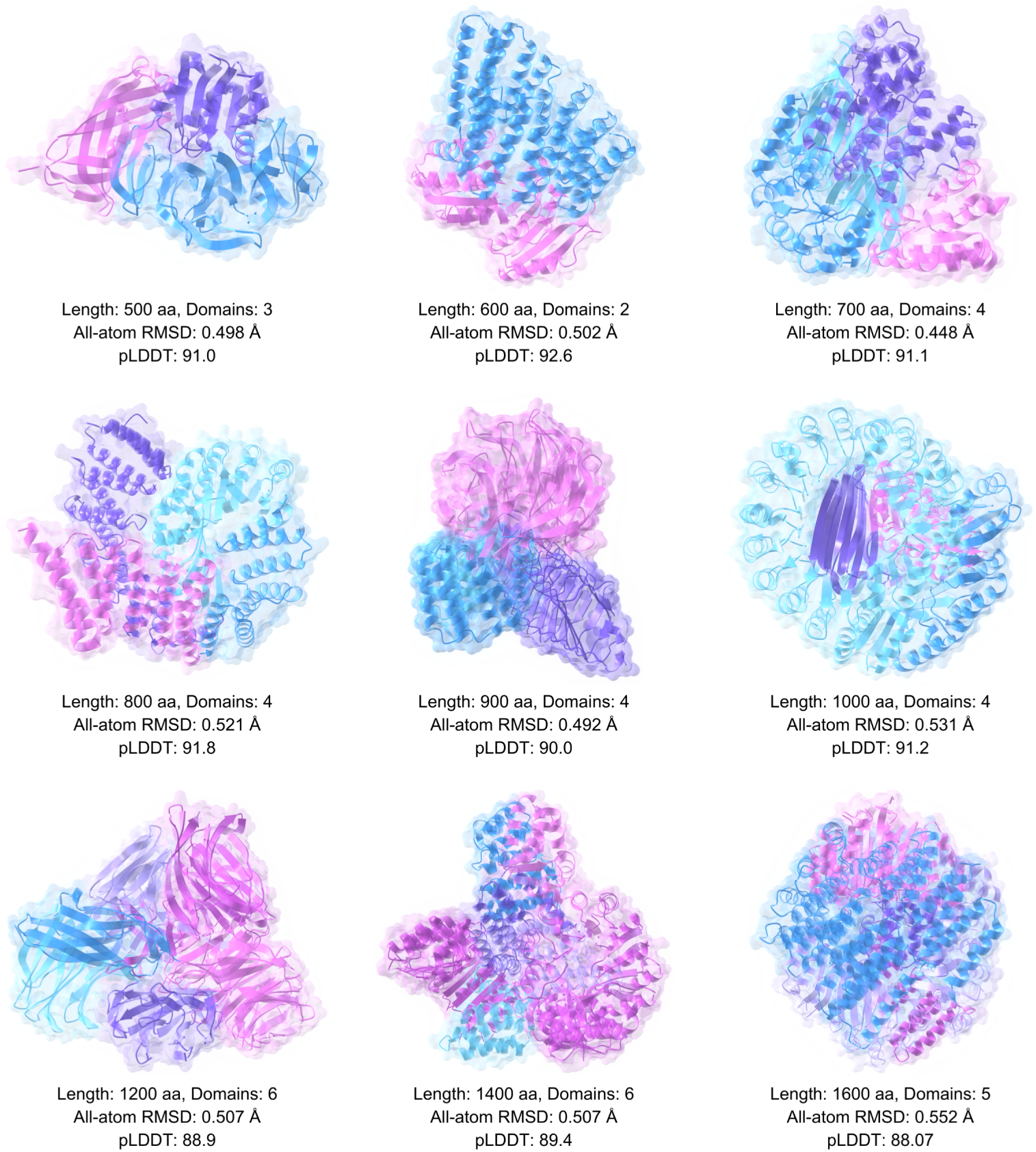
| Multi-domain protein sampling examples. From left to right and top to bottom, protein lengths are 500, 600, 700, 800, 900, 1000, 1200, 1400, and 1600 residues. Low all-atom RMSD and high pLDDT upon sequence-based structure reprediction support consistency between the designed sequences and generated structures.

The multi-domain examples have all-atom RMSD values of 0.448–0.552 Å and pLDDT values of 88.07–92.6; the 1200–1600-residue designs have all-atom RMSD values of 0.507–0.552 Å, extending sequence–structure consistency beyond the 1000-residue range. Their low full-chain RMSD supports consistency of the overall domain arrangement as well as the constituent folds, while the high pLDDT provides complementary confidence in local structure. These selected examples illustrate attainable sequence–structure consistency across chain lengths and domain architectures, while the long-chain monomer results in Figure 4 provide a length-dependent reference for sampling success and diversity. Domain-resolved alignment and inter-domain contact analysis remain necessary to distinguish local folding accuracy from errors in relative domain placement.

#### Homomultimers

Extending from organization within a single chain to assembly across chains, homomultimer sampling examines complexes of multiple protein chains with the same sequence. The model must generate well-folded subunits while forming compatible inter-chain interfaces and achieving the intended subunit number, relative arrangement, and symmetry. Figure 7 presents nine selected designs with C3, C4, C5, C6, C8, and C10 symmetry, spanning reported assembly lengths of 600–2000 residues.

**Fig. 7.**
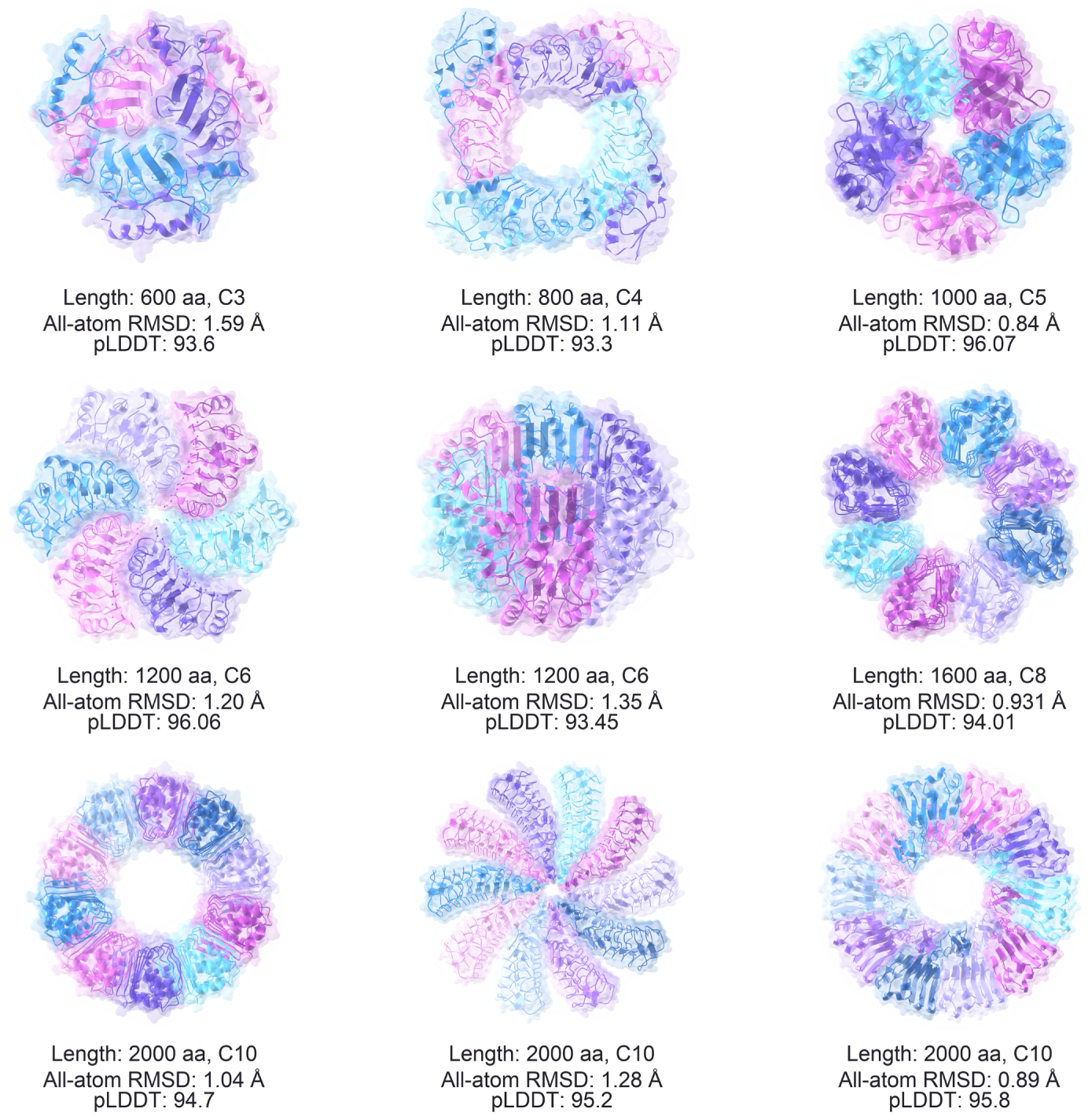
| Structural examples from homomultimer sampling. From left to right, top to bottom: 600 aa/C3, 800 aa/C4, 1000 aa/C5, two 1200 aa/C6 designs, 1600 aa/C8, and three 2000 aa/C10 designs. Panels are independently scaled. All-atom RMSD and pLDDT are annotated for each design.

Across the nine homomultimer examples, all-atom RMSD ranges from 0.84 to 1.59 Å and mean pLDDT ranges from 93.3 to 96.07. The C5 design has an all-atom RMSD of 0.84 Å and pLDDT of 96.07; the two C6 designs have RMSD values of 1.20 and 1.35 Å and pLDDT values of 96.06 and 93.45, respectively. The 1600-residue C8 design has an all-atom RMSD of 0.931 Å and pLDDT of 94.01. The three C10 designs, at 2000 reported residues each, have all-atom RMSD values of 0.89–1.28 Å and pLDDT values of 94.7–95.8. Together with the C3 and C4 examples, these results support sequence–complex structure consistency across the displayed assembly sizes and symmetries. They extend the structural examples from single-chain folds and multi-domain organization to assemblies of identical subunits; an aggregate homomultimer sampling success rate is not estimated from these selected cases.

### 3.2. Conditional protein binder design

Protein binder design starts from a target protein structure and a specified binding epitope. Conditioned on these inputs, a generative model produces a protein structure and its associated sequence intended to bind the designated site on the target. This task extends the generation of an independently folded protein to the coordinated design of the binder’s own fold and its interaction with the target.

#### The central challenge is to satisfy folding and interface requirements simultaneously

The binder must adopt a coherent structure, approach the specified epitope in an appropriate orientation, and form geometrically and chemically compatible contacts with the target. Sequence identity, backbone conformation, and side-chain arrangement must therefore be coordinated across both the binder fold and the binding interface. Differences in epitope shape, surface polarity, accessibility, and target assembly further change the design requirements across targets.

#### In silico evaluation

We screen Pallatom2 candidates using three criteria: **allatom_rmsd *≤* 2.5 Å**, **min-ipAE *≤* 1.5**, and **ptm-binder *≥* 0.8**. A candidate must satisfy all three thresholds to pass the in silico screen. These metrics assess agreement between the generated and repredicted structures, inter-chain alignment uncertainty, and predicted binder fold confidence, respectively.

#### Target diversity

The evaluation panel comprises ten targets spanning viral antigens, an immune checkpoint ligand, cytokines and their receptors, a growth factor, a TNF receptor superfamily member, a receptor tyrosine kinase, and a viral anti-apoptotic protein. Beyond their biological functions, the targets vary in surface geometry, polarity, size, and oligomeric organization, encompassing small globular domains, flat or polar surfaces, binding grooves, and multimeric structures. This diversity tests whether binder generation remains effective across distinct interface environments. Table 1 summarizes the target classes and biological roles, and Figure 8 shows their representative structures.

**Fig. 8.**
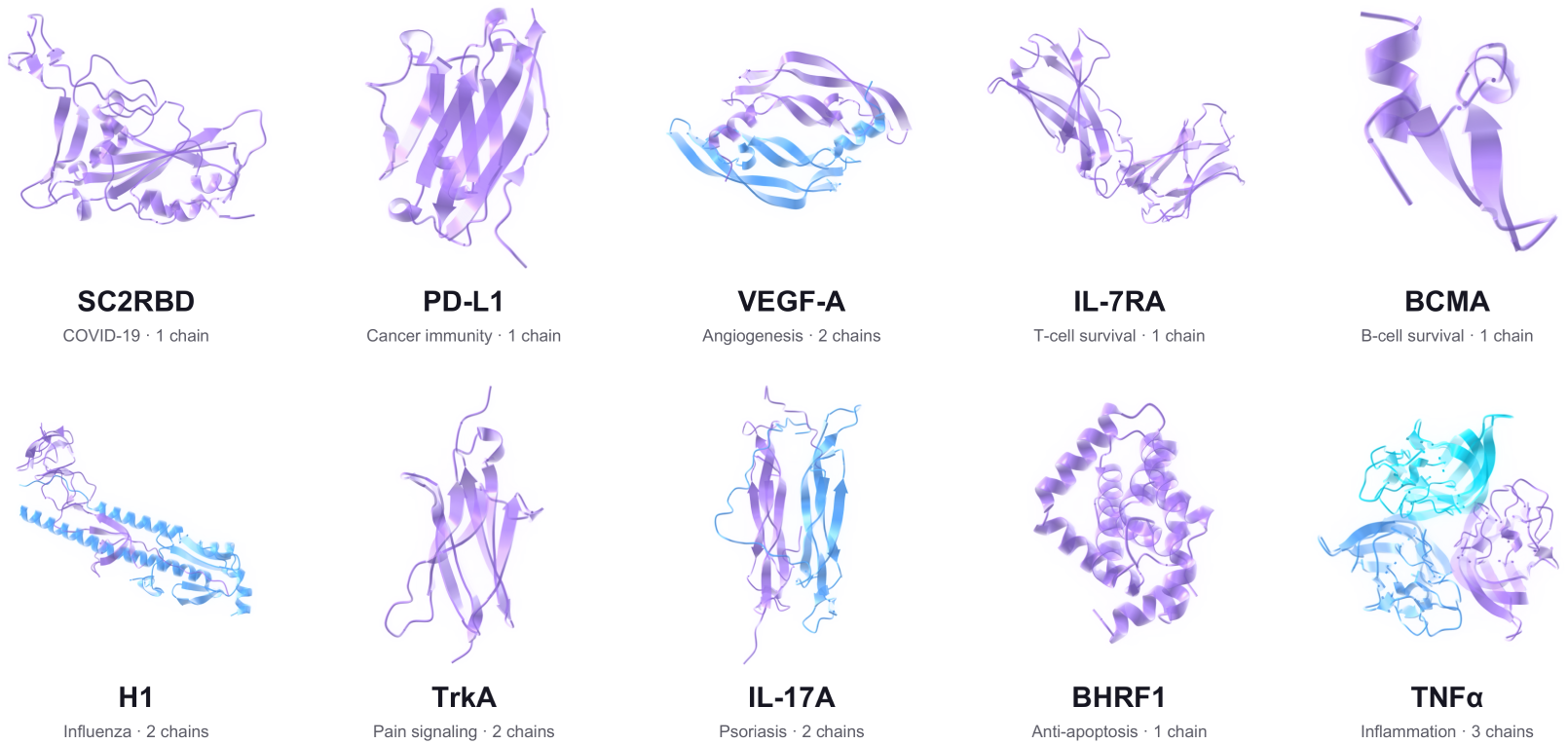
| Representative structure images for the ten benchmark targets, ordered as in Table 1. Each panel is independently scaled.

**Tab. 1.**
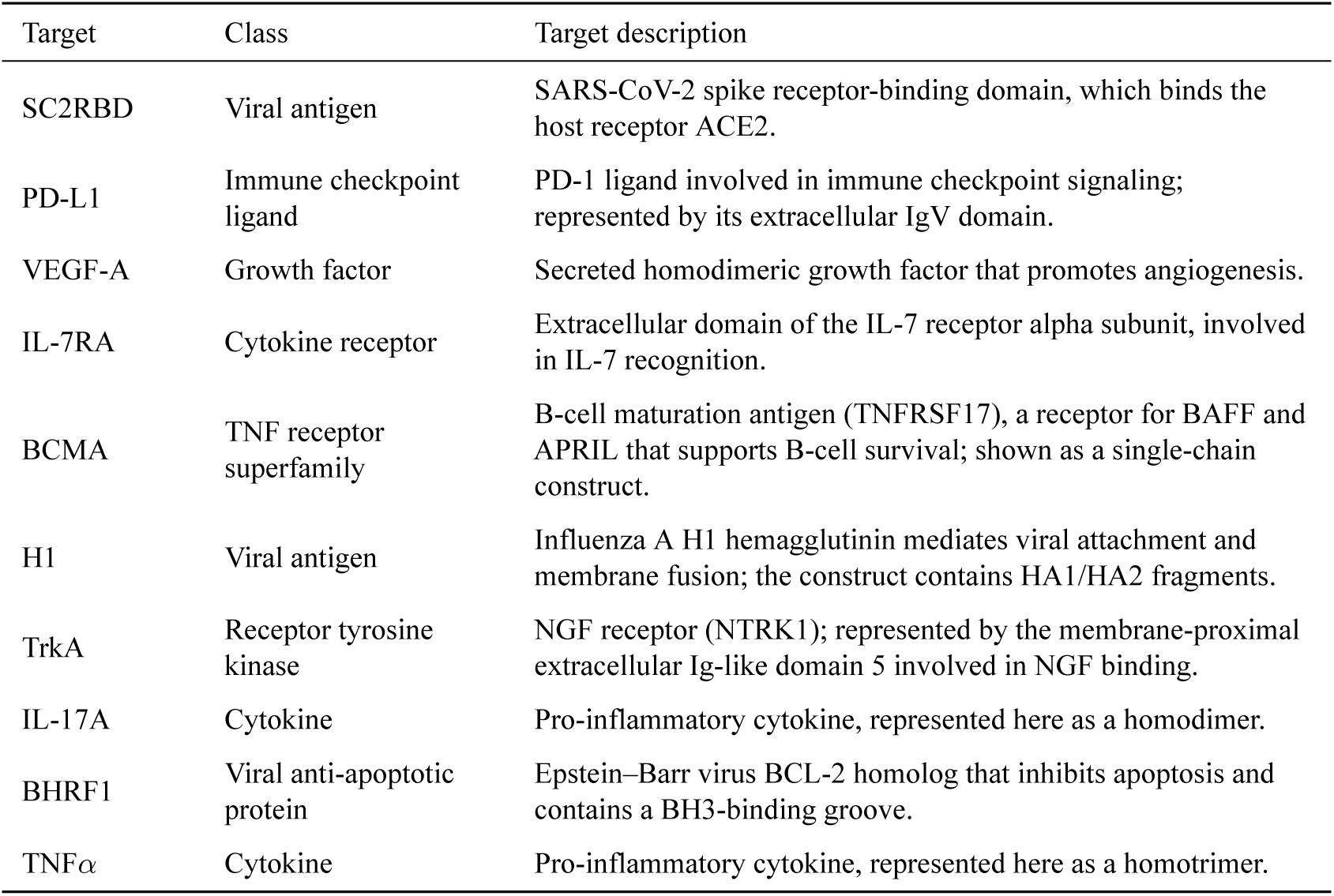
| Ten-target conditional binder benchmark panel.

#### Sampling and screening

Pallatom2 search explores and evaluates alternative branches of the generation trajectory to identify high-quality all-atom structures. This approach is conceptually similar to RFD4 search [11]. For each target–epitope case, Pallatom2 generates candidates in each of the Mainly *α*-helices, Mixed *α*/*β*, and Mainly *β*-sheets secondary-structure classes; the realized per-target sampling counts are summarized in Supplementary Fig. S5b. Candidates are screened against the three in silico criteria above. Sampling across these classes explores alternative binder folds for the specified target site.

#### Leading screening success across targets

Figure 9a compares the three Pallatom2 secondary-structure classes with RFD4 search, BindCraft, and BoltzGen. Mainly *α*-helices sampling achieves the highest reported screening success rate in all nine cases with cross-method data, ranging from 35.6% for TNF*α* to 98.5% for BHRF1. Its success rates reach 94.9% for PD-L1 and 93.8% for TrkA, while TNF*α* remains more challenging for all four methods. These results show a consistent lead across the displayed target panel. They describe in silico screening outcomes under the respective sampling protocols, rather than experimental binding hit rates or a comparison at matched computational cost.

**Fig. 9.**
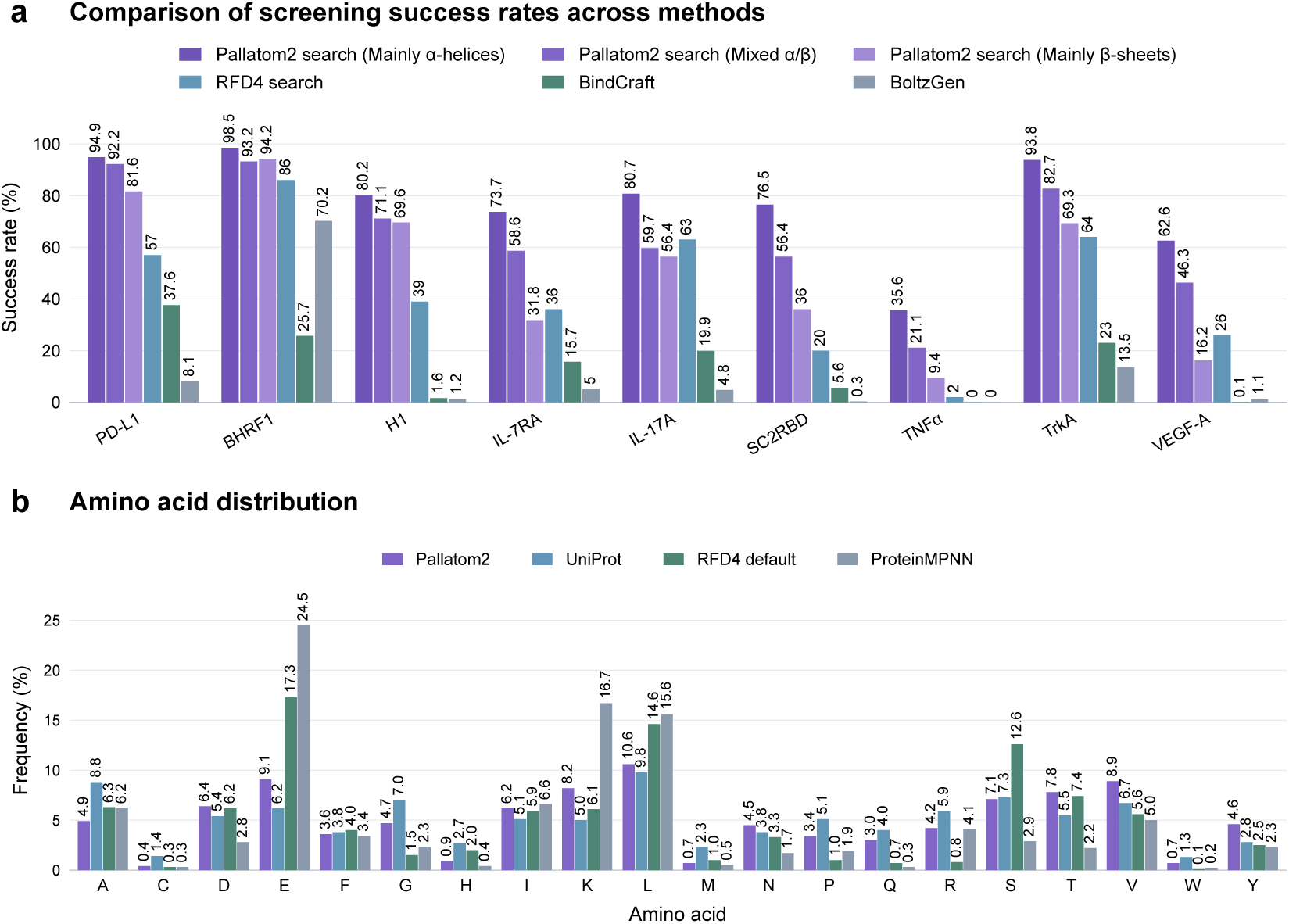
| In silico binder-design outcomes. (a) Screening success rates across nine targets for Pallatom2 search (Mainly *α*-helices), Pallatom2 search (Mixed *α*/*β*), and Pallatom2 search (Mainly *β*-sheets) (dark to light purple), alongside RFD4 search, BindCraft, and BoltzGen. BCMA is absent from this comparison. (b) Amino acid frequencies for Pallatom2 (17,990,544 binder residues), UniProt, RFD4 default sampling (*k* = (16, 4)), and ProteinMPNN redesigns of default RFD4 backbones. The latter two references are from unconditional generation. The success-rate sampling protocol uses 400 candidates per secondary-structure class per target–epitope case.

#### Secondary structure influences screening success

Across the nine cases in Figure 9a, Mainly *α*-helices sampling consistently yields the highest success rate. The Mixed *α*/*β* class generally performs between the Mainly *α*-helices and Mainly *β*-sheets classes, with BHRF1 as an exception. The magnitude of this effect depends on the target: Mainly *α*-helices and Mainly *β*-sheets achieve 73.7% and 31.8% for IL-7RA, compared with 98.5% and 94.2% for BHRF1. Thus, secondary-structure composition is an effective sampling control, with helix-rich designs favored in this benchmark and the benefit varying across target interfaces.

#### Natural-like composition with high designability

Figure 9b shows that Pallatom2 binder sequences have an overall amino acid distribution closer to the UniProt reference than the displayed RFD4 default and ProteinMPNN distributions. For example, glutamate and leucine are less enriched than in either comparator, although residue-specific differences from UniProt remain. RFD4-Proteina reports that sampling temperature changes sequence composition and provides control over the tradeoff between designability and nativeness; its default distribution remains closer to UniProt than to ProteinMPNN redesigns [11]. Here, the composition analysis and screening results together show that Pallatom2 combines a relatively natural-like amino acid distribution with high in silico designability. This coexistence indicates that high screening success does not require reproducing the strongly skewed ProteinMPNN composition shown here; it does not establish amino acid composition as the cause of designability.

Together, these in silico analyses assess screening success, its dependence on secondary structure, and sequence composition. Additional analyses of screening metrics and interface geometry are provided in Supplementary Section S3.2.

### 3.3. Design of D-form binders

We extended target-conditioned generation to D-form binders against four L-form protein targets: IL-7RA, IL-17A, VEGF-A and PD-L1 (Fig. 10). To assess backbone consistency, we first used AlphaFold3 (AF3) [4] to predict the L-form counterpart of each designed binder. We then manually inverted the predicted coordinates to obtain the corresponding D-form structure and superimposed its backbone onto that of the binder in the sampled D-binder–L-target complex.

**Fig. 10.**
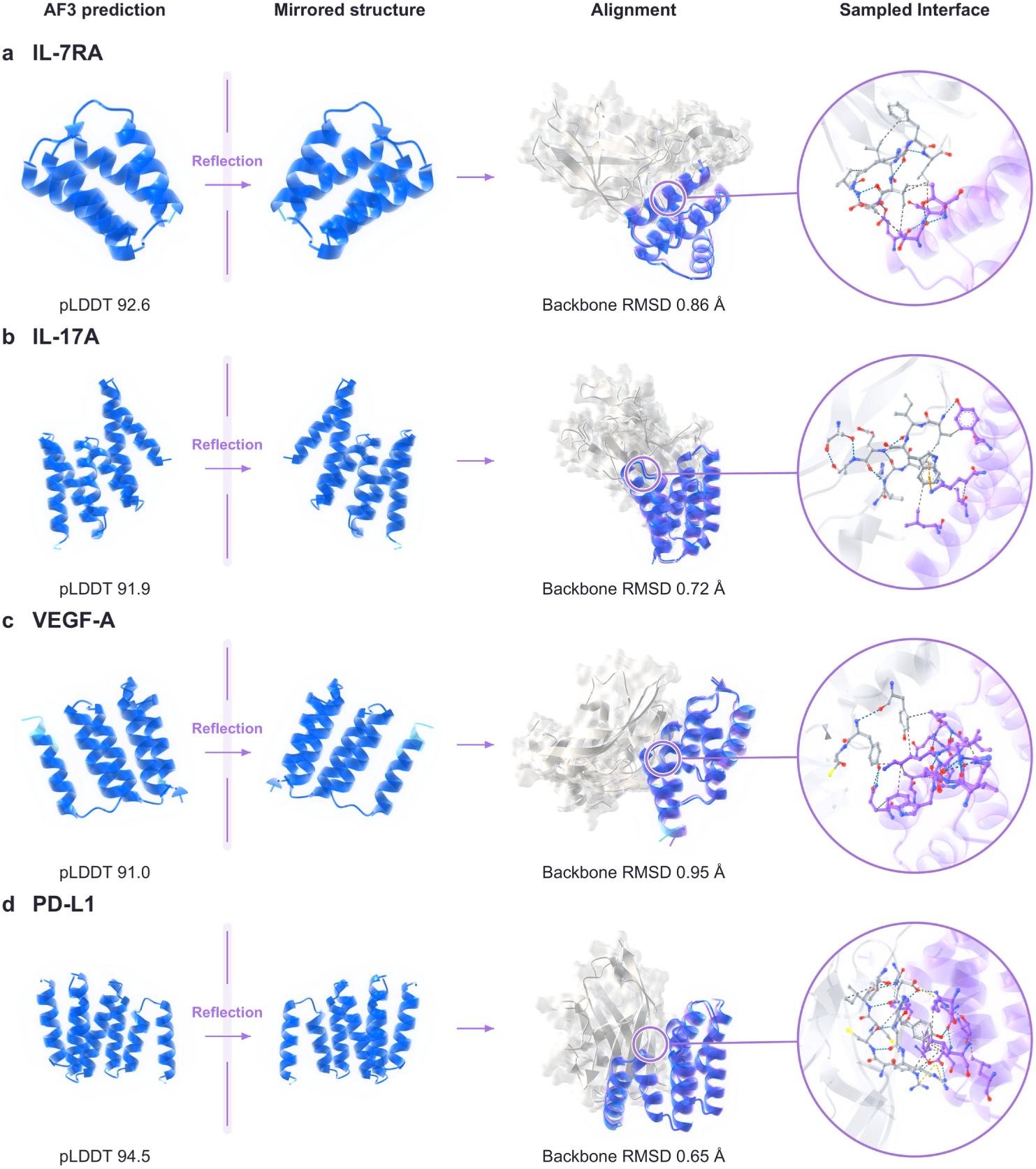
| D-form binder designs against L-form protein targets. Panels a–d show IL-7RA, IL-17A, VEGF-A and PD-L1, respectively. Each row presents the AF3-predicted L-form counterpart, the D-form structure obtained by manual coordinate inversion, its backbone alignment with the sampled D-binder in the L-target complex, and a close-up of the sampled interface. AF3 pLDDT scores are 92.6, 91.9, 91.0 and 94.5; the corresponding backbone RMSDs are 0.86, 0.72, 0.95 and 0.65 Å. pLDDT refers to the L-form predictions, and RMSD compares their mirror-transformed structures with the sampled D-binders.

Across the four examples, the AF3 predictions of the L-form counterparts had pLDDT scores of 91.0–94.5. After coordinate inversion and backbone superposition, the RMSDs relative to the sampled D-binders were 0.65–0.95 Å. These results show close agreement between the mirror-transformed predicted folds and the binder backbone conformations in the sampled complexes, supporting structural consistency across the four D-form designs.

## 4. In vitro experiments

Wet-lab experiments provide an independent test of the in silico results. Structural consistency, prediction confidence and favourable interface features support candidate selection, but they do not establish that a designed protein can be produced or bind its intended target. We therefore evaluated expression and measured binding kinetics across targets by biolayer interferometry (BLI), then used the resulting evidence to refine starting interfaces, affinity and molecular format.

### In silico screening

Candidates for wet-lab testing were selected using the in silico criteria introduced in Section 3.2: all-atom RMSD *≤* 2.5 Å, minimum inter-chain PAE *≤* 1.5, and binder pTM *≥* 0.8. These criteria assess sequence–structure consistency and confidence in the designed complex. Sequence and structural diversity were also considered to reduce redundant experimental testing.

### Agent-assisted candidate analysis

In addition to these metrics, we used Lévin™ Harness^1^ to analyze candidate structures and their evaluation results and support selection for wet-lab testing (Fig. 11). This contextual review complemented numerical screening and informed candidate prioritization; experimental measurements then provided independent validation of the selected designs.

**Fig. 11.**
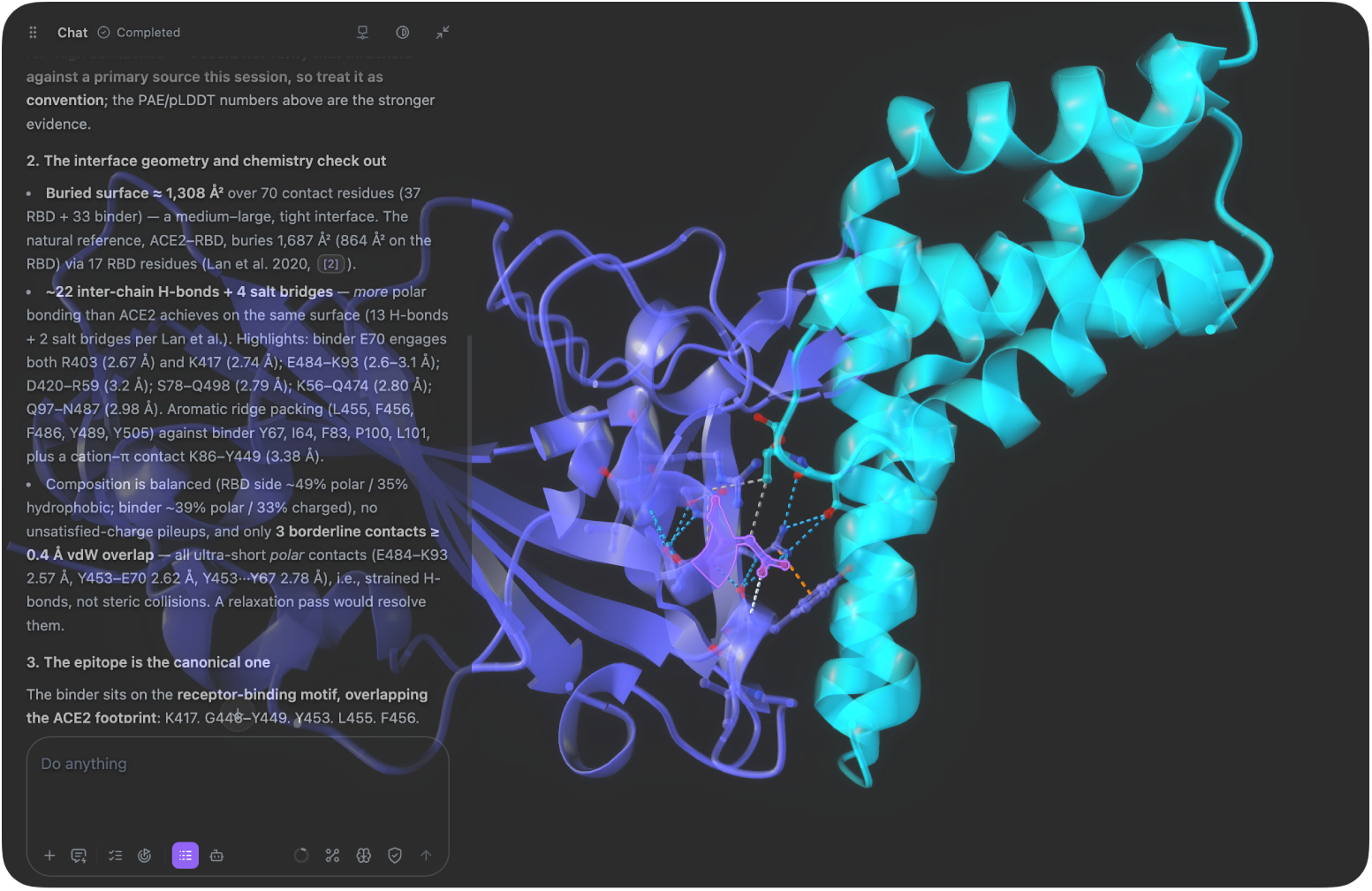
| Agent-assisted candidate analysis with Lévin™ Harness. A three-dimensional view of a binder–target complex is displayed alongside an assessment of interface geometry and chemistry, supporting candidate review and prioritization for experimental testing.

### 4.1. An experimental screening landscape across ten targets

We designed and experimentally screened minibinders against BCMA, PD-L1, VEGF-A, SC2RBD, IL-7RA, BHRF1, TNF*α*, TrkA, IL-17A and H1. After in silico filtering, ranking and manual nomination, we completed 210 BLI experiments, covering 12–24 candidates per target. Of these, 112 were active and 98 were inactive, giving an overall active fraction of 53.3% (Fig. 12d; Supplementary Fig. S5a). Each target also included one reference binder: AlphaProteo designs for PD-L1, IL-7RA, VEGF-A, IL-17A and TrkA [6]; BC40 for BCMA [15]; Latent-X designs for SC2RBD [8]; an ozoralizumab-derived nanobody for TNF*α* [16]; and binders from early landmark in silico design studies for H1 and BHRF1 [17, 18]. Reference binders confirmed target loading and assay performance and provided a benchmark for comparing measured apparent *K_D_* with the original reports. Available sequences and measured affinities of active minibinders and reference binders, together with the reported affinities and assay sources for the reference binders, are compiled in Supplementary Table S5.

**Fig. 12.**
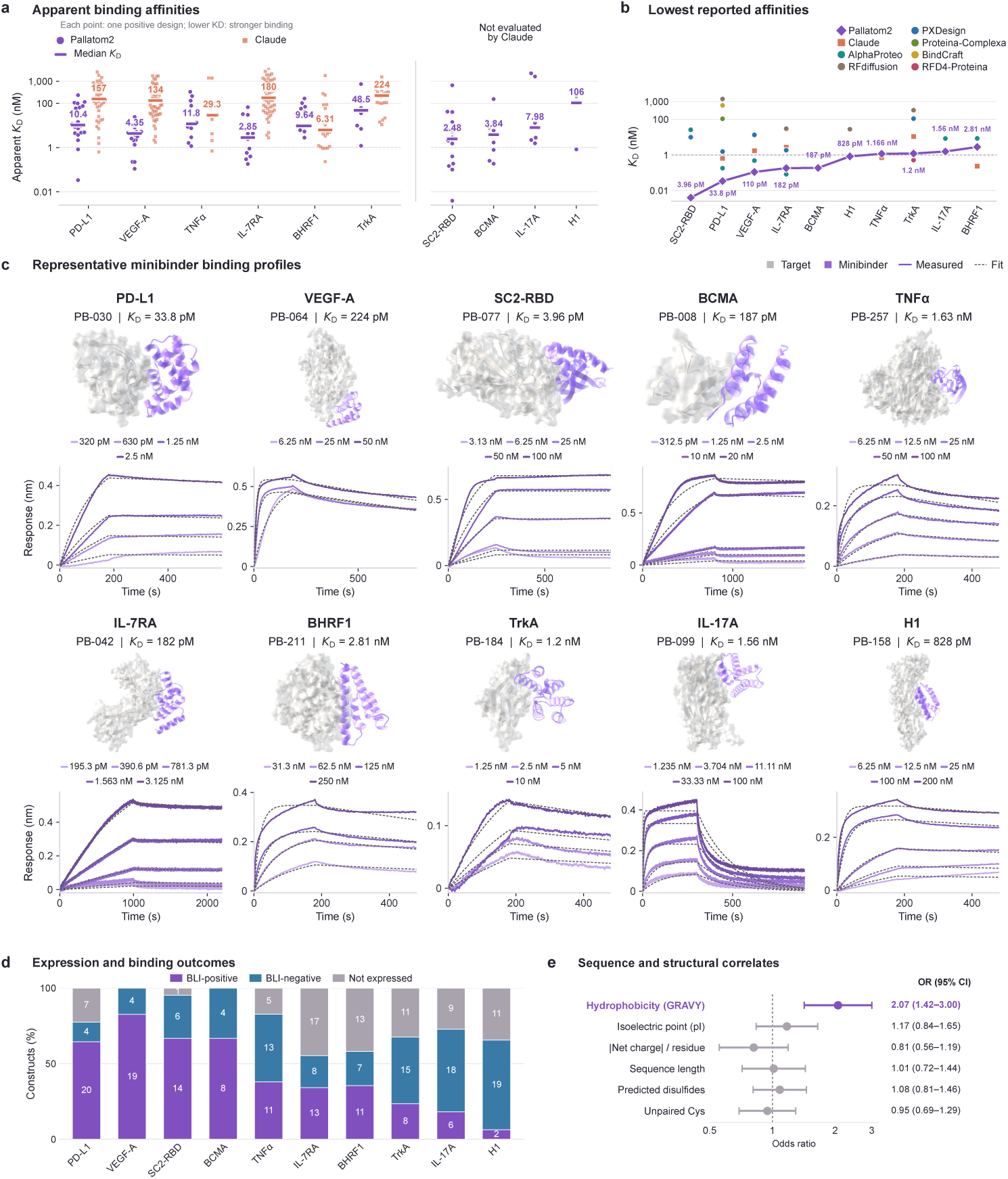
| Minibinder activity, affinity and expression outcomes across ten targets. (a) Apparent *K_D_*values for 112 active Pallatom2 designs and 197 numerical Claude records. (b) Per-target minimum apparent *K_D_*alongside published measurements (Supplementary Table S3). (c) Predicted complexes and BLI for representative designs. (d) Final outcomes for 284 designs: 112 active, 98 inactive and 74 nonexpressed. (e) Associations between six sequence and structural features and nonexpression (*n* = 248).

These candidates came from a much smaller generation campaign: 728–2,400 samples were recorded per target (Supplementary Fig. S5b). Claude’s public protocol [19] targets 20,000–200,000 designs for a single-target campaign and 100,000–1,000,000 for a 14-target campaign. Even without treating these ranges as executed counts, the sampling scales differ by approximately one to nearly three orders of magnitude. Across the six shared targets, Pallatom2 had the lower median *K_D_* for 5 of 6 targets and the lower minimum *K_D_* for 4 of 6 (Fig. 12a,b). After pooling the six shared targets, its active fraction was 44.1% (82/186), compared with 32.8% (197/600) for Claude, and it was higher in 5 of 6 targets. The lowest apparent *K_D_* values reached 3.96 pM for SC2RBD and 33.8 pM for PD-L1.

Expression remained a major bottleneck. Protein expression was unsuccessful for 74 of the 284 evaluated candidates (26.1%). We jointly modelled mean hydrophobicity, pI, absolute charge density, length, predicted disulfide count and unpaired cysteine count, adjusting for target and source era. Each standard deviation increase in hydrophobicity gave nonexpression odds of 2.07 (95% CI, 1.42–3.00; *q* = 0.000889); the other features showed no clear independent association (Fig. 12e). Hydrophobicity was therefore the dominant sequence feature associated with *de novo* minibinder expression in this cohort. This pattern is consistent with a model trained without distilled data, potentially favouring more natural-like sequence composition, and a nomination process that did not filter on expression or hydrophobicity, exposing sequence-associated expression liabilities in the experimental cohort.

Pallatom2 yielded experimental binders for all ten targets. Its 112 numerical *K_D_* records spanned 3.96 pM to 2.29 *µ*M, including picomolar-scale apparent *K_D_* records for six targets. Target-level median *K_D_* ranged from 2.48 to 105.86 nM. Representative predicted complexes paired with concentration-series BLI connect design models directly with measured responses (Fig. 12c); target-specific structure–BLI pairs are provided in Supplementary Figs. S6–S15.

### 4.2. Starting-interface-guided BCMA design preserves binding kinetics

To test whether starting-interface properties can be retained without retaining the original sequence, we used BCMA binder PR-001 as an interface template for sequence generation constrained only by the starting-interface geometry. BCMA is a CAR-T antigen [20, 21], for which recognition-module association and dissociation kinetics have functional relevance [22, 23, 24]; PR-001 uses the Arcellx BC40 reference sequence [15]. This section therefore tests whether interface transfer preserves the starting association rate *k*_on_ while *k*_off_ is reshaped.

Starting-interface-guided design reuses no BC40 sequence. Instead, it conditions generation on side-chain geometry at eight PR-001 interface sites (Fig. 13a). These sites may move or reorder within a new sequence while retaining their relative interface information. The five active interface-guided designs share only 12.5–37.3% sequence identity with BC40, and PB-002 and PB-003 reorder the template sites along the sequence (Fig. 13b). They therefore directly test decoupling starting-interface properties from the original sequence.

**Fig. 13.**
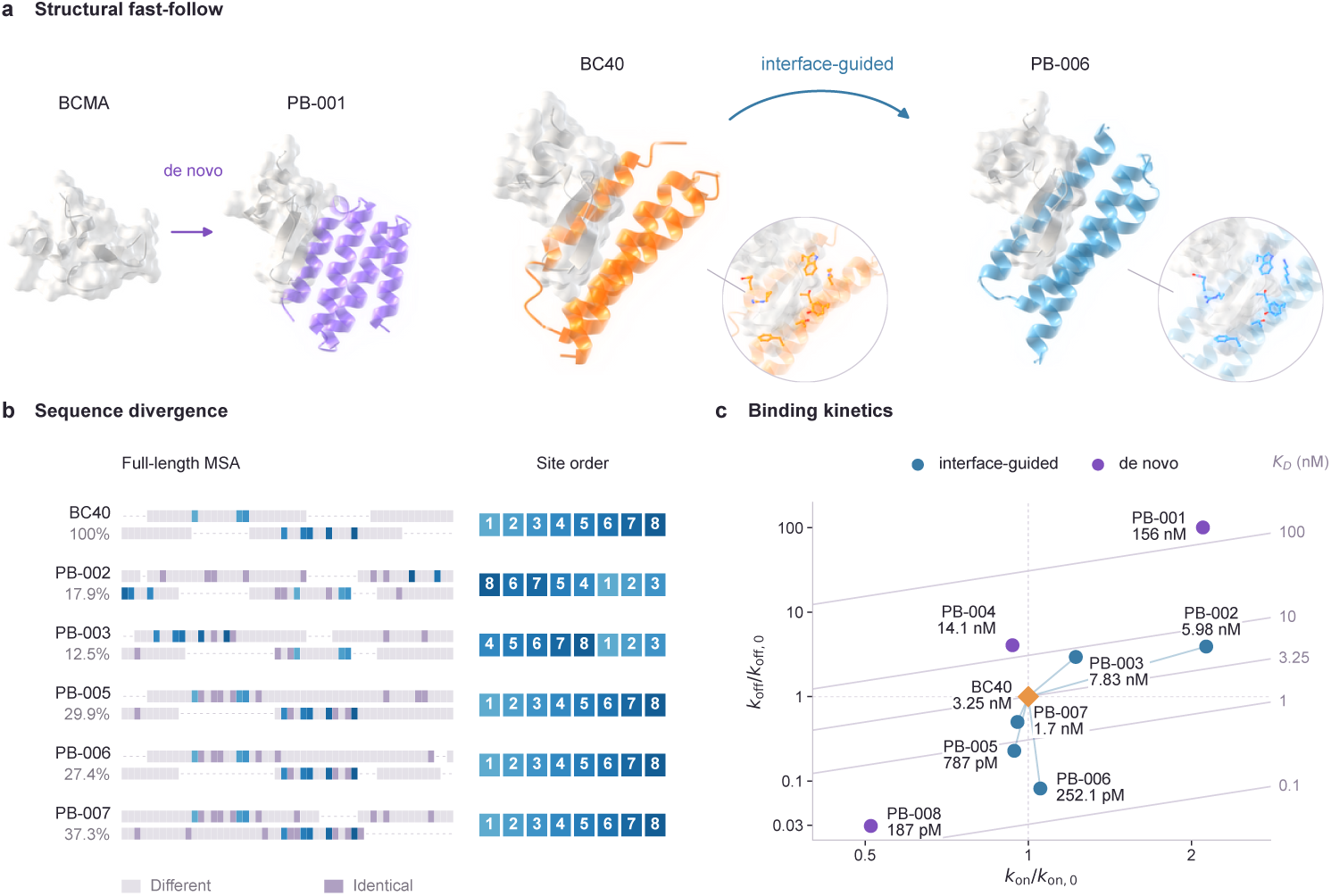
| Starting-interface-guided design preserves binding kinetics. (a) *De novo* and starting-interface-guided routes. (b) Sequence alignment of BC40 and five interface-guided designs. (c) Relative *k*_on_–*k*_off_ map with apparent-*K_D_*labels.

The twelve BCMA designs tested comprised six starting-interface-guided designs and six *de novo* designs. Five of the six interface-guided designs showed binding activity, compared with three of the six *de novo* designs. The starting binder PR-001 had *k*_on_ = 3.24 *×* 10^5^ M*^−^*^1^s*^−^*^1^ and *k*_off_ = 1.05 *×* 10*^−^*^3^ s*^−^*^1^. All five interface-guided designs retained a starting-order *k*_on_ (0.94–2.13-fold), whereas *k*_off_ was reshaped to 0.082–3.92-fold of the starting value. PB-005, PB-006 and PB-007 reduced *k*_off_ to 0.23-, 0.082- and 0.50-fold, respectively, and PB-006 reached an apparent *K_D_* of 252.1 pM (Fig. 13c; Supplementary Fig. S6). Site rearrangement and low sequence identity did not destroy the starting association rate; affinity differences arose primarily from dissociation.

*De novo* design also yielded a high-affinity binder: PB-008 had an apparent *K_D_* of 187 pM, but its *k*_on_ was 51% of that of PR-001, indicating a kinetic profile distinct from that of the starting binder. By comparison, the interface-guided group remained closer to PR-001: *k*_on_ clustered near the starting value, whereas *k*_off_ was reshaped over a broader range. Starting-interface-guided design therefore provides a structural fast-follow strategy: it carries forward a validated binding mode while exploring new molecular space through low-identity and site-rearranged designs. The same principle could support fast-follow across modalities, including transfer of an antibody binding interface to a cyclic peptide or minibinder.

### 4.3. In silico affinity maturation yields potent TNF*α* binders

After obtaining primary binders, we applied partial diffusion for in silico affinity maturation. Starting from TNF*α* binders PB-121 (13.37 nM) and PB-128 (78.78 nM), we nominated 24 variants, of which 21 were active, two inactive and one nonexpressed (Fig. 14a; Supplementary Fig. S16). Global sequence identity assigned variants to groups nearest PB-121 and PB-128, each with 12 nominations and yielding ten and eleven active designs, respectively.

**Fig. 14.**
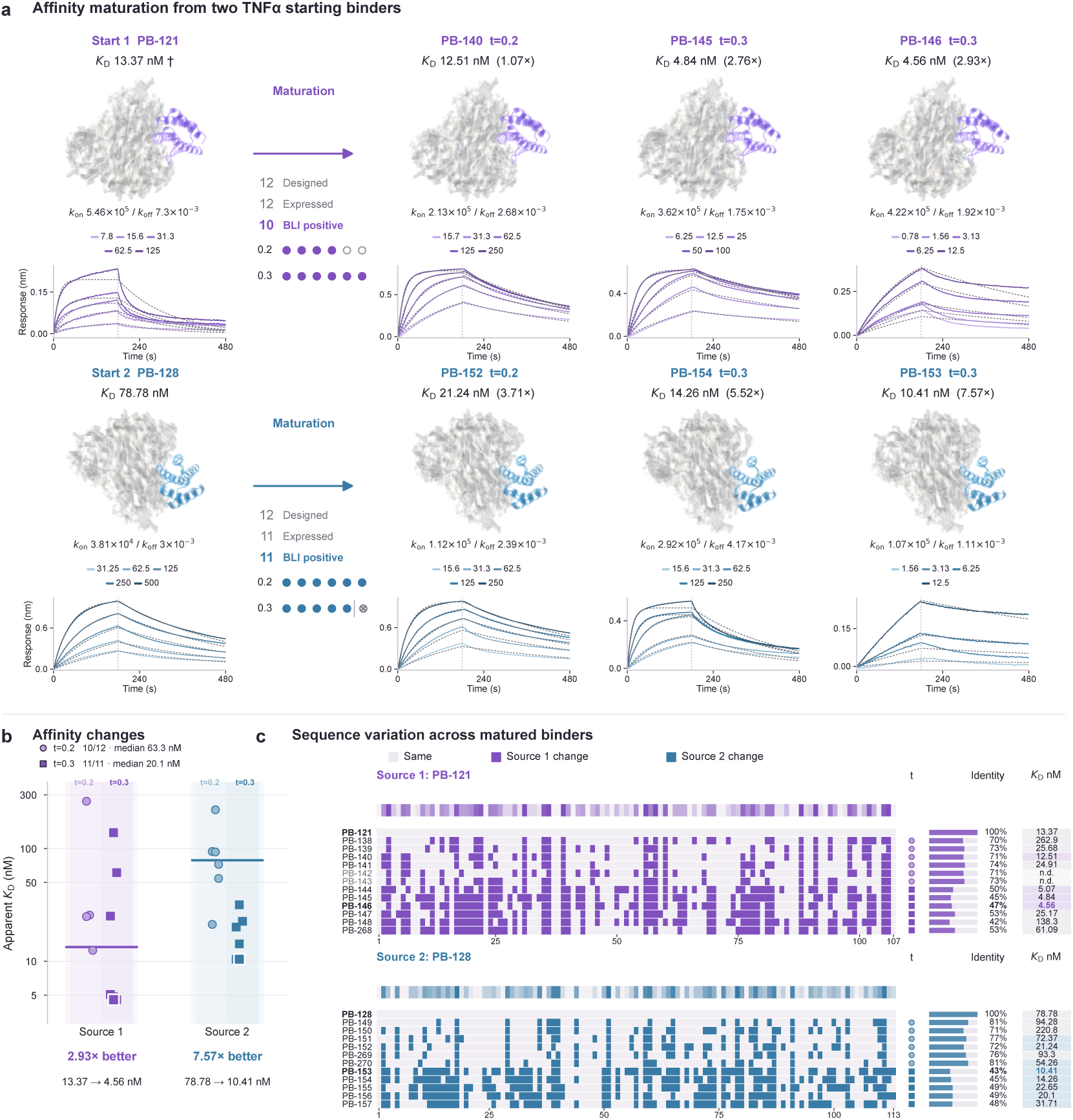
| In silico affinity maturation yields potent TNF*α* binders. (a) Predicted complexes and BLI for starts and representative variants. (b) Variant *K_D_* distributions relative to their starts. (c) Anonymous substitutions, site frequencies, sequence identity and *K_D_*.

Gains extended beyond the single best variant: four and eight variants in the two groups had lower apparent *K_D_* than their assigned starts. The best variants, PB-146 and PB-153, reached 4.56 and 10.41 nM, corresponding to 2.93- and 7.57-fold improvements (Fig. 14a,b). PB-146 retained approximately 77% of the starting *k*_on_ while *k*_off_ fell to approximately 1/3.8 of the starting value. PB-153 showed an approximately 2.8-fold increase in *k*_on_ and a decrease in *k*_off_ to approximately 1/2.7. Affinity gains thus arose through different rate changes.

The two partial-diffusion settings differed within the tested set. At *t* = 0.3, 12 variants were nominated; 11 completed BLI and all 11 were active, with a median active *K_D_* of 20.1 nM. At *t* = 0.2, 10 of 12 designs were active, with a median active *K_D_* of 63.315 nM. The *t* = 0.3 median was lower within both sequence groups, and both best variants used this setting (Fig. 14b). Thus, *t* = 0.3 was the better setting in this screen.

Position-wise comparisons showed that affinity gains could accompany substantial sequence change (Fig. 14c). The two best variants retained only 46.7% and 43.4% identity to their assigned starts, whereas the two inactive variants retained 71.0% and 72.9%; higher sequence conservation did not ensure binding.

### 4.4. Generative rescue unlocks nanobody binding

We extended generative optimization beyond affinity maturation by rescuing an inactive nanobody. The starting molecule, PB-272, was a previously generated but unpublished *de novo* nanobody with no detectable TNF*α* binding at 2,000 nM. Remodeling its CDR-associated interface produced 500 candidates. In silico screening nominated ten, of which seven completed expression and BLI testing.

PB-276 and PB-277 bound TNF*α* with apparent *K_D_* values of 4.31 and 5.93 nM, respectively (Fig. 15a). Both variants produced concentration-dependent BLI responses across 6.25–100 nM, establishing binding well below the 2,000 nM concentration at which the start remained inactive. CDR-associated interface remodeling therefore converted an inactive starting molecule into two high-affinity nanobodies.

**Fig. 15.**
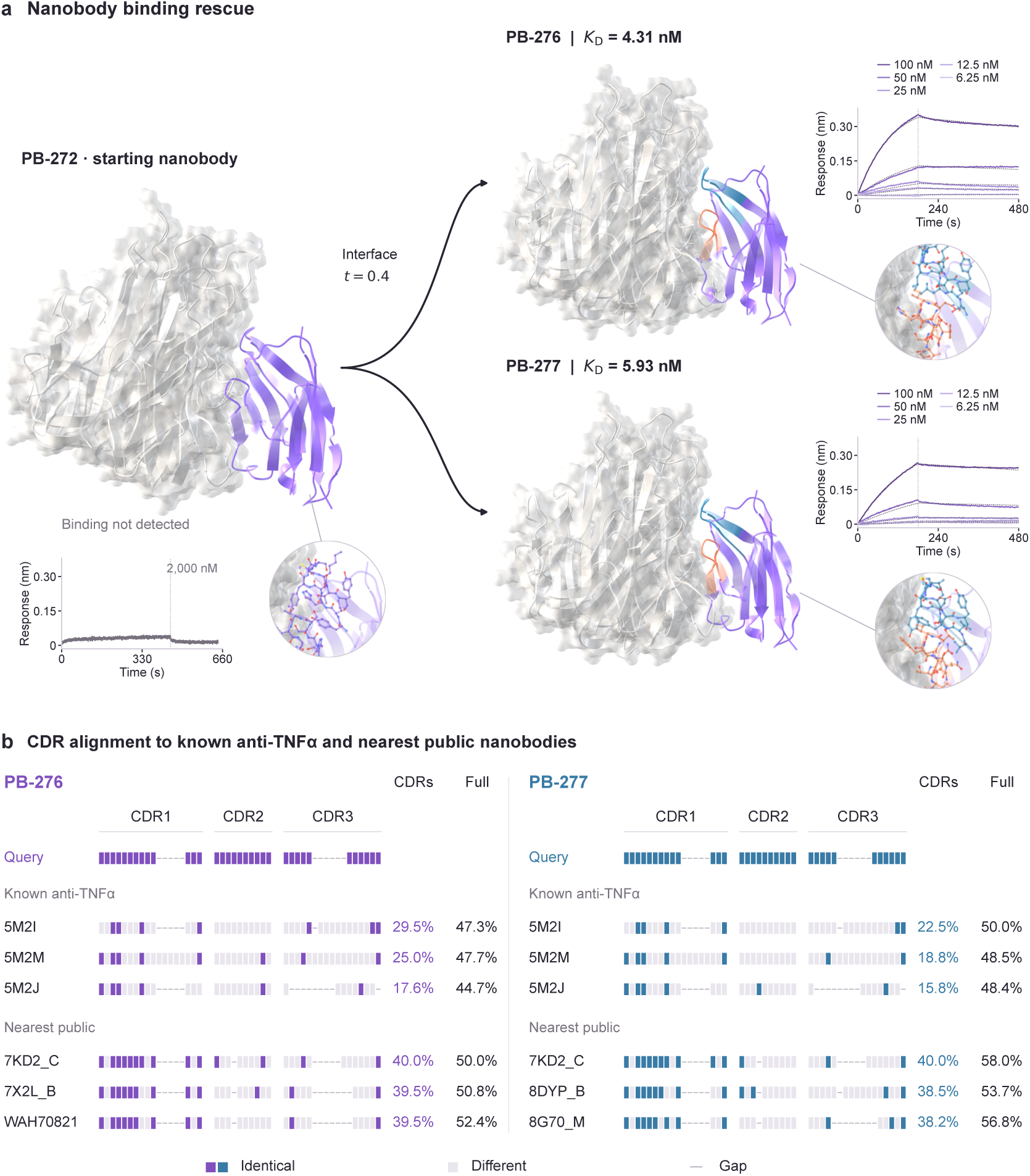
| Generative rescue unlocks nanobody binding. (a) Predicted complexes, interface closeups and BLI for PB-272, PB-276 and PB-277. (b) CDR alignments to the three closest known anti-TNF*α* nanobodies and three nearest public nanobodies for each rescue variant, ranked by combined CDR identity. The right-hand columns report pairwise CDR and full-sequence identity to the same reference.

The rescued variants combine high-affinity binding with CDR sequences distinct from those of known anti-TNF*α* nanobodies. Combined CDR identity to the closest known anti-TNF*α* reference was 29.5% for PB-276 and 22.5% for PB-277; the maximum across the public nanobody set was 40.0% for both (Fig. 15b). These results establish nanobody rescue as a direct extension of generative interface optimization beyond affinity maturation.

## 5. Experimental methods

### 5.1. Binder generation and in silico screening

Minibinders were generated using the target-conditioned Pallatom2 workflow. The ten targets recorded 12,428 samples, corresponding to 728–2,400 per target. Counts used unique generation samples. Structures passing the successful-fold flag were retained using the same criteria as the in silico evaluation: **allatom_rmsd *≤* 2.5 Å**, **min-ipAE *≤* 1.5 Å**, and **ptm-binder *≥* 0.8**. These denote all-atom RMSD between generated and repredicted structures, minimum inter-chain predicted aligned error (PAE), and binder predicted TM-score (pTM), respectively. Passing candidates were ranked and manually nominated considering structural quality and design diversity. Expression predictors, hydrophobicity thresholds and experimental expression outcomes were not used during generation or nomination.

BCMA *de novo* designs used binder lengths of 50–75 residues. Starting-interface-guided generation additionally conditioned on side-chain geometry at eight PR-001 (BC40) interface residues, with binder lengths of 55–85 residues. The supplied BCMA generation records contained 900 samples. All BCMA structures used the same structural criteria described above, and both routes advanced six designs to testing.

### 5.2. Affinity maturation and nanobody rescue

TNF*α* affinity maturation started from PB-121 (13.37 nM) and PB-128 (78.78 nM) using partial diffusion. Twenty-four variants were nominated: 12 at *t* = 0.2 and 12 at *t* = 0.3. Structural outputs used the same filters as minibinder generation, followed by manual review.

Partial diffusion remodeled the CDR-associated interface of PB-272 at *t* = 0.4, producing 500 candidates. Candidates were evaluated independently with AlphaFold3 and OpenDDE (abag model) [25] using the same three structural criteria. Candidates passing both evaluations were ranked by ascending AlphaFold3 min-ipAE, and the top ten were nominated.

Public nanobody references used the PLAbDab-nano snapshot downloaded on 4 November 2026 [26]; 1,968 sequences received complete three-CDR annotations. The known anti-TNF*α* reference panel comprised PR-007 and nanobodies from structures 5M2I, 5M2J, 5M2M and 8Z8M; identical variable-domain sequences were merged. All CDRs used PyIgClassify modified AHo definitions [27]. Each CDR was globally aligned with BLOSUM62 (gap open *−*10, extension *−*0.5), and combined CDR identity was the summed exact-match count divided by the summed alignment length, including gaps. For each rescue variant, the three highest-identity distinct sequences were selected independently from the known anti-TNF*α* panel and the public reference set. Full-sequence identity used the complete variable-domain sequences of the same query–reference pair. Display alignments used one MAFFT G-INS-i multiple alignment per CDR across both rescue variants and all selected references, with shared alignment columns; reported identities used independent pairwise alignments. AlphaFold3-predicted complexes were aligned through receptor C*α* atoms in symmetry-equivalent TNF*α* chains.

### 5.3. Protein expression and BLI

Genes were synthesized by Tsingke, cloned into pET-24a, verified and transformed into *Escherichia coli* BL21(DE3). Constructs carried a C-terminal His6 tag. Single colonies were cultured in 5 mL kanamycin-containing LB at 37 *^◦^*C and 250 rpm to OD600 0.4–0.6, transferred to fresh LB and grown for 3 h at 37 *^◦^*C and 220 rpm. Expression was induced with 0.5 mM IPTG overnight at 16 *^◦^*C and 220 rpm. Cells were lysed for 10 min and centrifuged at 12,000 rpm for 10 min. Supernatants were loaded onto Ni-NTA columns equilibrated with five column volumes of 20 mM phosphate, 0.5 M NaCl and 5 mM imidazole, pH 7.4. Columns were washed with five volumes and eluted with three volumes containing 500 mM imidazole. SDS–PAGE inspection was specified in the workflow.

BLI measurements used an Octet R8 instrument. Target proteins and tag formats are listed in Supplementary Table S4. H1 and BHRF1 were chemically biotinylated in-house before loading. All targets were captured on streptavidin (SA) biosensors, and designed binders were tested as analytes at different concentrations. Each cycle comprised a 60 s baseline, target loading, a 120 s wash, and analyte association and dissociation. The assay buffer was PBS containing 0.02–0.05% Tween-20 and 0.1% BSA. Curves were fitted in Octet analysis software using a 1:1 binding or mass-transport model to obtain *k*_on_ and *k*_off_; apparent *K_D_* = *k*_off_/*k*_on_.

### 5.4. Sequence and expression analyses

Full-length sequences were aligned with MAFFT 7.526 G-INS-i. Figure 13b identity is the fraction of identical residues among columns containing residues in both sequences. Structural-site mapping was independent of sequence alignment. TNF*α* sequences were globally aligned with Biopython PairwiseAligner using match 2, mis-match *−*1, gap-open *−*10 and gap-extension *−*0.5. Each maturation variant was assigned to the *de novo* design with maximum identity; nominal affinity gain was the assigned starting *K_D_* divided by variant *K_D_*.

GRAVY, pI, absolute net charge at pH 7 and sequence length were calculated with Biopython. Predicted disulfides used an intrabinder Cys SG distance cutoff of 2.5 Å, pairing each cysteine at most once; remaining cysteines were counted as unpaired. Six features were standardized by the fitted-sample SD and entered a logistic model with target intercepts and source-era adjustment. Odds ratios per SD, Wald 95% CIs and BH-adjusted *q* values are reported; BCMA and VEGF-A were excluded from coefficient estimation, leaving *n* = 248.

## 6. Discussion

Pallatom2 was developed to make protein design directly programmable at atomic resolution, coordinating molecular conformation with the contacts needed for target recognition. The results demonstrate broad in silico capability across long-chain monomers, multi-domain proteins, homomultimers, and target-conditioned binders. Long-chain sampling extends to 1,600 residues, while binder design achieves state-of-the-art screening performance across the nine targets with cross-method benchmark data under the reported protocols. High screening success rates for BHRF1 and the challenging target TNF*α* highlight the model’s binder-design capability, while performance remains dependent on target-specific interface difficulty (Fig. 9).

These in silico capabilities translate into experimentally measurable binding across all ten tested targets. Among 210 candidates evaluated by BLI, 112 were active, with picomolar-scale apparent affinities obtained for six targets, including 3.96 pM for SC2RBD and 33.8 pM for PD-L1; TNF*α* reached 1.166 nM (Fig. 12). The experimental evidence also extends beyond *de novo* discovery. Starting-interface-guided BCMA design retained association kinetics despite substantial sequence divergence, partial diffusion improved TNF*α* affinity by up to 7.57-fold, and interface remodeling converted an inactive nanobody into binders with apparent *K_D_* values of 4.31 and 5.93 nM. Together, these results support a common generative framework for discovering, transferring, and refining molecular recognition.

The architectural basis of this framework is direct modeling of atom-level single and pair information. The coordinate-sensitive AtomPairFormer combines interatomic distances, directions, and bonding relationships in a 384-channel pair representation, allowing backbone, side-chain, and interface geometry to be considered within the same modeling process. Residue-guided sparse attention restricts detailed atom-to-atom information exchange to selected relationships, making this representation more tractable as molecular size increases. Strong screening performance with 65.9 million parameters indicates that a compact model can support demanding design tasks (Fig. S2).

Programmable conditions turn this atomic representation into a means of expressing design intent. Interface controls specify where recognition should occur, which contacts should be avoided or retained, and which regions may adapt, while structure controls define the conformations and local geometries available to realize those requirements. The BCMA experiments illustrate the value of conditioning on an existing interface without requiring retention of its sequence: five active interface-guided designs preserved association rates near that of the starting binder while exploring different dissociation kinetics (Fig. 13). TNF*α* maturation and nanobody rescue further show that selective structural modification can improve existing candidates or recover binding from an inactive design. This connects *de novo* generation and redesign within a shared, controllable molecular search space.

The training-data and sequence-composition results provide another perspective on designability. Pallatom2 uses TED and AFDB structures with their associated sequences, without ProteinMPNN sequence redesign during training-data preparation. Its high in silico screening success coexists with a binder amino acid distribution closer overall to the UniProt reference than the displayed ProteinMPNN distribution (Fig. 9b). These observations support the feasibility of combining strong designability with relatively natural-like sequence composition. They do not isolate the contribution of training-data preparation from architecture or sampling, motivating controlled ablations of these components and broader evaluation of the properties of generated sequences.

Generation, search, and candidate selection offer complementary opportunities for further improvement. Pallatom2 search explores branches of the generation trajectory, while in silico screening, interface assessment, and Lévin™ Harness support prioritization of completed candidates. The supplementary analyses identify minimum inter-chain PAE as a major screening bottleneck and associate lower interface error with more extensive and confident inter-chain contacts (Figs. S3 and S4). These observations motivate testing search criteria that account for interface engagement and atomic geometry alongside fold confidence. Prospective comparisons with fixed sampling and experimental budgets will be needed to quantify the separate contributions of trajectory search and agent-assisted review.

The current results also define the next experimental priorities. The 53.3% active fraction is conditional on the 210 candidates that completed BLI testing; 74 of the broader 284-candidate cohort were nonexpressed. Improving expression and calibrating candidate selection against experimental outcomes therefore remain necessary, even when in silico screening is favorable. Apparent affinities measured by BLI require complementary characterization of specificity, stability, aggregation, and target-dependent function, while experimental structures would test whether the intended atomic contacts are realized. Extending validation across additional target classes and molecular formats will establish how broadly the demonstrated capabilities transfer.

Pallatom2 provides a foundation for linking atomic design specifications to experimentally testable proteins. Its combination of direct atom-level modeling, programmable conditions, and experimentally demonstrated interface optimization supports a workflow in which successful and unsuccessful designs can inform subsequent hypotheses. Further integration of these capabilities with systematic experimental feedback could improve both the reliability of protein engineering and our understanding of how sequence, geometry, and molecular recognition are connected.

## S1. Training datasets

Pallatom2 was trained on both monomer and protein-complex data derived from the AlphaFold Protein Structure Database (AFDB) and The Encyclopedia of Domains (TED) [28, 29]. These resources provide predicted protein structures and domain annotations at different levels of structural organization. The training set was assembled through an internal processing pipeline. **We retained the original sequences associated with these structures, without ProteinMPNN-based or other sequence redesign during training-data preparation.**

### TED: domain-level structural information

TED identifies and classifies structural domains within AFDB protein models using automated domain parsing and structure comparison. Its domain boundaries and classifications describe the constituent folding units of proteins and their relationships to known structural families. This domain-level organization provides a structured view of the diversity of protein folds [28].

### AFDB: predicted protein structures

AFDB provides AlphaFold-predicted three-dimensional protein structures together with the corresponding sequences and prediction-confidence information. These models describe protein-chain conformations and provide structural context beyond individual domain boundaries [29].

TED is derived from AFDB, so the two resources are related views of predicted protein structure rather than independent collections of experimentally determined structures. Here, TED supplies the domain-level description and AFDB supplies the underlying protein-structure resource.

### ProteinMPNN redesign improves designability but shifts sequence composition

MultiFlow replaces the original sequences of its PDB training structures with ProteinMPNN-designed sequences selected for sequence–structure consistency. Proteina Atomistica similarly constructs a synthetic training set by redesigning sequences on AFDB-derived structures, followed by refolding and consistency filtering. Both studies report improved in silico co-designability from their respective data-distillation procedures [30, 31]. However, this strategy shifts the training-sequence distribution toward ProteinMPNN preferences, which can propagate to the amino acid composition of generated proteins. Proteina Atomistica explicitly reports enrichment of particular residue types, including glutamate and lysine, relative to its natural-sequence dataset [31].

### High designability with a natural-like amino acid distribution

Pallatom2 achieves high in silico designability using training data without sequence redesign. In our binder benchmarks, the strong screening performance in Figure 9a coexists with an amino acid distribution that is closer overall to the UniProt reference than the displayed ProteinMPNN distribution (Figure 9b). Pallatom2 thus combines high in silico designability with a natural-like amino acid composition, without relying on ProteinMPNN-redesigned training sequences.

### S1.1. Domain-level diversity in TED

The TED-derived training data used in this study were generated through an internal processing pipeline and are not identical to the official TED dataset. In this dataset, most domains are compact folding modules: the median original-domain length was 119 aa in the single-domain group and 131 aa in the multidomain group, while median domain length remained between 119 and 138 aa as the number of domains in the parent structure increased (a and f). Thus, individual domains did not become systematically longer as overall structures became more complex.

Longer protein structures instead arise primarily from combinations of multiple short domains. The domain-weighted cumulative distribution of total residue counts shows that structures containing more domains contribute greater weight to the overall length distribution (e), while domain length remains largely stable across parent structures of different sizes (a and f). Protein length therefore increases mainly through the accumulation of domains rather than through progressive elongation of individual domains. The TED-derived training set consequently covers both isolated compact folding modules and larger multidomain structures assembled from multiple short domains.

**Fig. S1.**
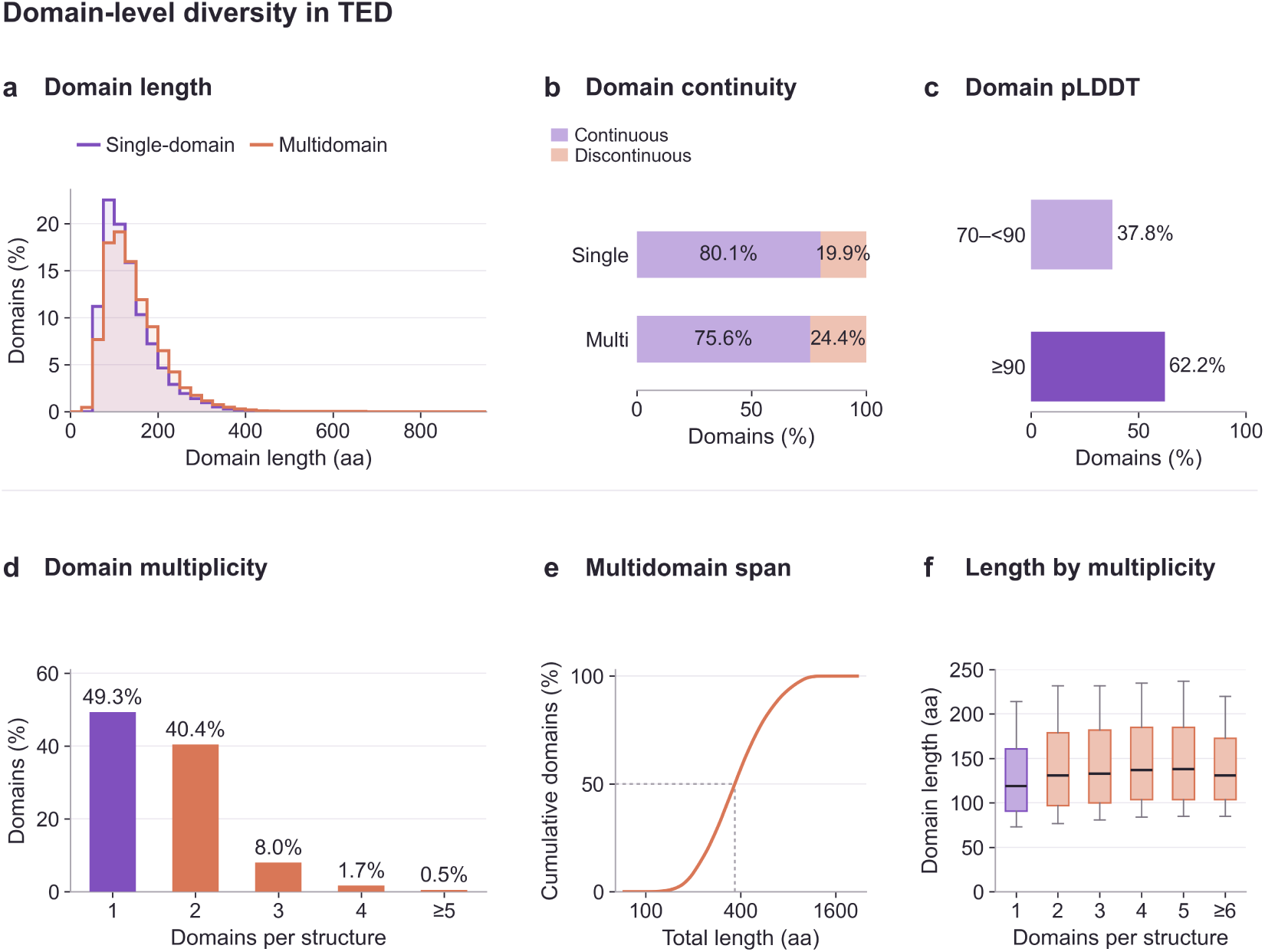
| Domain-level diversity in the internally processed TED-derived training data. **a,** Original-domain length distributions for the single-domain (purple) and multidomain (orange) groups, normalized separately. **b,** Continuous and discontinuous domains. **c,** Distribution of mean pLDDT across original domains. **d,** Domain proportions grouped by the number of domains in the parent structure. **e,** Domain-weighted cumulative distribution of total residue counts in processed multidomain structures. **f,** Original-domain lengths grouped by parent-structure domain count. Boxes show the interquartile range, central lines the medians and whiskers the 10th–90th percentiles.

## S2. Conditional control for protein design

**Tab. S1.** | Pallatom2 binder–target interface control conditions.

| Condition | Design intent | Typical use case |
|---|---|---|
| Hotspot | Required target epitope | Force burial or recognition of a functional region. |
| Coldspot | Forbidden target region | Avoid conserved sites, glycosylation sites, or off-target surfaces. |
| Binderspot | Required binder-side contact | Ensure selected binder residues enter the interface. |
| Flexspot | Flexible target segment | Permit loop or local side-chain rearrangement during binding. |
| Unindex | Known side-chain geometry | Transfer validated atomic interactions into a generated binder. |

**Tab. S2.** | Pallatom2 all-atom structural control conditions.

| Condition | Primary control | Design role |
|---|---|---|
| motif | Structural motif and backbone pattern | Specify a structural element or arrangement to be represented in the candidate. |
| redesign | Designable residues or regions | Define which parts of the structure are available for redesign. |
| norepack | Selected side-chain conformations | Preserve designated side chains against repacking. |
| slevel | Global or fragment-level secondary-structure composition | Set desired secondary-structure proportions. |
| sse | Secondary-structure state | Specify secondary-structure character for selected positions or segments. |
| seq_cond | Residue identity and sequence | Condition amino-acid identities at designated positions. |
| hb_bond | Hydrogen-bond geometry | Specify hydrogen-bond relationships between relevant atoms. |

## S3. Supplementary sampling analyses

### S3.1. Model size and in silico screening performance

#### High screening success with a compact parameter budget

Pallatom2 contains 65.9 million parameters, compared with 435.6 million for RFD4-Proteina, 496 million for BoltzGen, and 93 million for the AF2 model used in BindCraft (Figure S2). Its parameter count is approximately 15.1%, 13.3%, and 70.9% of these respective model sizes. Despite this smaller parameter budget, Pallatom2 maintains strong in silico screening performance in the binder-design benchmark (Figure 9a), reaching success rates of 98.5% for BHRF1, 94.9% for PD-L1, and 93.8% for TrkA, with 35.6% on the challenging target TNF*α*. Together, these results show that high in silico screening success can be achieved with a comparatively compact model, supporting the effectiveness of Pallatom2’s all-atom modeling architecture.

**Fig. S2.**
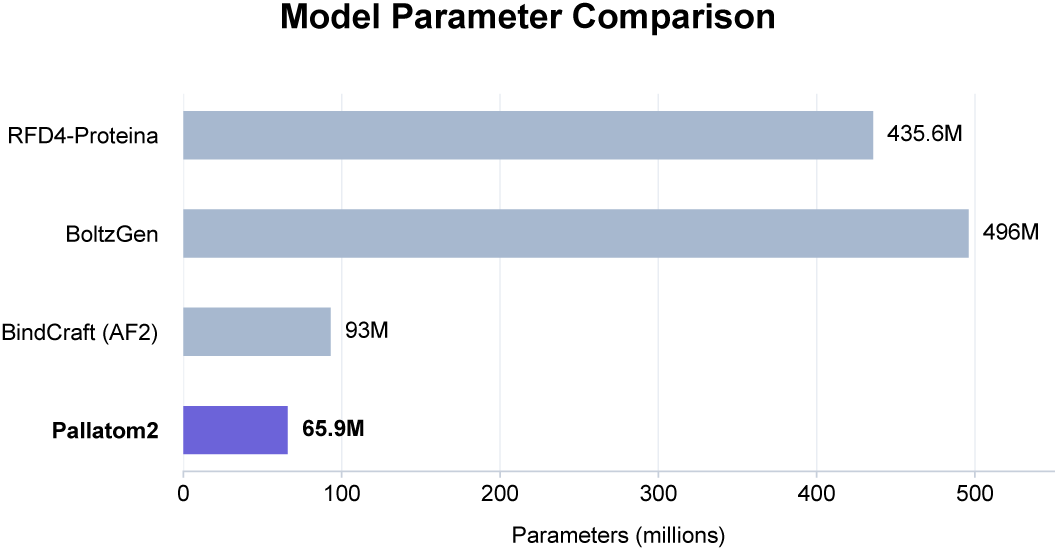
| Model parameter comparison. Pallatom2 has 65.9M parameters, compared with 435.6M for RFD4-Proteina, 496M for BoltzGen, and 93M for BindCraft’s AF2 model. M denotes one million parameters. Pallatom2 is highlighted in purple. The associated in silico screening results are reported in Figure 9a.

### S3.2. Supplementary interface diagnostics

#### Joint metrics reveal distinct candidate populations

Figure S3a resolves the screening landscape by plotting all-atom RMSD against minimum inter-chain predicted aligned error, with color indicating binder pTM. BHRF1, PD-L1, and TrkA show prominent clusters with both low RMSD and low interface error, whereas TNF*α*, SC2RBD, and VEGF-A also contain substantial populations with high interface error. High-pTM candidates occur in both low– and high-error regions, indicating that confidence in the binder fold alone does not establish consistency of the designed complex or its binding interface. The lower-left region satisfies the two geometric thresholds, while pTM remains a separate screening dimension. These distributions explain why joint evaluation is needed to distinguish internally confident folds from candidates that also reproduce the intended complex geometry.

#### Interface confidence is a major screening bottleneck

Figure S3b separates the pass rates for all-atom RMSD, minimum inter-chain PAE, and binder pTM. The min ipAE criterion has the lowest individual pass rate in nine of the ten cases. This limitation is most pronounced for TNF*α*, VEGF-A, and SC2RBD, where only 5%, 13%, and 18% pass the interface-error threshold, respectively, compared with RMSD pass rates of 33%, 60%, and 47% and binder pTM pass rates of 59%, 65%, and 79%. H1 is the exception: its pTM pass rate (49%) is slightly below its min ipAE pass rate (52%). These results identify interface confidence as a major current limitation on overall screening success and motivate improvements in target–binder interface modeling alongside binder fold quality. Because all criteria must be satisfied, the joint pass rate cannot exceed the lowest individual pass rate for the same candidate set; the marginal rates alone do not determine the exact joint rate.

**Fig. S3.**
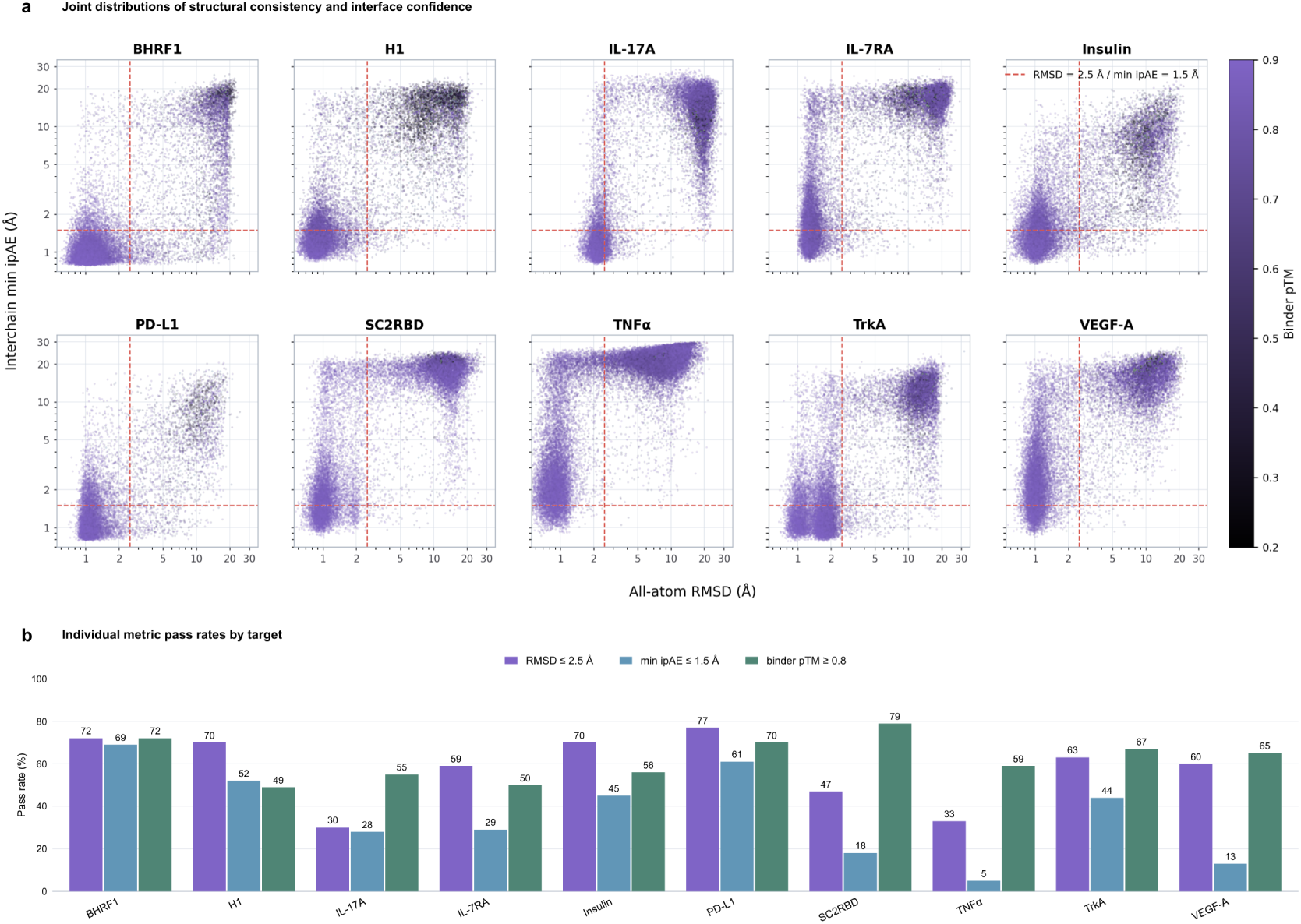
| Supplementary in silico screening diagnostics. (a) All-atom RMSD versus minimum inter-chain PAE on logarithmic axes across ten targets, colored by binder pTM; dashed lines mark 2.5 Å and 1.5 Å, respectively. (b) Individual pass rates for RMSD *≤* 2.5 Å, min ipAE *≤* 1.5 Å, and binder pTM *≥* 0.8. Both panels retain Insulin from their supplied datasets. Panel b reports marginal rates for the supplied metric-analysis set, not joint pass rates; these values are not used to recalculate Figure 9a.

#### Interface geometry offers clues to min ipAE differences

Figure S4 compares candidates with min ipAE *≤* 1.5 Å and *>* 5 Å. Across the nine displayed targets, the low-error group has more heavy-atom contacts, more binder residues participating in the interface, more close polar-atom contacts, and higher interface mean pLDDT. These patterns suggest that differences in interface engagement and atomic packing may contribute to the confidence gap: fewer contacting residues and fewer inter-chain contacts may leave the relative placement of the two proteins less well constrained, while differences in polar-contact opportunities may reflect how side chains are arranged at the interface. The lower interface pLDDT in the high-error group also associates high min ipAE with less confident local structural predictions. Thus, the observed gap may involve both the extent and geometry of the generated interface and the quality of its local structural organization. These group-level associations do not establish causality; close polar-atom contacts alone do not establish hydrogen bonds.

**Fig. S4.**
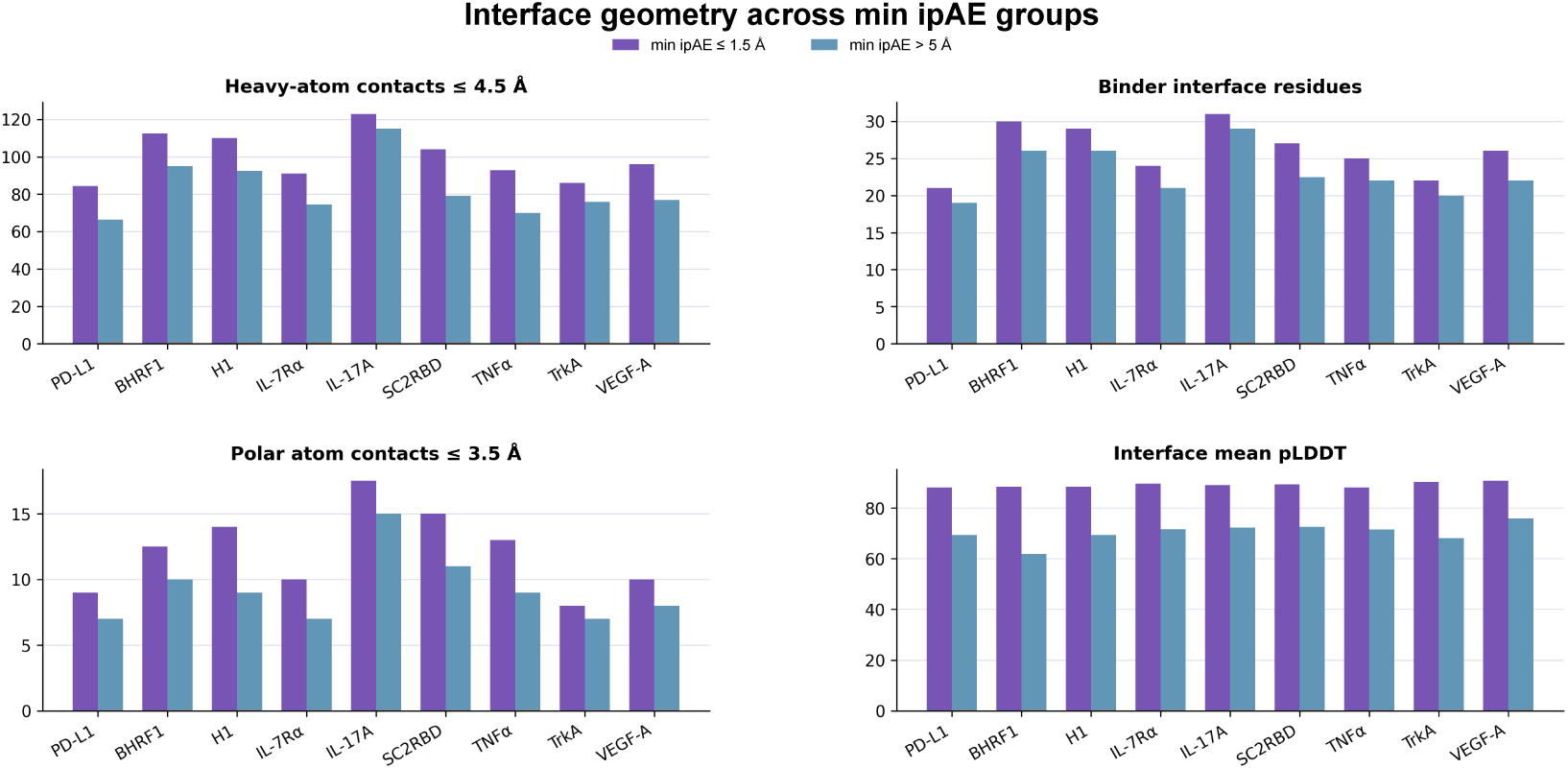
| Interface geometry across min ipAE groups. Interface descriptors for the nine displayed targets, comparing min ipAE *≤* 1.5 Å (purple) with min ipAE *>* 5 Å (blue). The four plots show heavy-atom contacts within 4.5 Å, binder interface residues, polar-atom contacts within 3.5 Å, and interface mean pLDDT. Candidates with intermediate min ipAE are outside this two-group comparison.

## S4. Experimental supplementary data

External comparison sources and target reagents are listed in Supplementary Tables S3 and S4, respectively. Supplementary Figures S5–S16 present experimental binding success, sampling scale and the reviewed molecular/BLI pairs. Available sequences and affinities of active minibinders and reference binders are compiled in Supplementary Table S5 at the end of the Supplementary Information.

**Supplementary Table S3.** Sources and assays for the external affinity comparisons.

| **Method** | **Assay** | **Source** |
|---|---|---|
| Claude | SPR / BLI | [19] |
| RFdiffusion | BLI | [5] |
| AlphaProteo | HTRF | [6] |
| Proteina-Complexa | SPR | [2] |
| BoltzGen | SPR / BLI | [9] |
| BindCraft | SPR | [1] |
| PXDesign | BLI | [32] |
| RFD4-Proteina | SPR | [11] |

**Supplementary Table S4.** Target proteins used for BLI measurements.

| Target | Protein | Tag / format | Vendor | Catalog |
|---|---|---|---|---|
| TNFα | Biotinylated human TNFα, active trimer (MALS verified) | His, AviTag | ACROBiosystems | TNA-H82E3 |
| SC2RBD | Biotinylated SARS-CoV-2 spike RBD (MALS verified) | His, AviTag | ACROBiosystems | SPD-C82E9 |
| PD-L1 | Biotinylated human PD-L1 / B7-H1 (19–134) (MALS verified) | His, AviTag | ACROBiosystems | PDL-H82E4 |
| IL-17A | Biotinylated human IL-17A / CTLA-8 | His, AviTag | ACROBiosystems | ILA-H82Q1 |
| IL-7RA | Recombinant human IL-7RA / CD127 ECD, biotinylated (HPLC verified) | His, AviTag | Sino Biological | 10975-H49H-B |
| VEGF-A | Biotinylated human VEGF165 (HPLC verified) | His, AviTag | ACROBiosystems | VE5-H82Q0 |
| TrkA | Human TrkA ECD, biotinylated | Fc, AviTag | Sino Biological | 11073-H41H-B |
| BCMA | Biotinylated human BCMA / TNFRSF17, premium grade | His, AviTag | ACROBiosystems | BCA-H82E4 |
| H1 | Recombinant influenza A H1N1 (A/Taiwan/01/1986) HA | His | Novoprotein | DRA127 |
| BHRF1 | Recombinant BHRF1 | His | Aipti Biotech | — |

**Supplementary Fig. S5.**
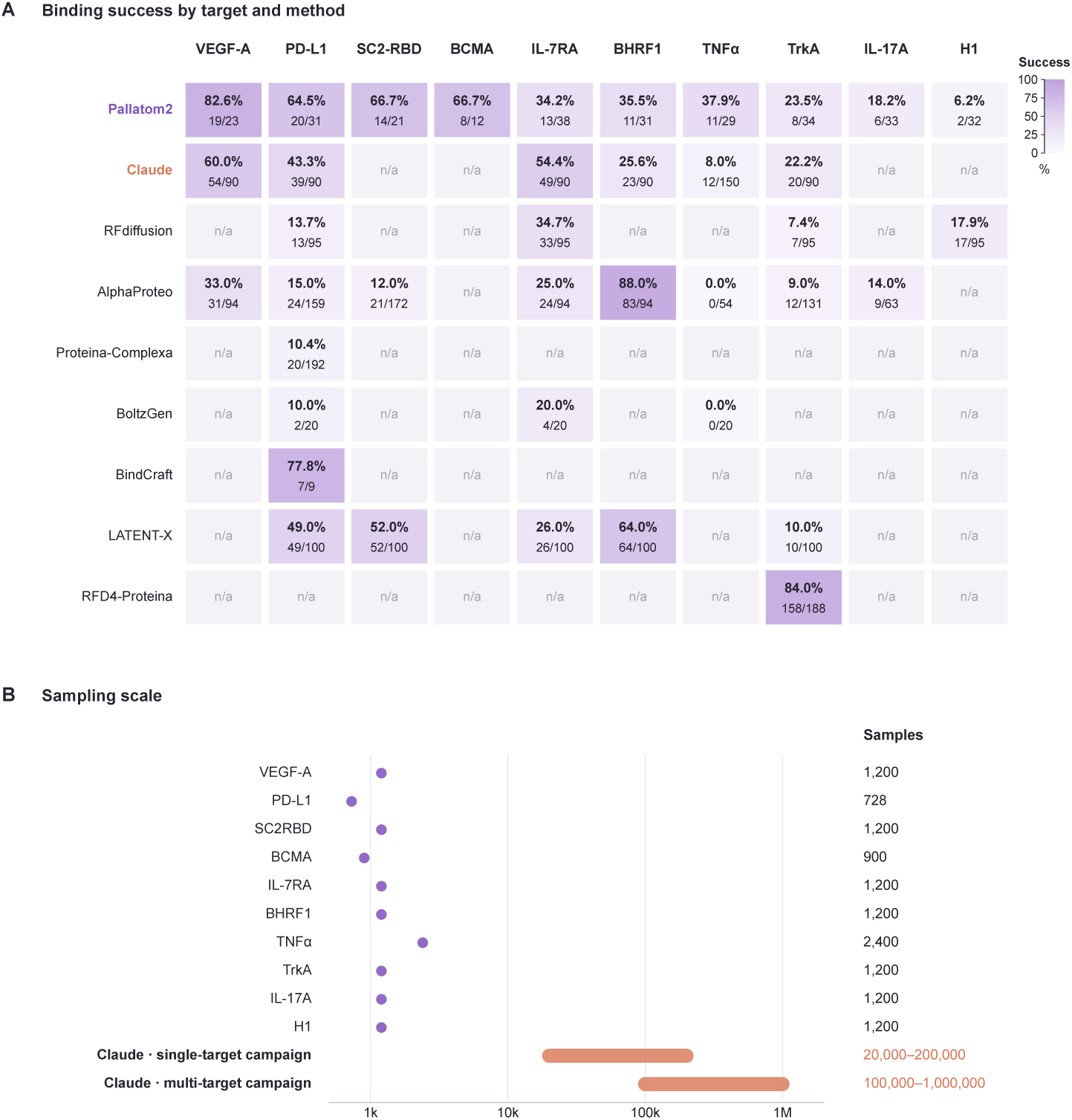
Experimental binding success and sampling scale. **a**, Binding success across ten targets and nine design methods. **b,** Per-target Pallatom2 sampling counts and Claude campaign sampling ranges.

**Supplementary Fig. S6.**
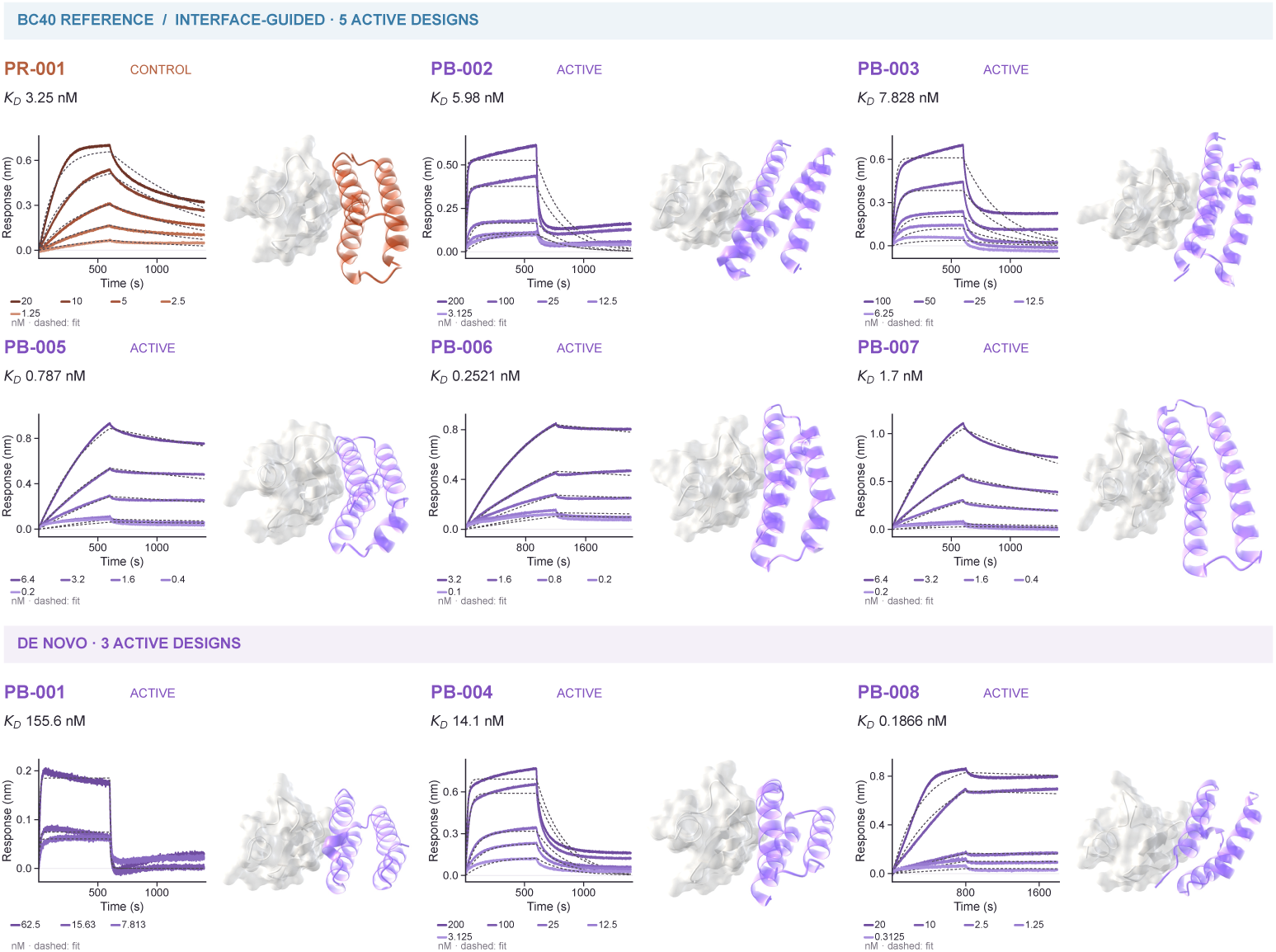
BCMA. Predicted complexes and BLI for five starting-interface-guided and three *de novo* designs. BC40 (PR-001) is included as the reference binder.

**Supplementary Fig. S7.**
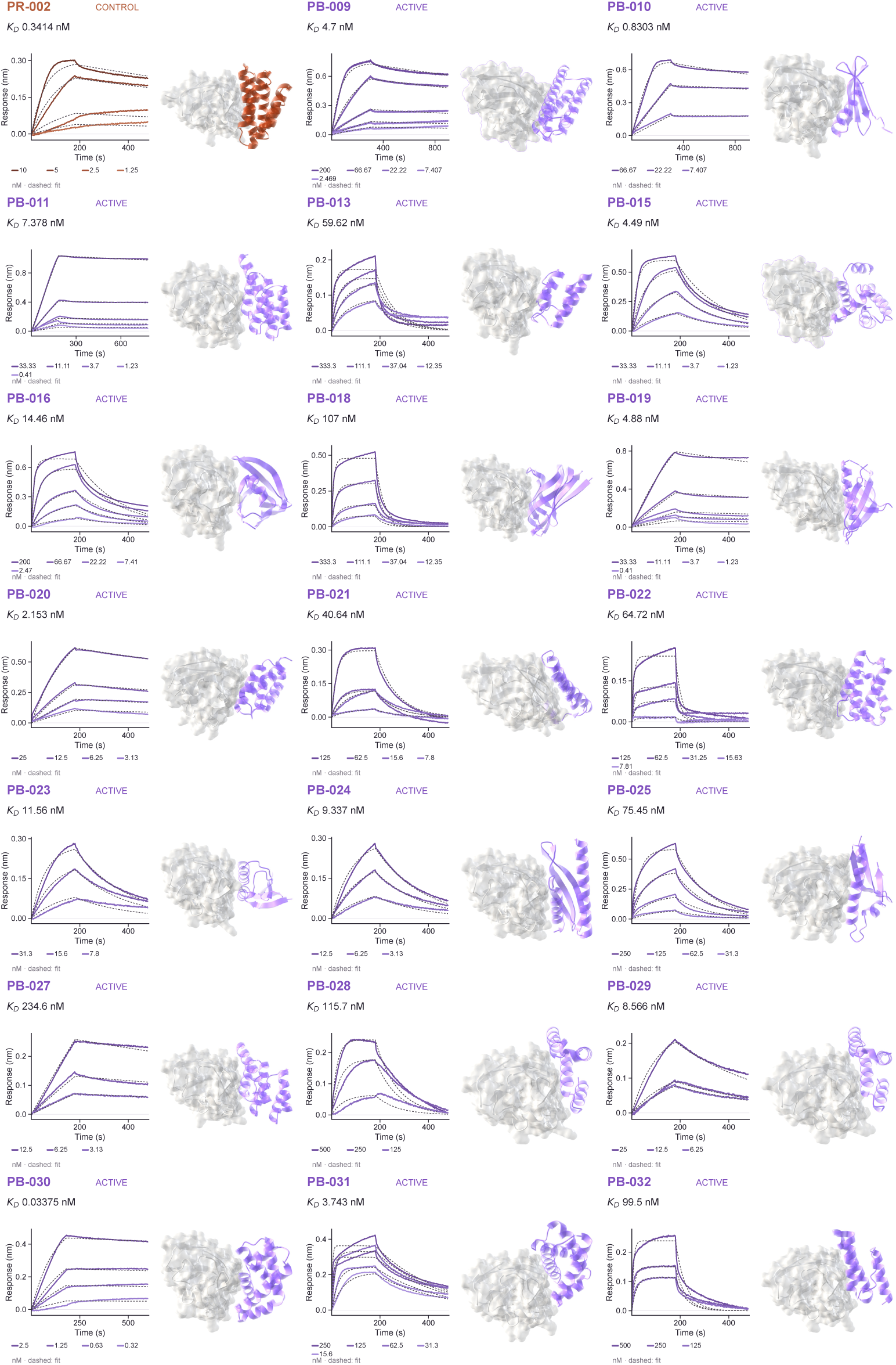
PD-L1. Predicted complexes and BLI for 20 active designs. PR-002 is included as the reference binder.

**Supplementary Fig. S8.**
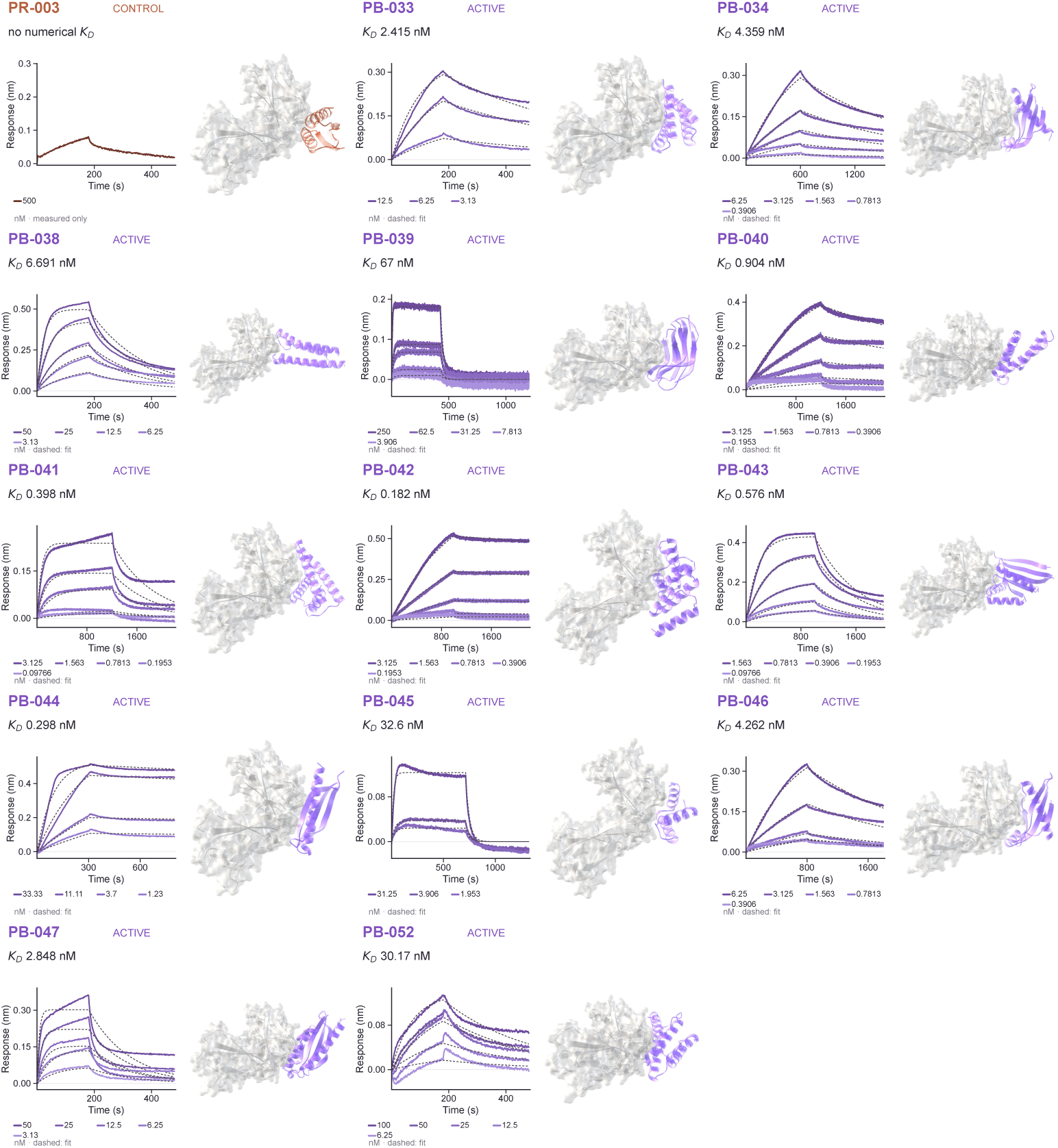
IL-7RA. Predicted complexes and BLI for 13 active designs. PR-003 is included as the reference binder.

**Supplementary Fig. S9.**
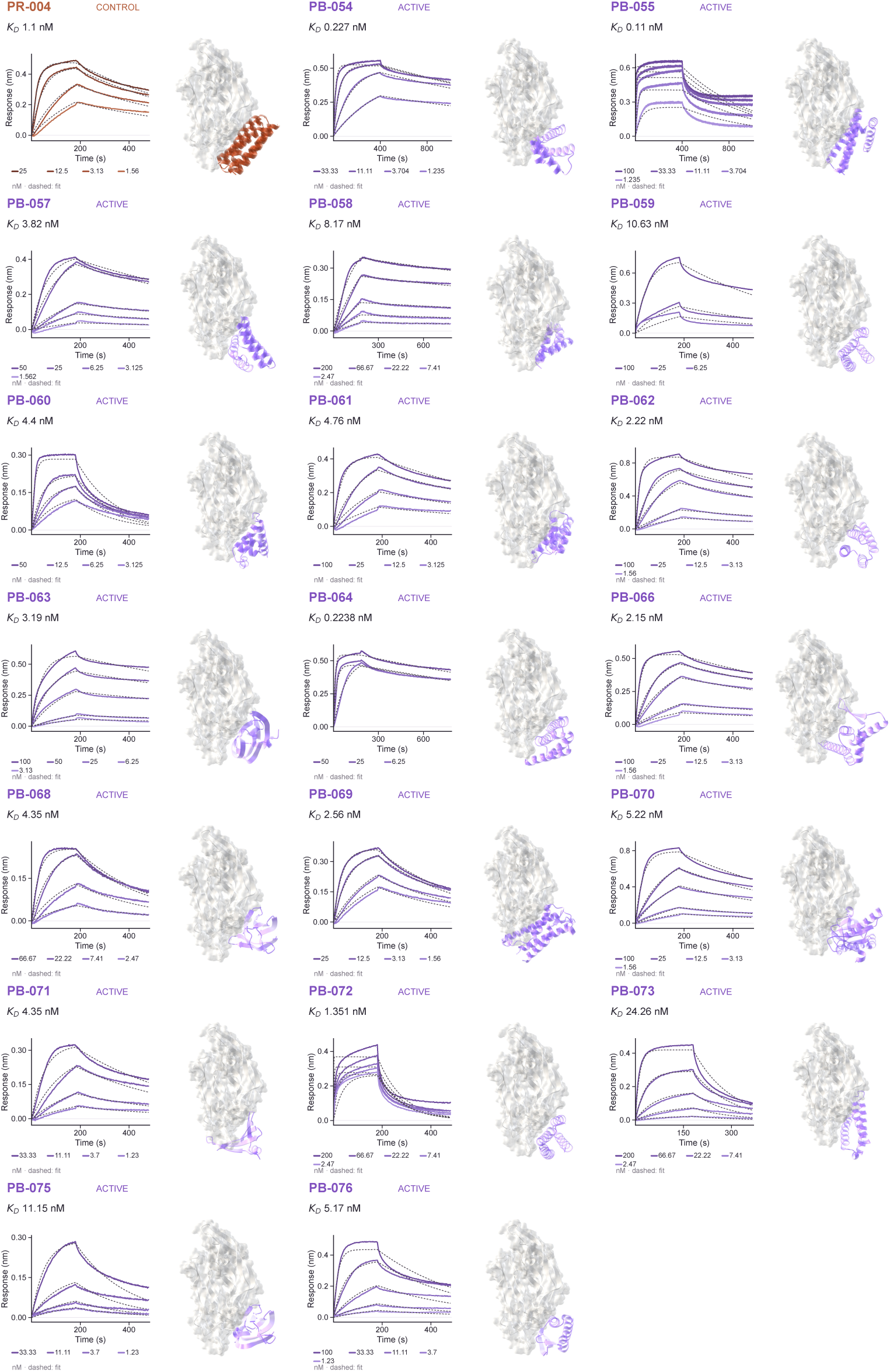
VEGF-A. Predicted complexes and BLI for 19 active designs. PR-004 is included as the reference binder.

**Supplementary Fig. S10.**
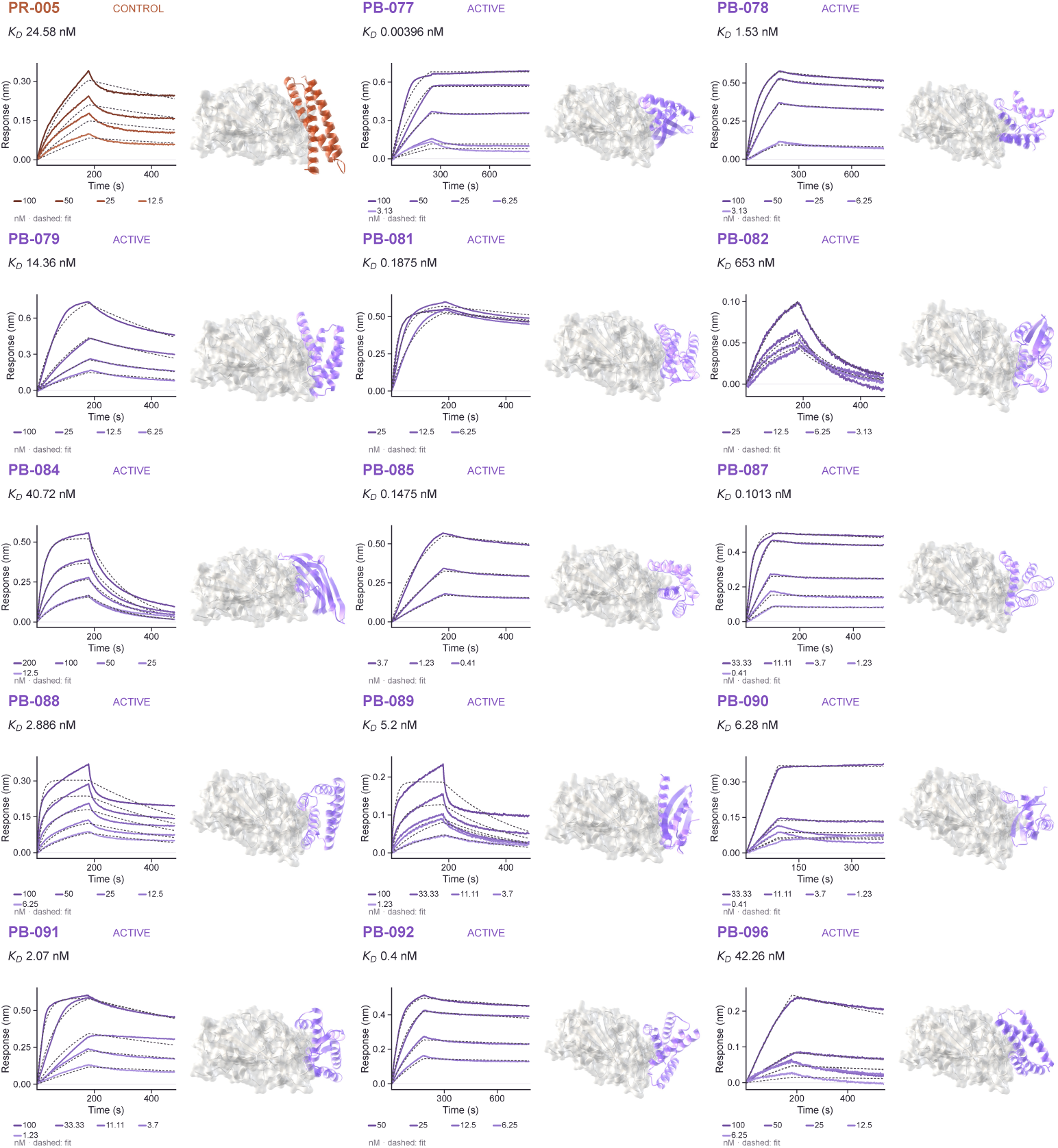
SC2RBD. Predicted complexes and BLI for 14 active designs. PR-005 is included as the reference binder.

**Supplementary Fig. S11.**
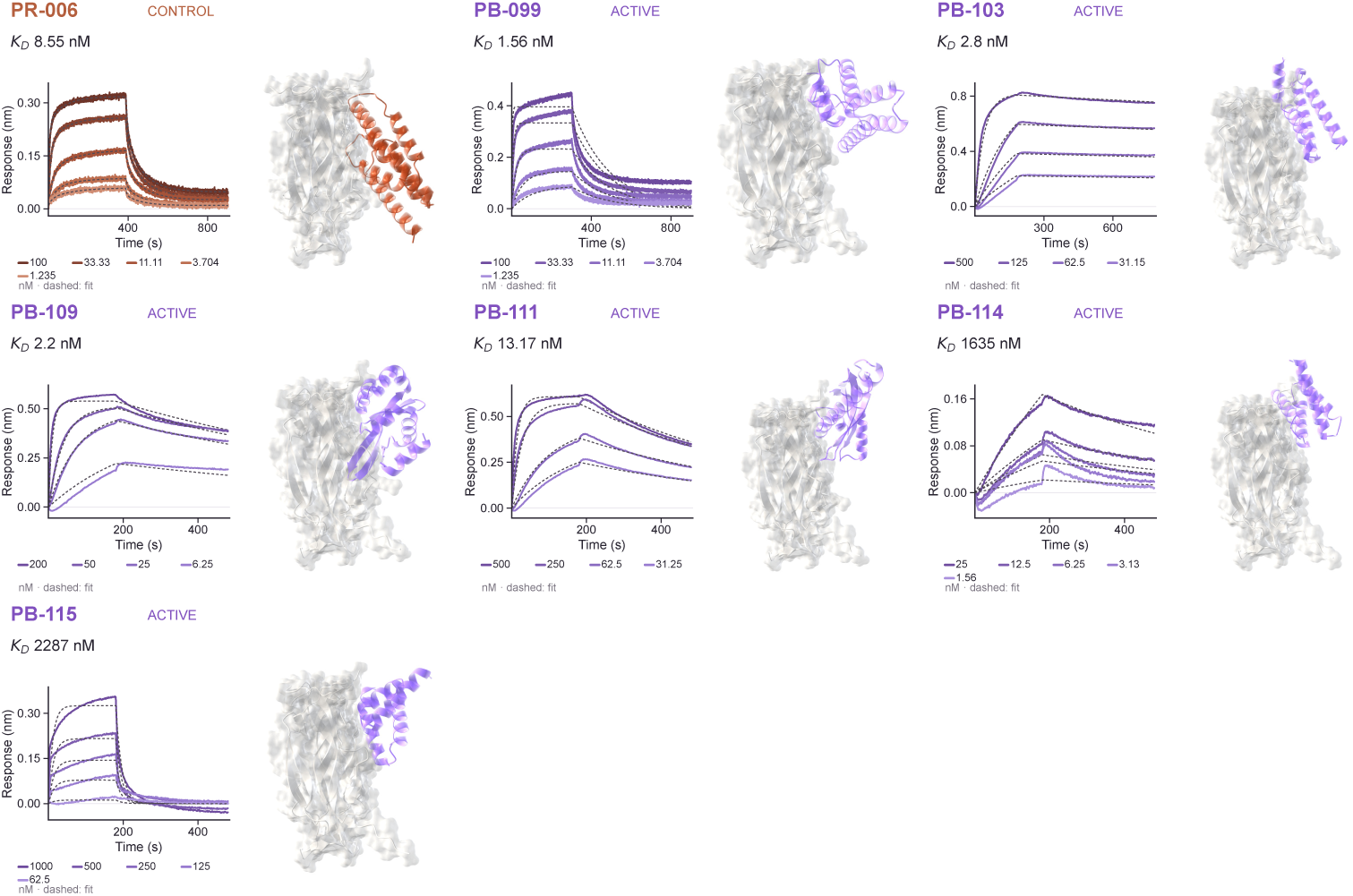
IL-17A. Predicted complexes and BLI for 6 active designs. PR-006 is included as the reference binder.

**Supplementary Fig. S12.**
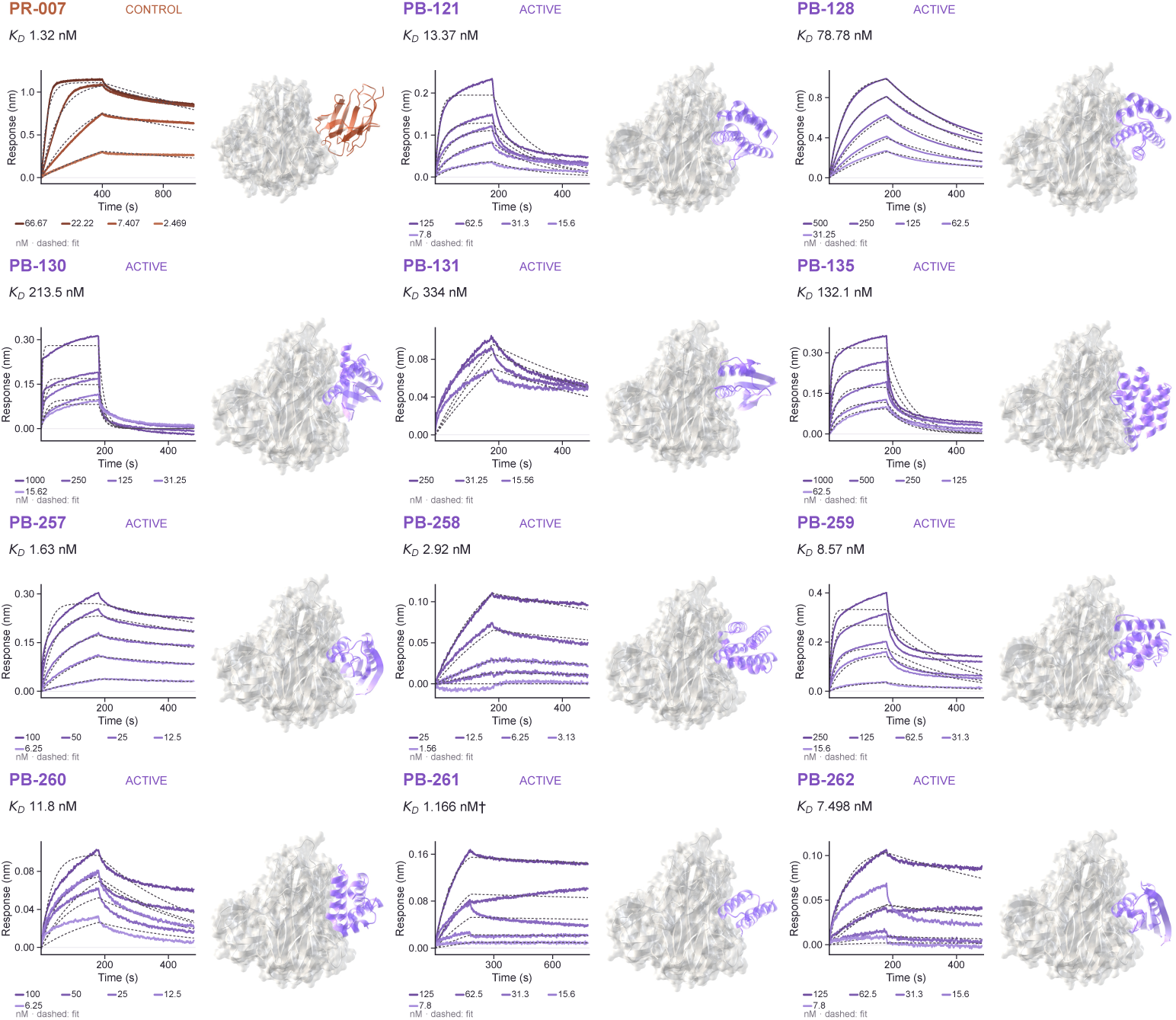
TNF*α de novo*. Predicted complexes and BLI for 11 active designs. PR-007 is included as the reference binder.

**Supplementary Fig. S13.**
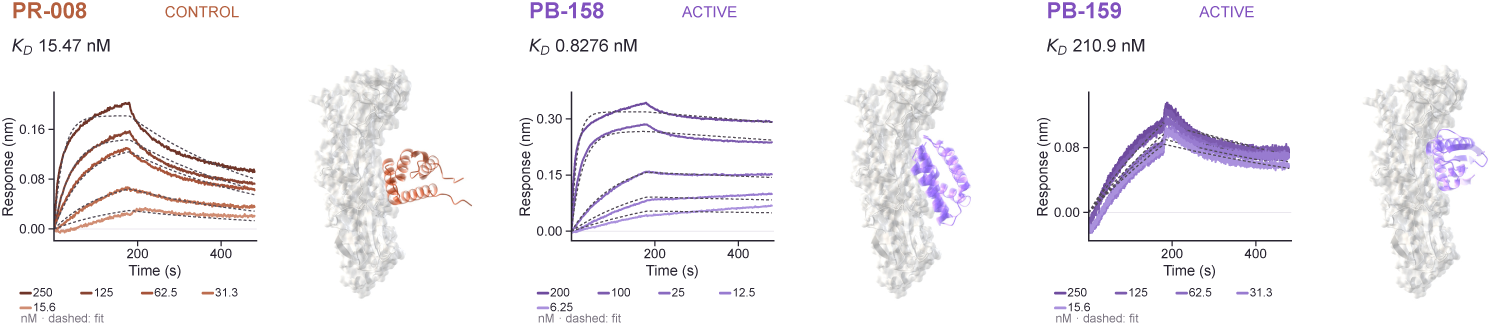
H1. Predicted complexes and BLI for 2 active designs. PR-008 is included as the reference binder.

**Supplementary Fig. S14.**
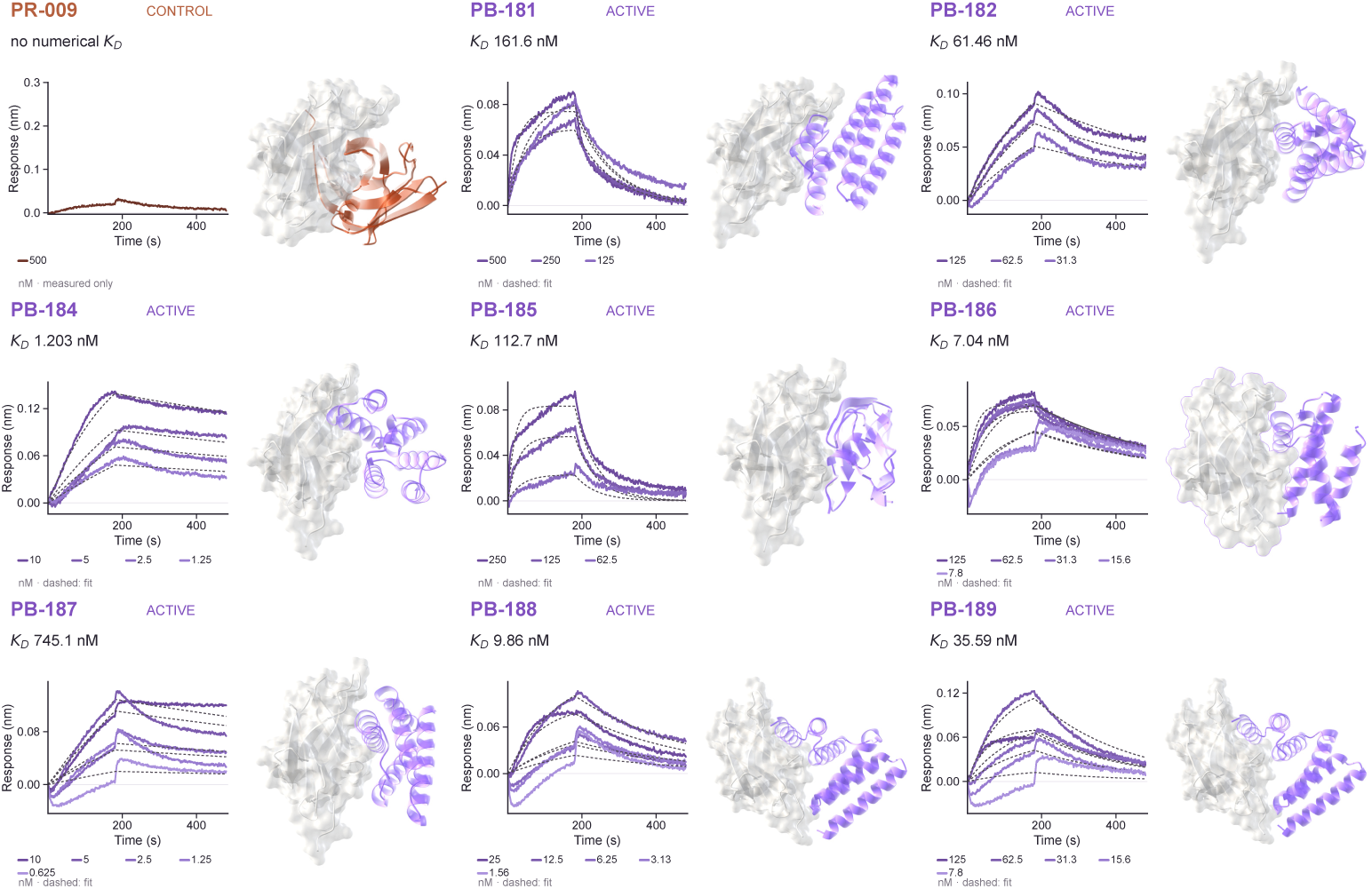
TrkA. Predicted complexes and BLI for 8 active designs. PR-009 is included as the reference binder.

**Supplementary Fig. S15.**
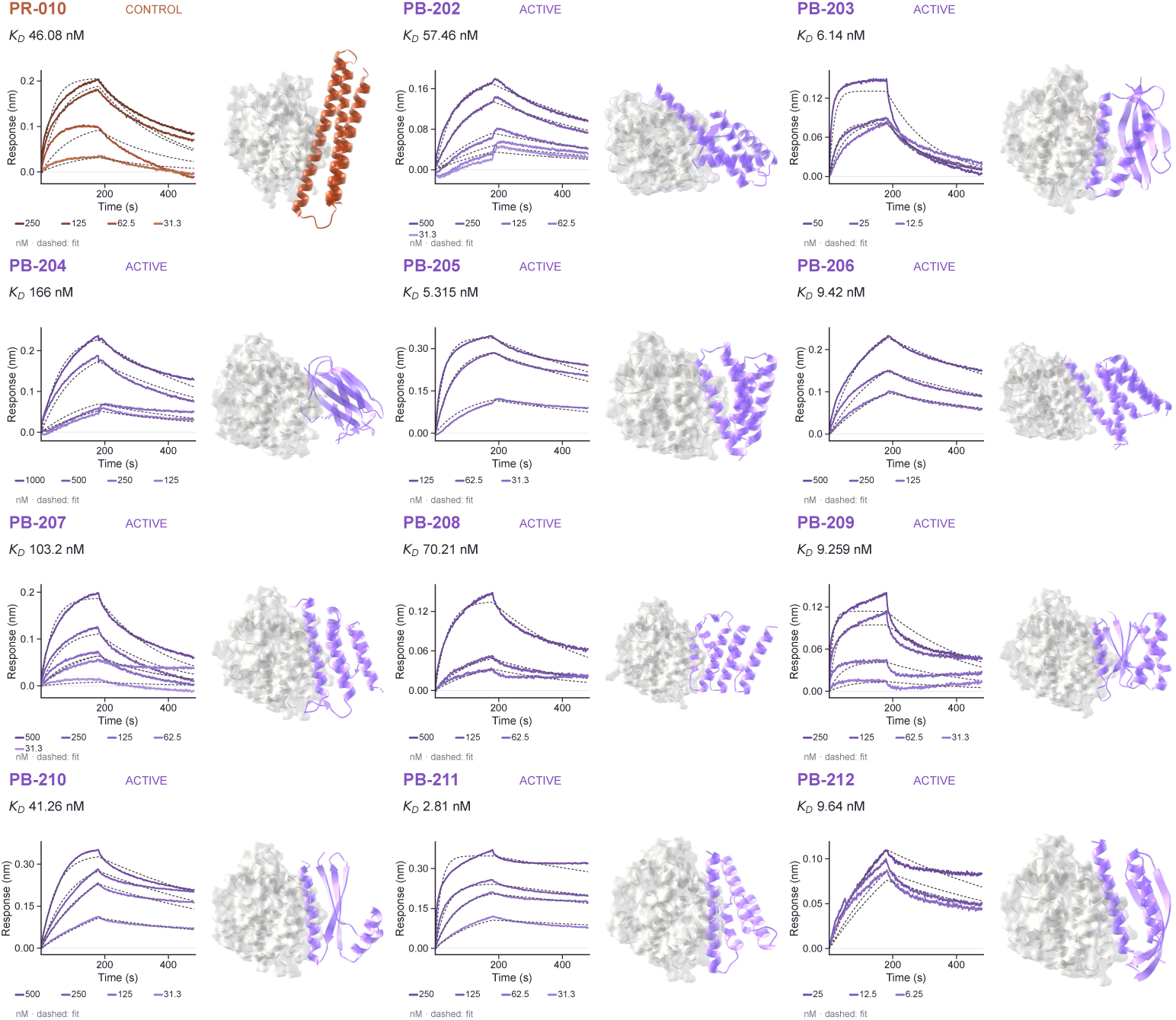
BHRF1. Predicted complexes and BLI for 11 active designs. PR-010 is included as the reference binder.

**Supplementary Fig. S16.**
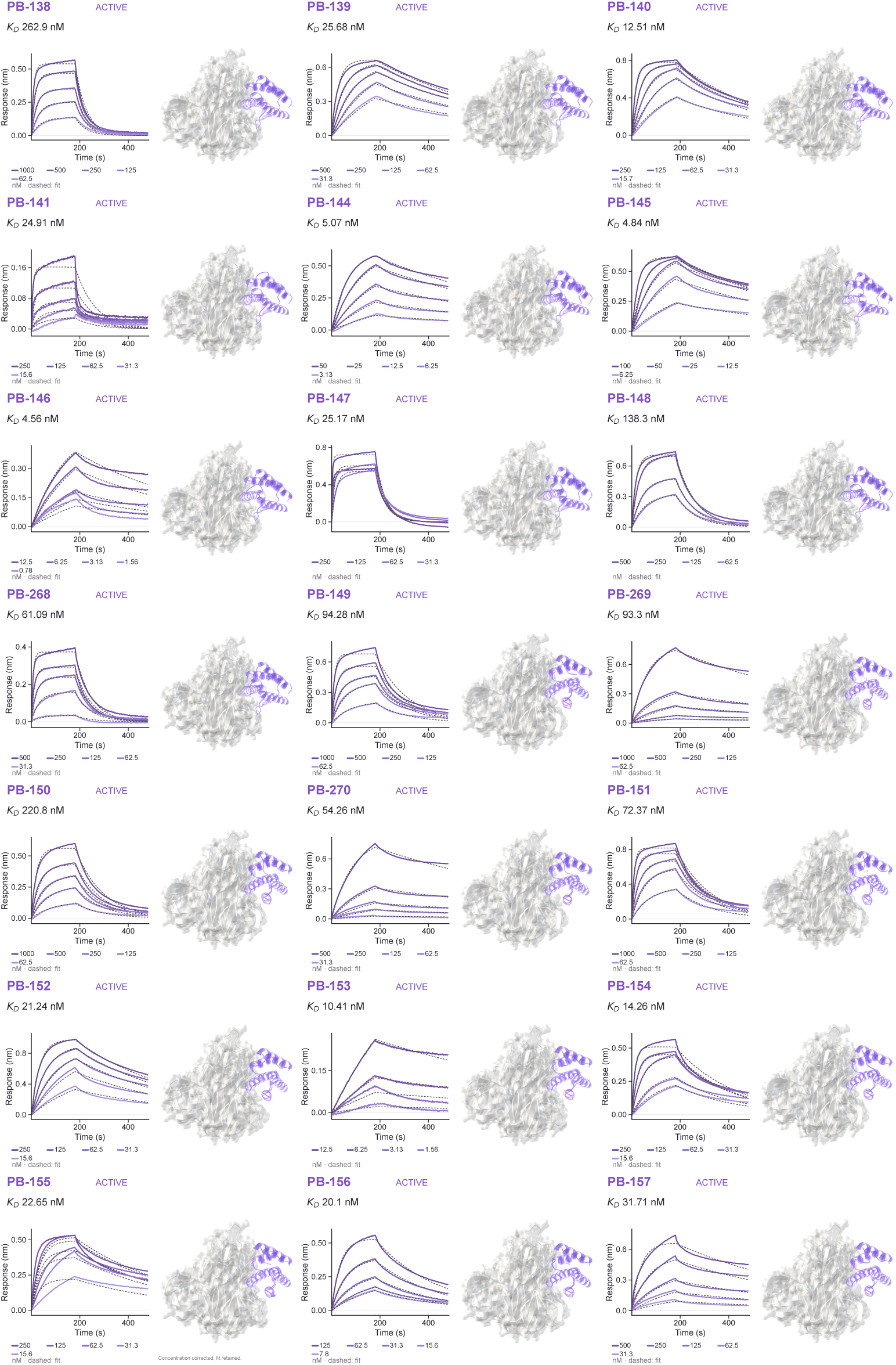
TNF*α* affinity maturation. Predicted complexes and BLI for 21 active affinity-matured variants. The variants were generated from minibinders PB-121 and PB-128.

**Supplementary Table S5.**
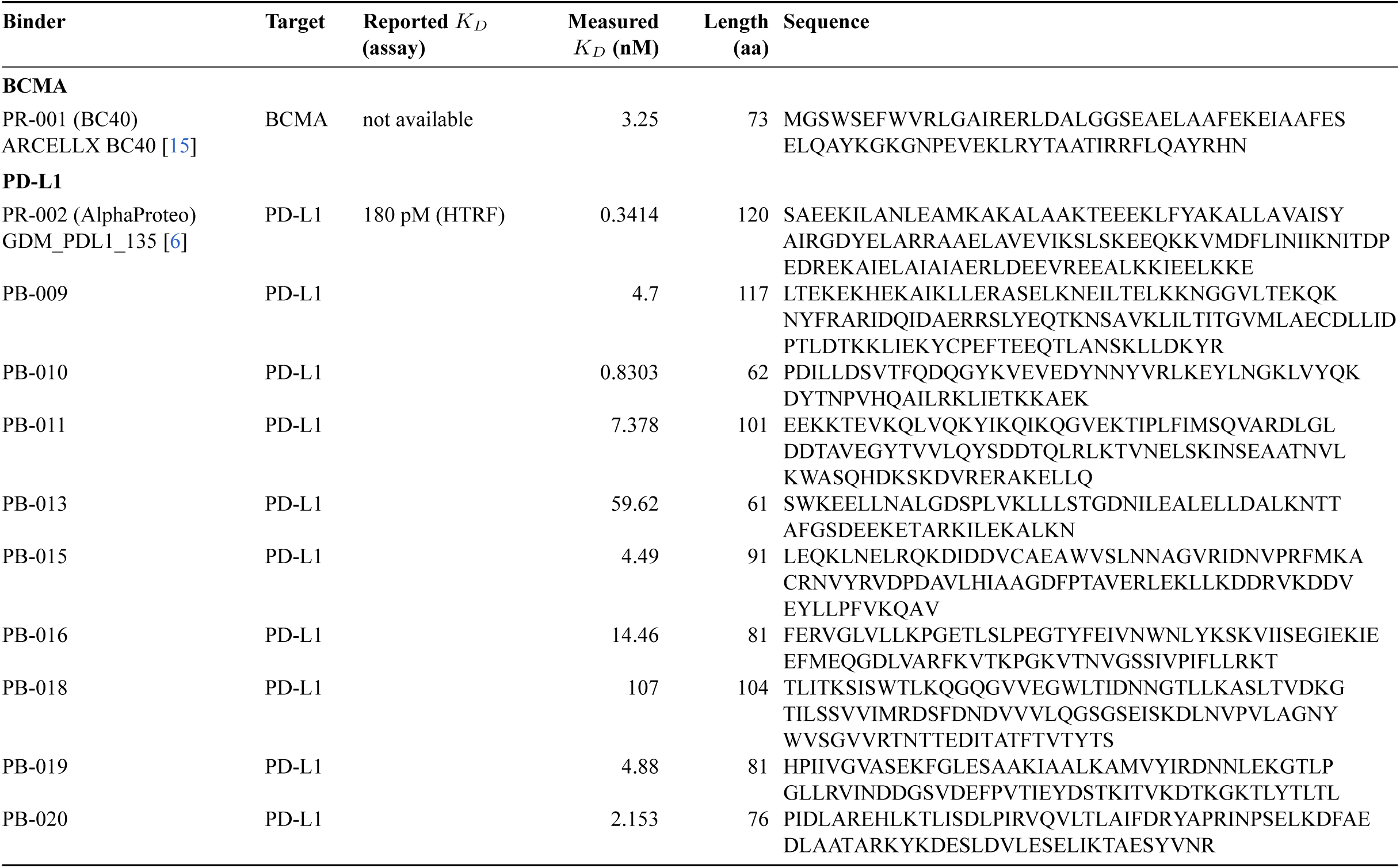
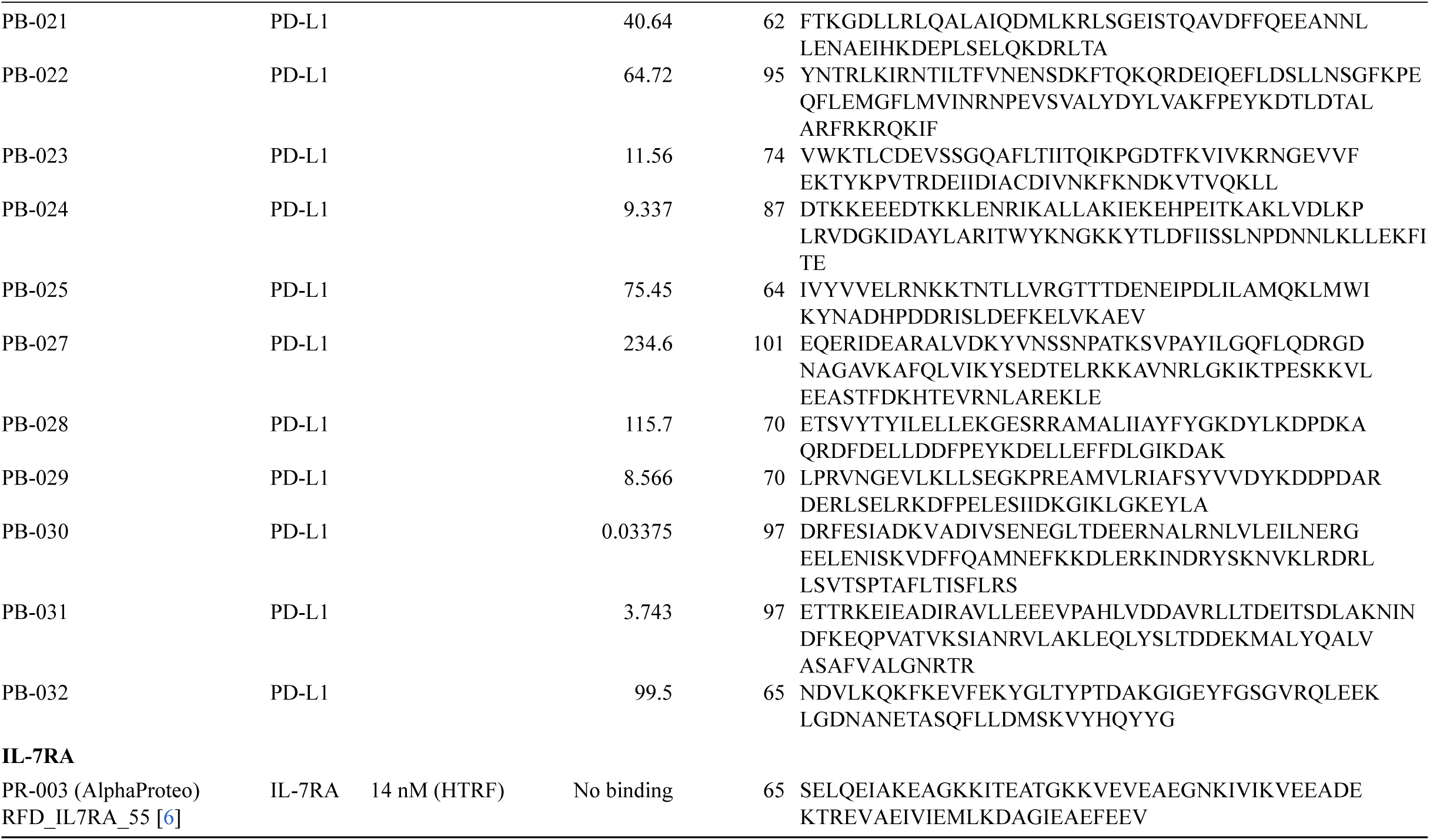
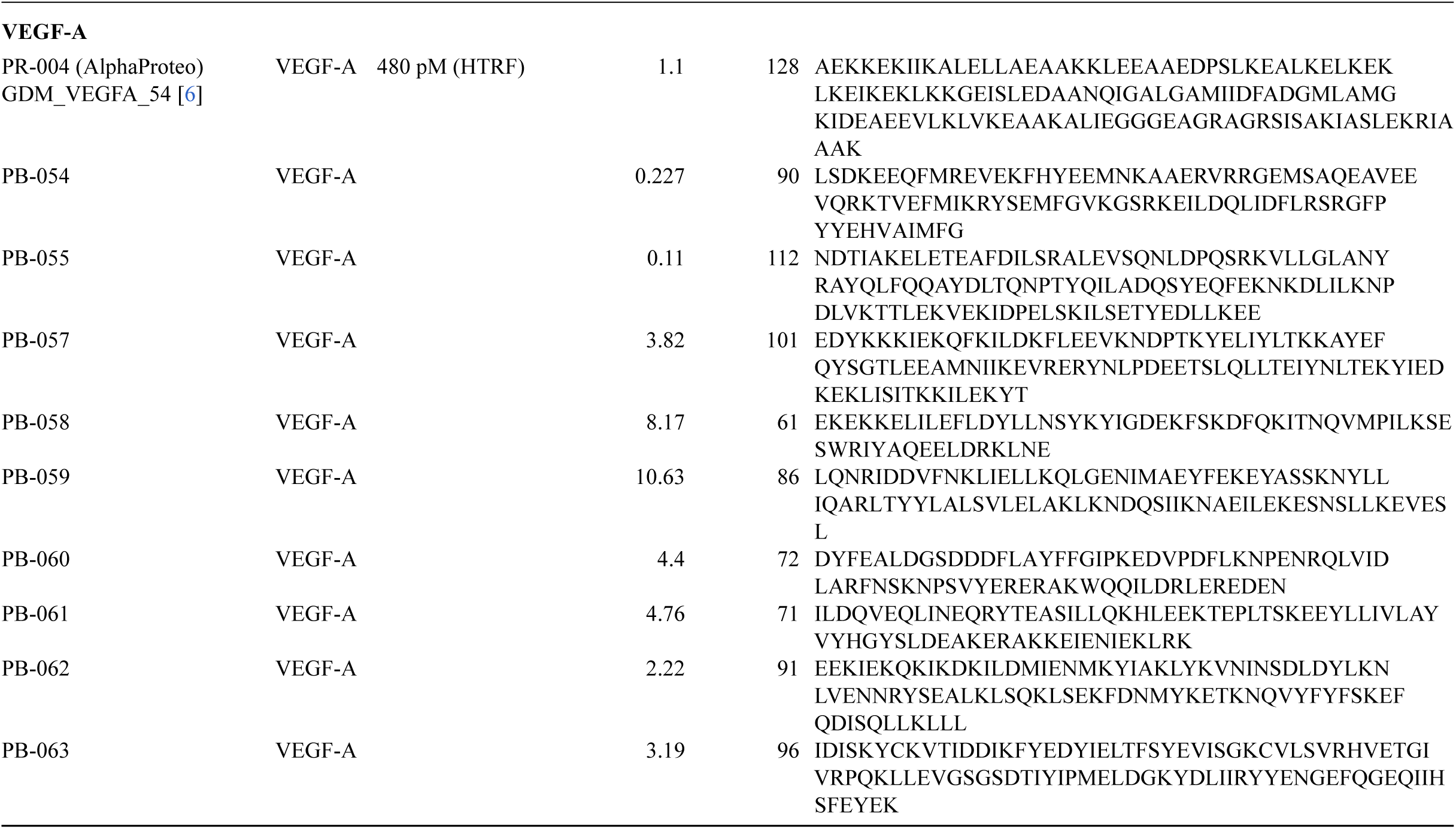
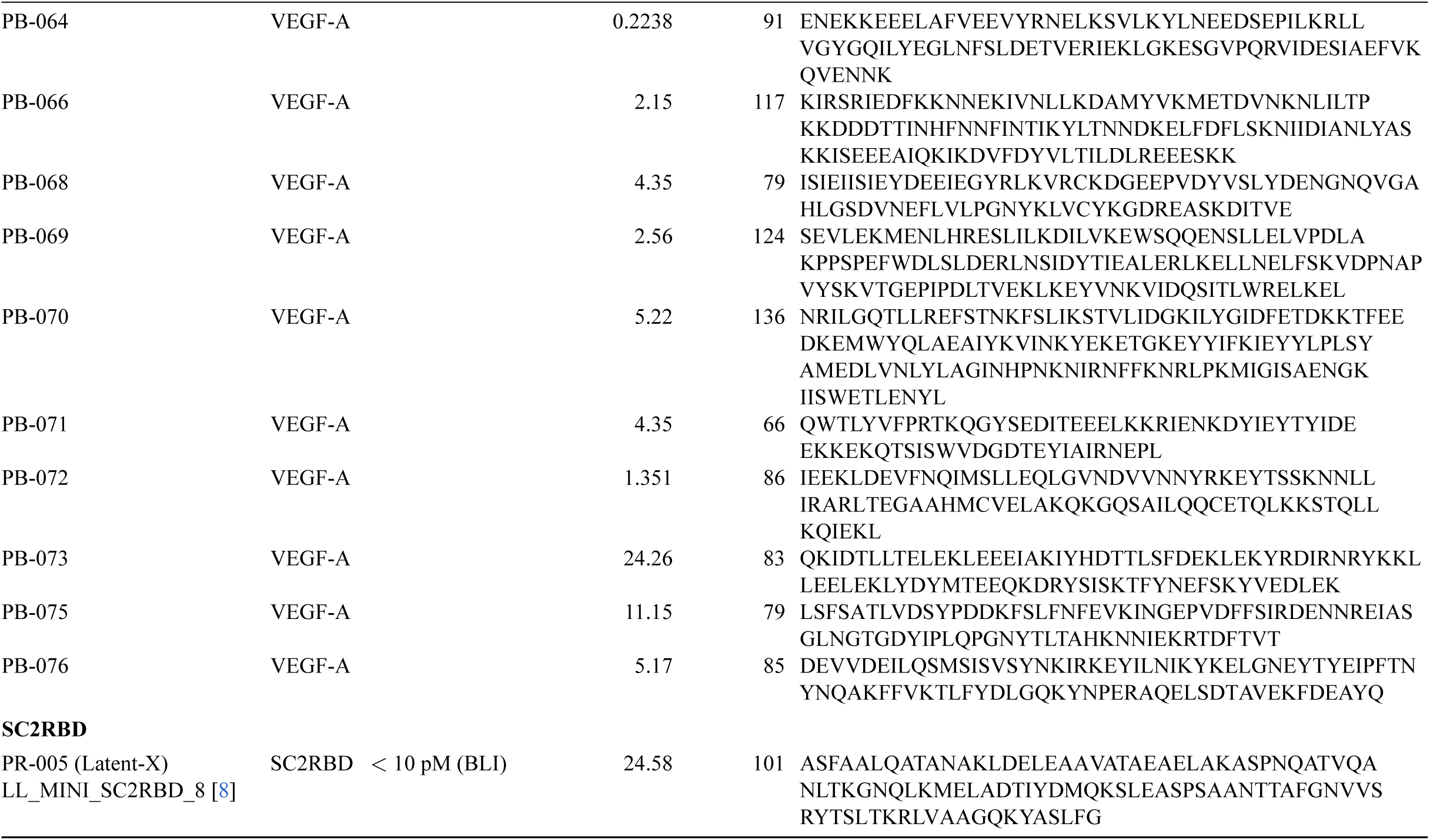
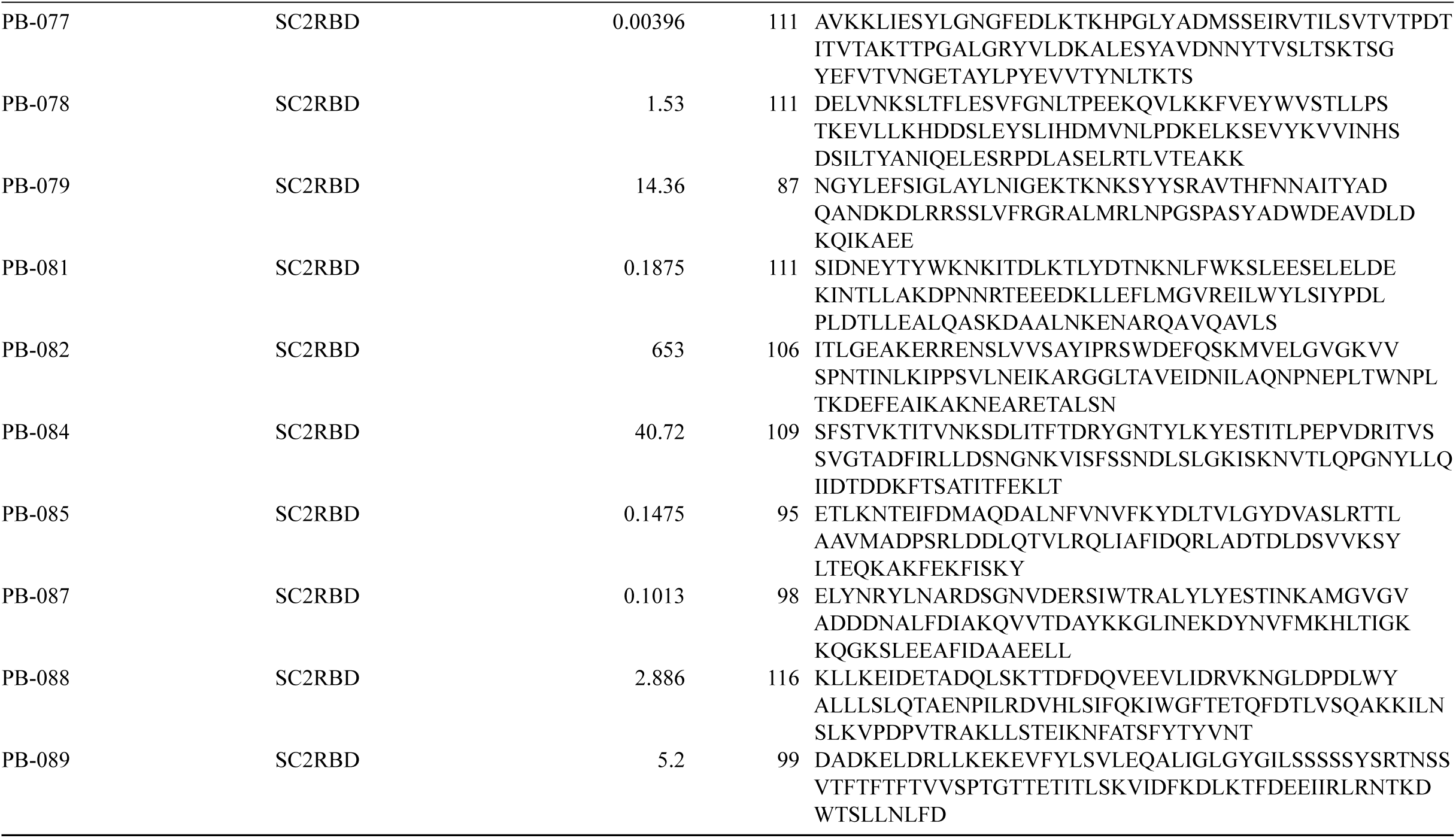
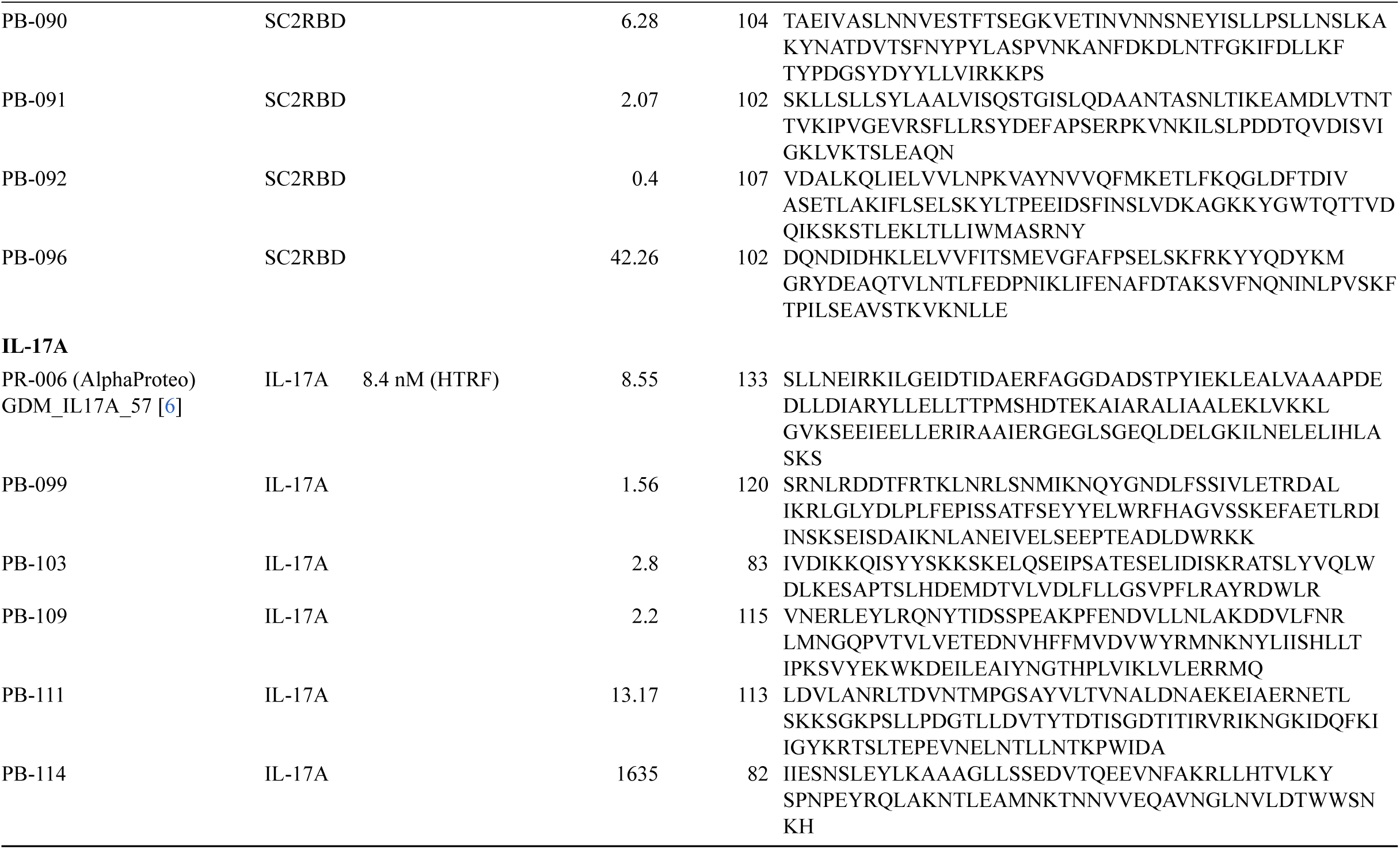
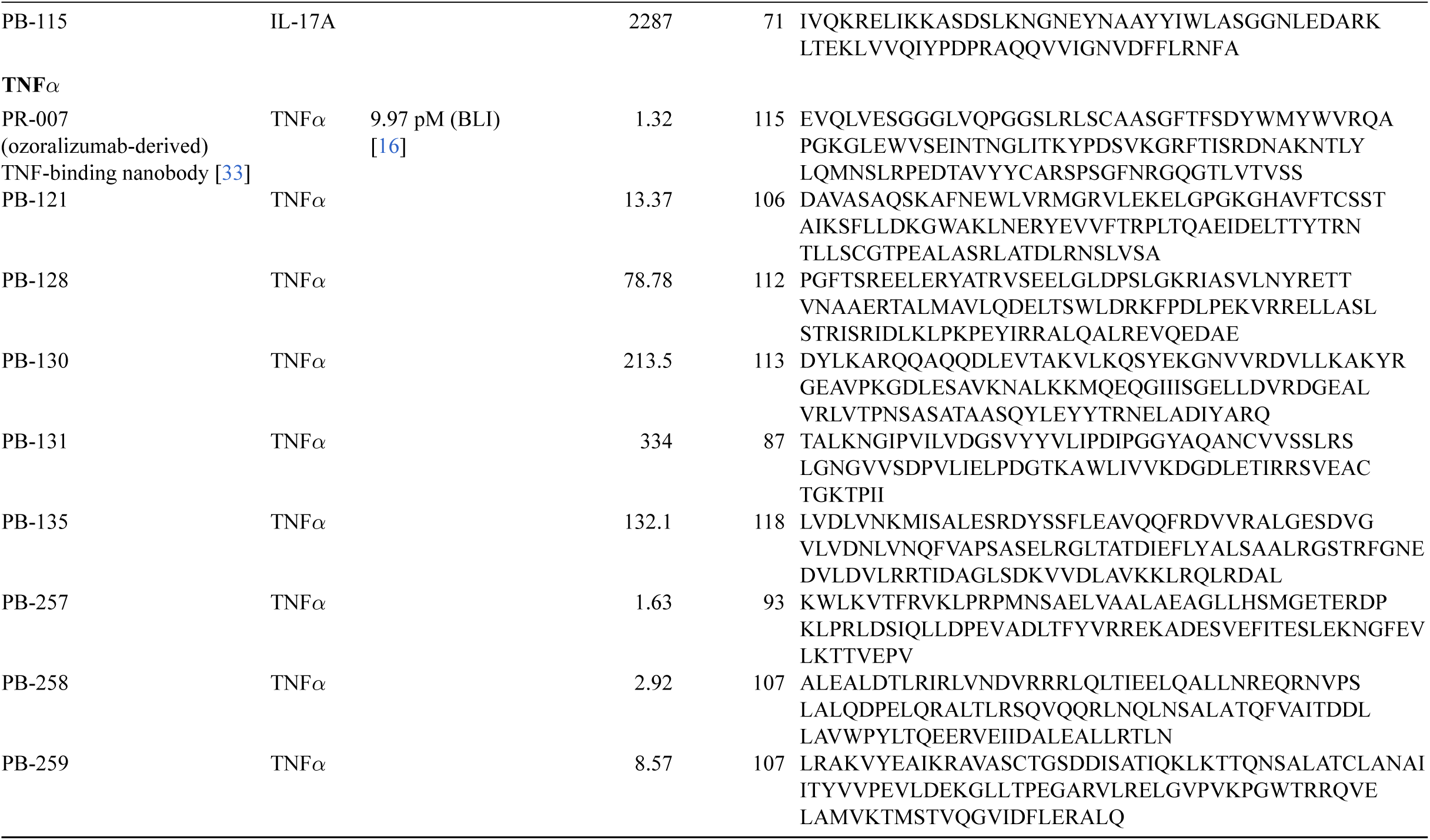
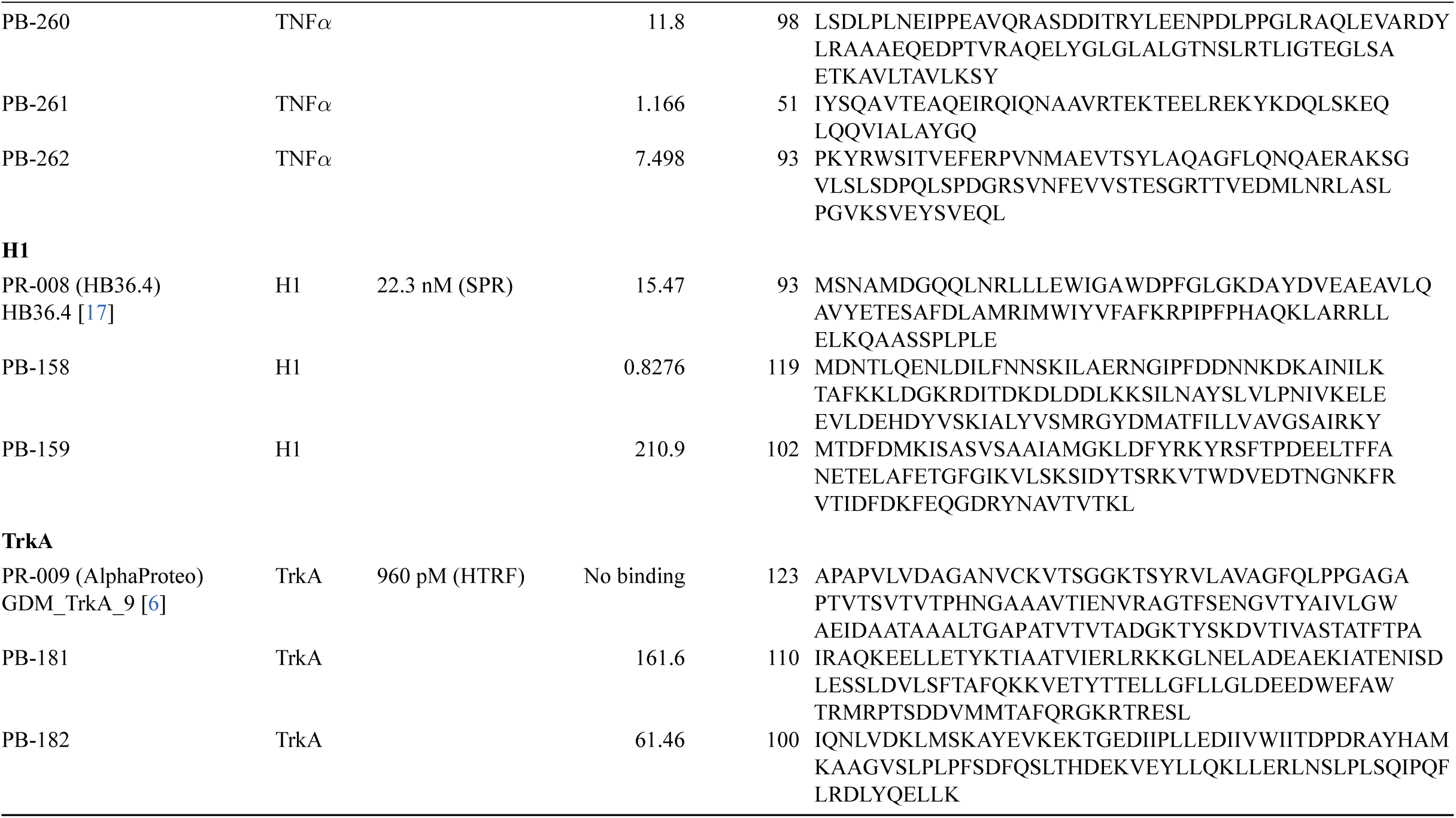
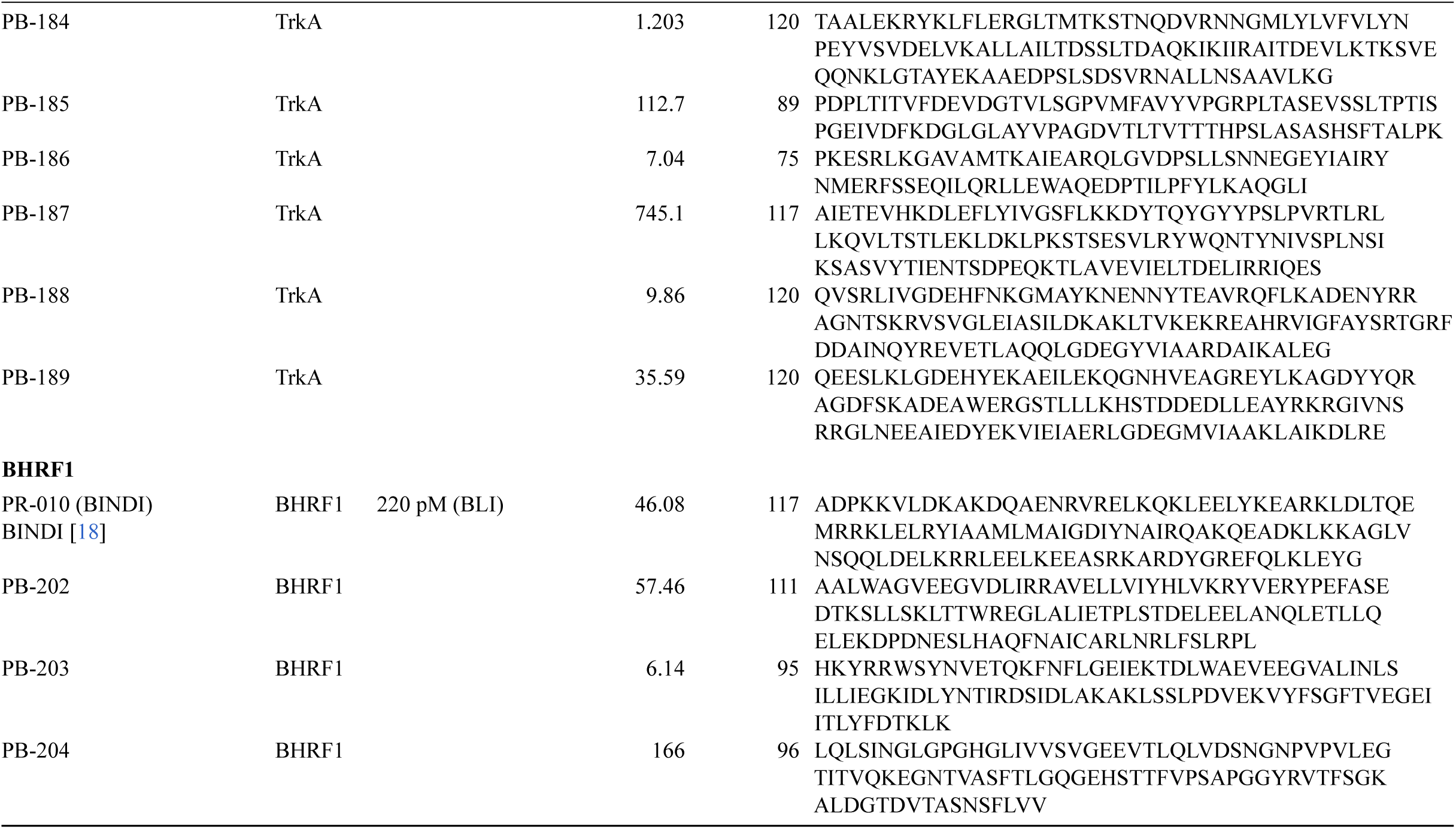
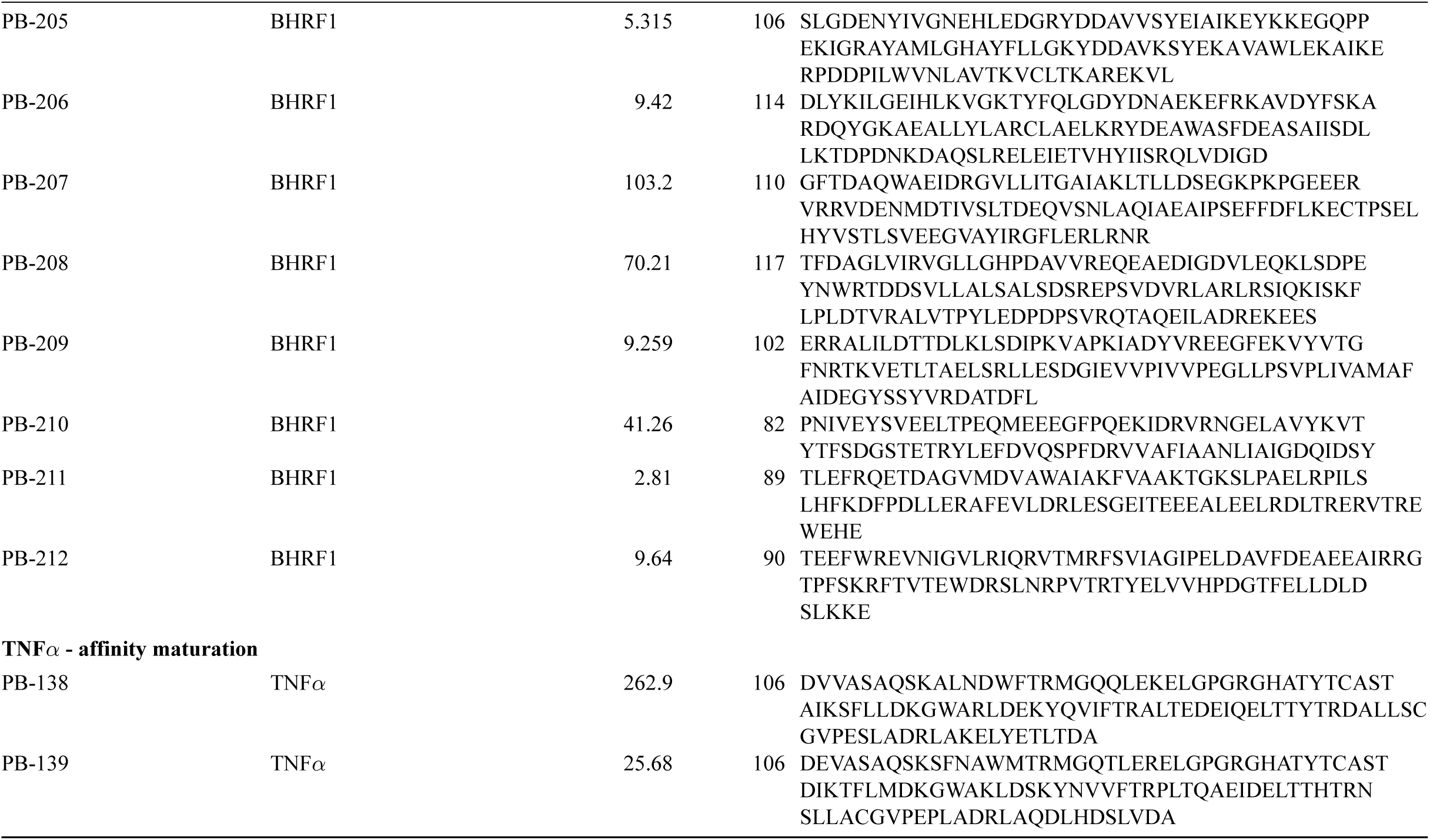
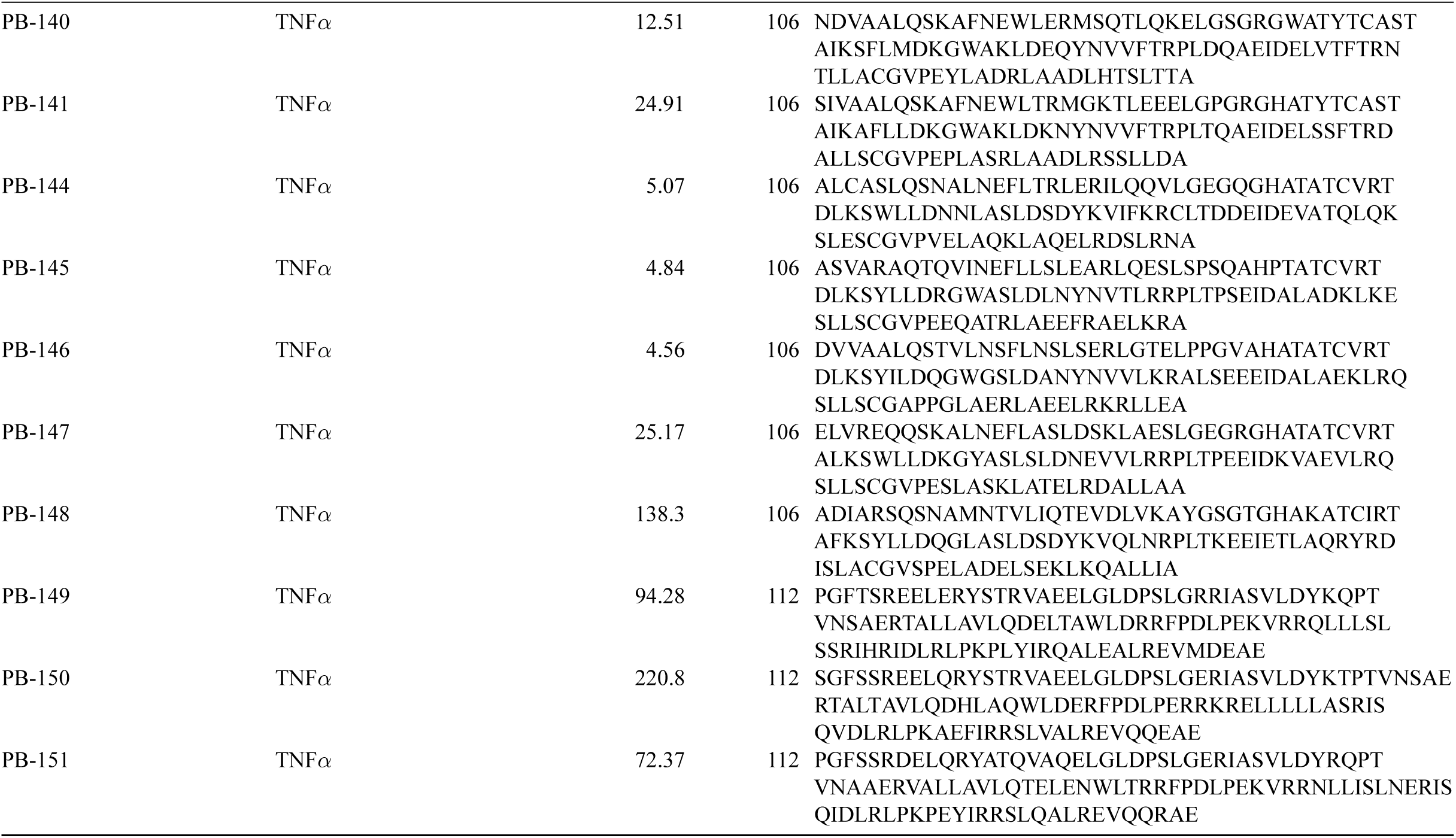
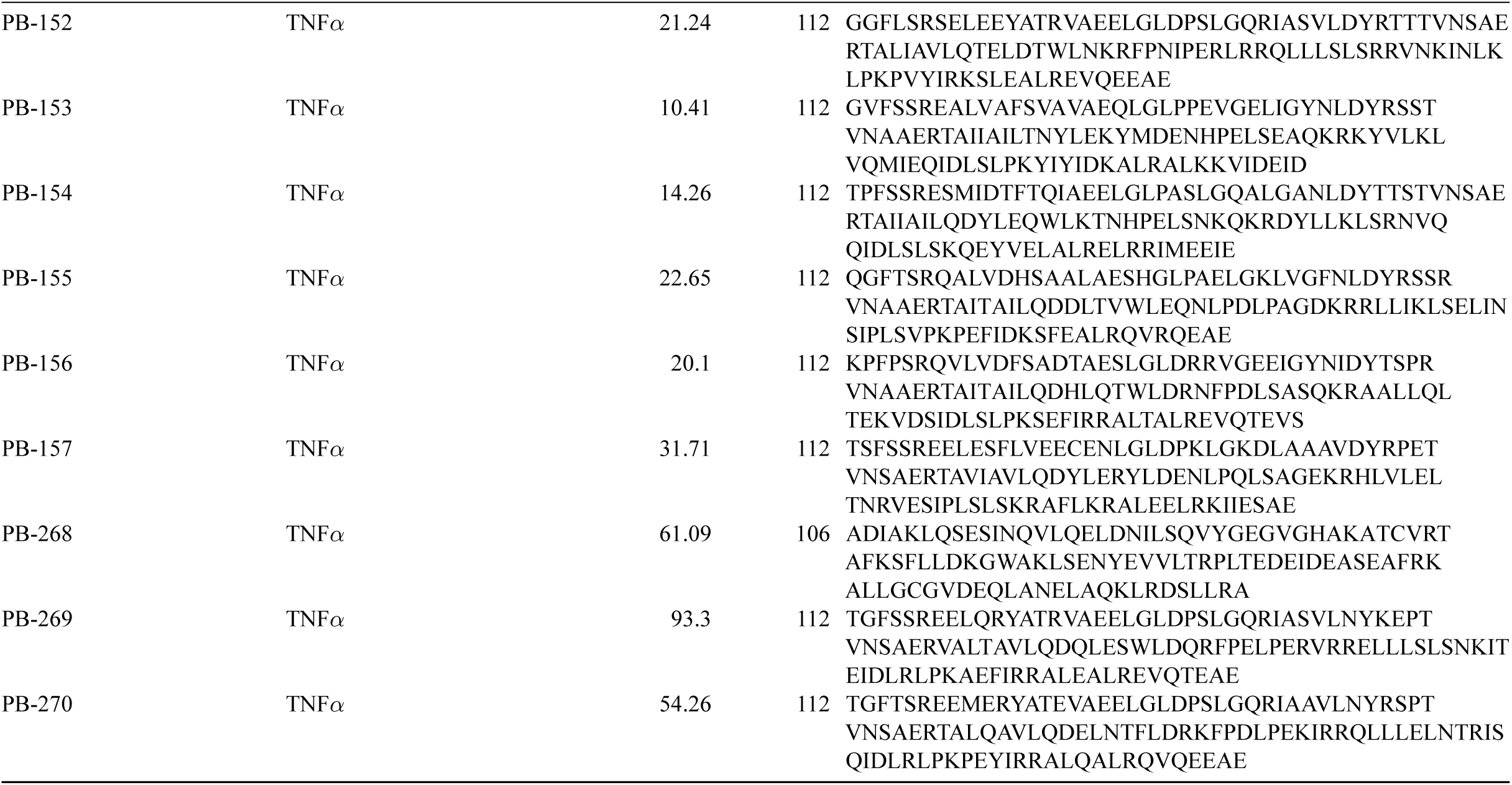
Sequences and affinities of active minibinders and reference binders. Sequences of the nanobodies, BCMA minibinders, and IL-7RA minibinders designed in this work are not disclosed at this stage.

## Footnotes

1 Lévin™ Harness is an AI agent developed by Levinthal Biotechnology specifically for protein design. It connects molecular structure analysis, computational tools and research workflows to support candidate analysis and design iteration; see levinthal.design and the GitHub repository levinthal/levin-harness.

